# Temporal dynamics of gene expression during metamorphosis in two distant *Drosophila* species

**DOI:** 10.1101/2024.06.17.599350

**Authors:** AM Ozerova, DA Kulikova, MB Evgen’ev, MS Gelfand

## Abstract

Complete metamorphosis of holometabolous insects is a complex biological process characterized by profound morphological, physiological, and transcriptional changes. To reveal the temporal dynamics of gene expression during this critical developmental transition, we conducted a detailed analysis of the developmental transcriptomes of two Drosophila species, *Drosophila melanogaster* and *Drosophila virilis*. We confirmed partial recapitulation of the embryonic transcriptional program in pupae, but unlike the traditional hourglass model, suggesting maximal conservation at mid-embryonic stages, at different stages of pupae we observed a more complicated pattern of alternating low and high diversity, resembling an inverted hourglass, or “spindle”. This underscores the complexity of developmental processes during complete metamorphosis. Notably, recently formed genes (specific to insects) exhibit pronounced expression peaks during mid-pupal development, indicating their potential role in developmental transitions.

**Significance Statement:** This study describes the transcriptomic dynamics of complete metamorphosis in holometabolous insects. By analyzing developmental transcriptomes of *Drosophila melanogaster* and *Drosophila virilis*, we characterize the spindle pattern of gene expression during pupal stages, contrasting with the classical hourglass model of embryogenesis. This alternating pattern of low and high transcriptional diversity highlights the unique complexity of the metamorphose processes.

## Introduction

Insects undergoing complete metamorphosis (holometabolous insects) represent around 60% of all animals (Mora et al. 2011), suggesting evolutionary success. Their intricate life cycle has been proposed to be a key contributing factor to this accomplishment. One of the adaptive benefits is the occupation of multiple niches which allows for switching between alternative resources (Ten Brink, de Roos, and Dieckmann 2019)

Upon hatching, larvae reach a considerable size within a short timeframe (Texada, Koyama, and Rewitz 2020). Through successive molts, they experience rapid growth, eventually culminating in the stage of transformation, the pupa. Rather than resembling miniature adults, as seen in nymphs of hemimetabolous species, holometabolous insects undergo a radical change that endows them with the appearance of an entirely distinct organism. This process involves rebuilding of virtually all larval organs with those required for adult life.

Several mechanisms underlie the terminal differentiation of tissues and organ formation. One such mechanism is dedifferentiation, followed by redifferentiation into adult tissues, a distinguished example being restructuring of the insect longitudinal gut muscles. In the course of this process syncytial larval muscle cells dissociate into mononuclear cells before redifferentiating to form the adult muscles (Schultheis and Frasch 2023).

Unlike the musculature, the intestinal epithelium undergoes a complete renewal process during metamorphosis, representing another mechanism for rebuilding of the organs. Adult midgut precursors are maintained on the basal side of the larval midgut throughout the larval development and undergo rapid divisions only after formation of the puparium (Micchelli et al. 2011).

Another example of a complex process during metamorphosis is development of the adult brain. While larval neurons are retained, the peripheral and central nervous systems are significantly expanded through activation of numerous neurons that remained in the state of developmental arrest throughout larval growth (Truman 1990).

The intricate processes inherent in pupal development share a level of complexity comparable to embryogenesis, demanding precise orchestration of genes involved in the development of emerging adult organs and systems. Transcriptomic analyses across various insect orders revealed reactivation of the embryonic gene expression program at pupal stages (Ozerova and Gelfand 2022). While the developmental hourglass model has been traditionally applied in the context of embryogenesis, its potential extension to the metamorphic phase has not been considered so far. It is somewhat parallel to phenomena observed in plants, where the hourglass pattern emerges in transcriptomes at several crucial stages, such as germination and floral transition (Drost et al. 2017).

The developmental hourglass model postulates that mid-embryogenesis stages exhibit the highest level of conservation, while early and late stages are more diverged (Duboule 1994). These mid-embryogenesis stages, often referred to as the phylotypic period, play a crucial role in establishing the body plan of the future organism. The term “phylotypic” originates from the period’s association with the maximum similarity observed among species within each animal phylum (Richardson 1995). This conservation is believed to stem from the network of interactions between genes and developmental processes, where any evolutionary modifications could prove to be deleterious due to potential side effects (Raff 2012). On average, genes that are expressed during the phylotypic stage tend to have a higher predicted age compared to genes expressed earlier and later in the course of development (as measured by the transcriptome age index in (Domazet-Lošo and Tautz 2010)). Another explanation is that the phylotypic period is essential for the precise coordination between growth and patterning (Duboule 1994).

The hourglass pattern in embryogenesis was observed on morphological and molecular levels across various groups of organisms, including mammals (based on the expression of human and mouse orthologous genes) (Hazkani-Covo, Wool, and Graur 2005), *Drosophila* spp. *(Kalinka et al*. *2010), Caenorhabditis* spp. *(Levin et al*. *2012)*. Despite this evidence, the hourglass hypothesis remains a subject of controversy, with some studies challenging both the hourglass model and the existence of the phylotypic stage (Irie 2017). A recent study, analyzing gastrulation in rabbits and mice at the single-cell resolution, demonstrated clear differences in mid-phases between the two species (Mayshar et al. 2023). In this case, earlier and later developmental phases were more similar in terms of cell type composition as revealed by the single-cell transcriptome analysis. This observation supports a *spindle model*, which is a pattern characterized by low diversity at early stages, increasing in mid-development, and decreasing again at the later stages, i.e. an apparent inversion of the known hourglass pattern.

Here, we explore the developmental transcriptomes of two distant cosmopolitan species *Drosophila melanogaster* and *Drosophila virilis*, aiming to characterize the dynamics and conservation of gene expression throughout metamorphosis. The divergence of 63 million years between the species (Tamura, Subramanian, and Kumar 2004) enables quantitative comparisons across orthologous genes. While transcriptomes have been extensively studied and compared in embryogenesis (Kalinka et al. 2010), gene expression divergence during metamorphosis remains largely unexplored, and it is not clear what model of developmental divergence, if any, applies to this process.

## Results

### Reproducibility and preprocessing of RNA sequencing data

To investigate the dynamics of gene expression during development, we performed RNA sequencing (RNA-seq) experiments on the developmental time course of *Drosophila melanogaster* and *Drosophila virilis*. A comprehensive analysis was conducted across ten distinct developmental stages in each species, including the embryonic phylotypic stage (Kalinka et al. 2010), larvae, seven stages spanning pre-pupae and pupae development, and the adult (imago) stage. Each developmental stage was sampled in three biological replicates to capture the inherent variability in gene expression.

Whole body samples rather than specific tissues were analyzed to maintain consistent sampling conditions across all developmental stages, as specific tissues do not persist throughout each stage. This approach captured overall gene expression dynamics reflecting the developmental program as a whole. While tissue-specific expression differences could provide additional insights, a whole-organism approach allowed us to maintain comparability across stages, facilitating the identification of broader developmental patterns. On the other hand, focusing on whole bodies may overshadow more subtle differences in expression that could be observed at the tissue or cell type level.

RNA sequencing generated approximately 2–2.5 million reads per sample for both *D. melanogaster* and *D. virilis*. After quality filtering, around 1 million reads remained, with a similar number successfully mapping to the reference genomes. However, uniquely mapped reads differed between species, with *D. melanogaster* having approximately 0.3 million uniquely mapped reads and *D. virilis*, around 0.6 million reads. A detailed summary of read processing and mapping statistics is provided in Supplementary Figure 1.

To evaluate the reproducibility of the RNA-seq data, the concordance among the biological replicates for each sample was assessed. The replicates demonstrated a high degree of similarity, indicating robust and reproducible measurements across biological replicates (the Spearman correlation coefficient being equal to 0.97 on average, Supplementary Figure 2). The Spearman correlation coefficient was higher than 0.84 for all replicate pairs in both species, the lowest example being larval samples of *D. melanogaster*. Consequently, gene expression values were averaged across the replicates, generating a unified gene expression matrix for the downstream analyses.

In the *D. virilis* dataset, one of the replicates from the P3 and P4 pupal stages showed increased variation compared to other two replicates, with the largest divergence observed along the first principal component (Supplementary Fig. 3b). To prevent potential bias in the results, downstream analyses were conducted excluding these outlier replicates. All observed dynamics of expression conservation between species remained consistent when the full dataset was analyzed (see below and Supplementary Fig. 4).

An average of 25,000 transcripts per sample for *D. melanogaster* and 20,000 transcripts per sample for *D. virilis* were identified. To simplify the data and facilitate comparative analysis, the transcripts were aggregated at the gene level, resulting in an average of 12,000 genes for each developmental stage in both species. Grouping of genes into orthologous groups yielded 10,000 one-to-one orthologous groups shared by *D. melanogaster* and *D. virilis*. To ensure robust and informative downstream analyses and focus on genes that exhibited significant expression changes during development, the dataset was refined by filtering out genes with low expression values and low variance across the developmental stages (see *Materials and Methods*). The final dataset comprised 6,200 orthologous groups of genes that demonstrated dynamic expression profiles across the developmental time course in both *D. melanogaster* and *D. virilis*.

### Intra-species analysis of transcription profiles during pupal development in *D. melanogaster* and *D. virilis*

To characterize processes occurring in each species, initial analyses of the transcription profiles were performed for *D. melanogaster* and *D. virilis* separately. During the pupal development phase, gradual modifications in the expression profiles were observed, with the transcriptome at each stage exhibiting higher correlation with neighboring stages rather than more distant ones, as determined by the Spearman correlation coefficient and the Euclidean distance (Fig. 1a and Supplementary Fig. 5).

**Figure 1.**
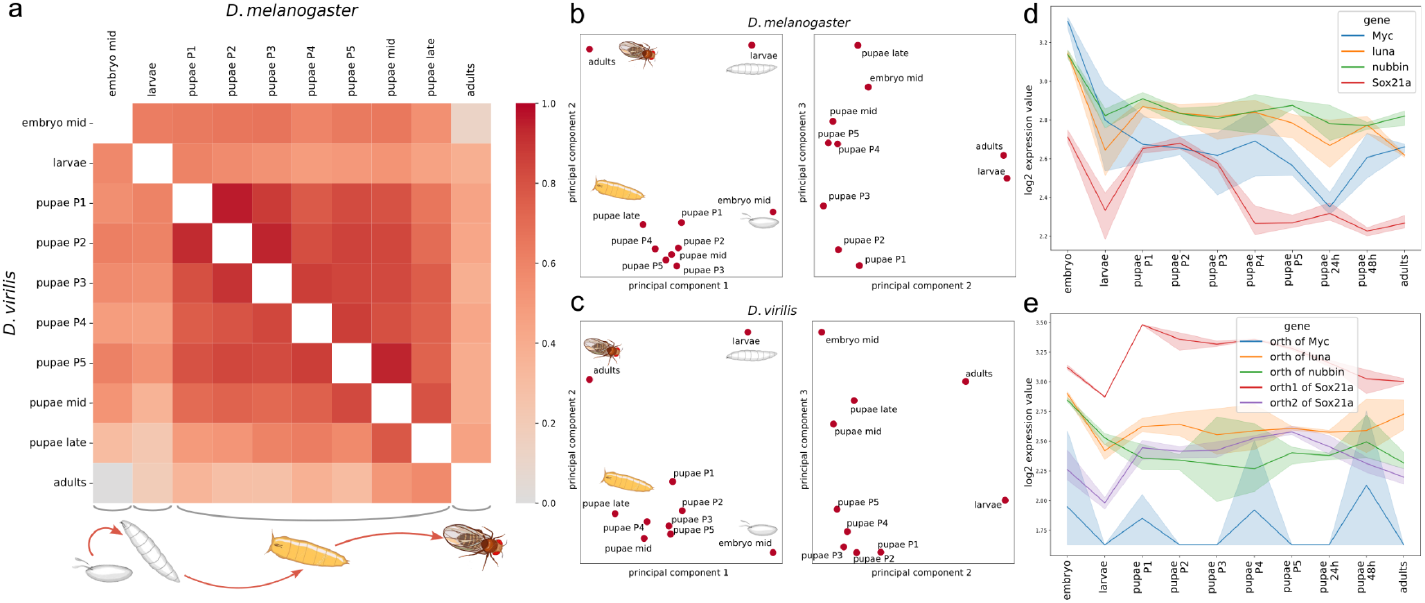
**(a)** Heatmap of the Spearman correlation coefficients for *D. melanogaster* (above the diagonal) and *D. virilis* (below the diagonal) developmental stages for each species separately. Each square represents the correlation coefficient between the respective stages of development. The color ranges from grey for low correlation coefficients (close to zero) to dark red for coefficients close to 1, with more intense colors indicating higher correlation coefficients. **(b, c)** Principal Component Analysis (PCA) of developmental stages in *D. melanogaster* **(c)** and *D. virilis* **(c)**. Each dot represents a distinct developmental stage. The first three components are shown. **(d, e)** Expression dynamics in the course of development for genes homologous to the Yamanaka factors for *D. melanogaster* (d) and *D. virilis* (e). For each gene, mean expression values are shown in various colors with shaded areas representing the standard deviation of expression across replicates.

However, several pupal stages were closer to the embryonic state than the larval state. Vice versa, the embryonic transcriptome was more similar to the pupal transcriptomes than to the larval ones, suggesting a partial reversion to the embryonic developmental programs after pupation. These findings are consistent with our previous study conducted on several holometabolous species belonging to various orders (Ozerova and Gelfand 2022), providing further evidence for the recapitulation of the embryonic transcriptional program during pupation. Additionally, a decreased similarity between the P4 stage and the embryo was observed in both *D. melanogaster* and *D. virilis*, indicating the activation of some pupal-specific transcriptional programs at the P4 stage, while earlier and subsequent stages of early pupal development are more similar to the embryonic state. Lower similarity observed in the late pupal stages is likely due to their increasing similarity to the imago, as late pupae are nearly fully developed adults. Since the imago has a transcriptome distinct from the embryo, it is expected that the latest pupal stages also exhibit lower similarity to the embryonic state.

Principal component analysis (PCA) further corroborates these findings (Fig. 1b,c). In both *D. melanogaster* and *D. virilis*, the first principal component, explaining 34% and 45% of the variance, respectively, effectively separates the four main developmental stages in the sequence of ontogeny. The second principal component, accounting for 28% of the variance in *D. melanogaster* and 18% in *D. virilis*, distinguishes the active larvae and imago stages from the developmental embryo and pupae stages, when the embryonic transcriptome program is activated. The third principal component (13% and 16% of the explained variance, respectively) primarily reflects temporal progression, with pupal stages aligning along this axis. These observations are consistent in both species, with *D. melanogaster* PCA results shown in Figure 1b and *D. virilis* results, in Figure 1c. The percentages represent the proportion of the total variance in the data.

### Identification of stage-specific genes and their associated biological processes

Differential expression analysis was performed to identify genes that exhibited significant upregulation at a specific stage compared to other stages.

The larval and imago stages exhibited the largest number of stage-specific genes, with over 1,000 genes identified for each stage in both *D. melanogaster* and *D. virilis*. We also identified approximately 900 genes that were specifically expressed during embryogenesis and all pupal stages.

To gain insights into the functions and roles of genes specifically expressed during the pupal stage, we performed gene ontology (GO) enrichment analysis (see Supplementary Tables 1-6).

Among genes upregulated during the P3 stage (the *bubble prepupa*), we found significant enrichment of terms related to *isoprenoid metabolic process* and *isoprenoid biosynthetic process* (GO:0006720 and GO:0008299, *p*-value<10^−3^ in both species). This observation suggests that lipid metabolism and protein modification are important during the P3 stage of the *Drosophila* development. However, the precise functions of isoprenoids during this stage are not yet fully understood, although the mevalonate pathway and heart formation may be linked by isoprenylation (Yi et al. 2006). Genes upregulated during the P4 stage (the *buoyant*) are enriched in the molecular function *structural constituent of cuticle* (GO:0042302, *p*-value<10^−4^). Statistically significant intersections between GO terms enriched in two species were not observed for other pupal stages, likely due to a high similarity in gene expression between temporarily close pupal stages (Fig. 1a and Supplementary Fig. 5a), yielding minimal statistical power to detect differences in the GO-term enrichment.

We also specifically analyzed several groups of genes known to be important for development. The Yamanaka factors were used as a general example of genes associated with cellular reprogramming and differentiation. Homologs of the Yamanaka factors in flies were analyzed (Fig. 1de) (Kaur et al. 2018). As expected, these factors exhibited increased expression in pupae in both species. The *Sox* genes had a strong expression peak during early pupal stages that was particularly prominent during the P1-P3 stages in *D. melanogaster*. On the contrary, the *Luna* genes had stable expression values after the onset of pupal development.

### Comparative analysis of developmental conservation and divergence in the *Drosophila* species

At the next step, a comparative analysis was performed with the goal to investigate similarities and differences in the developmental gene expression patterns and to identify a potential phylotypic stage.

The comparison between the two species, as shown in Figure 2 and Supplementary Figures 4 and 6, revealed high Spearman correlation coefficients and low Euclidean distance along the diagonal, indicating that each developmental stage demonstrated a close similarity to the corresponding stage in the other species, and therefore the stages had been correctly assigned during the sample collection.

**Figure 2.**
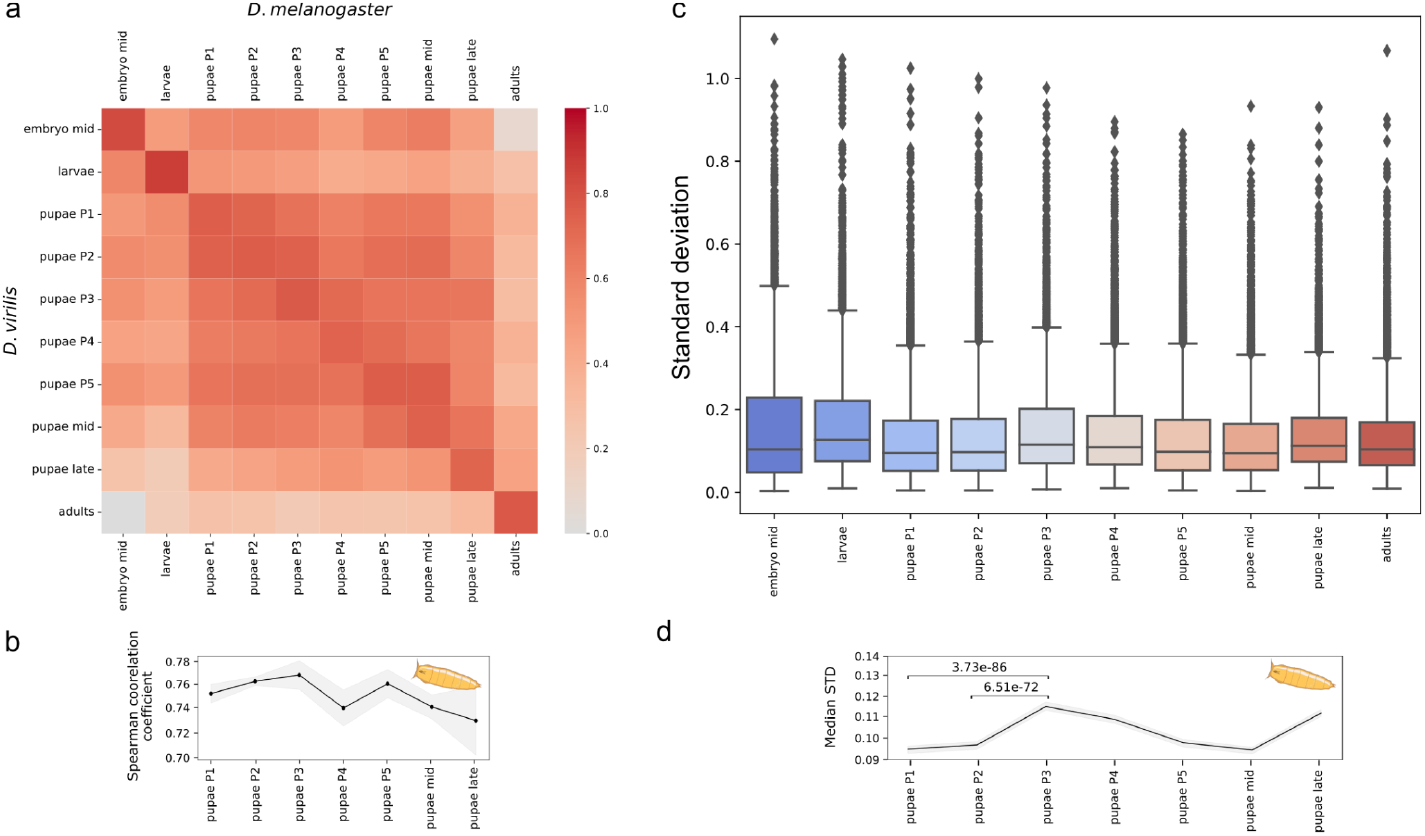
**(a)** Heatmap of the Spearman correlation coefficients for *D. melanogaster* and *D. virilis* developmental stages. Each square represents the correlation coefficient between the expression profiles of the corresponding stages between the two species. The horizontal axis lists the stages of *D. melanogaster*, while the vertical axis lists the stages of *D. virilis*. The color scheme is as in Fig. 1. **(b)** The Spearman correlation coefficients are shown as a separate line plot, representing the correlation between corresponding stages of *D. melanogaster* and *D. virilis*, that is, the main diagonal of the heatmap in (a). The x-axis represents the developmental stages, and the y-axis represents the correlation coefficients. The shaded area shows the standard deviation across replicates. **(c)** Standard deviation (STD) values for gene expression are shown for each developmental stage. The x-axis represents the developmental stages, and the y-axis represents the STD values, for details see the *Results* section. **(d)** The median STD values for each pupal stage are shown as a line plot. The x-axis represents the developmental stages, and the y-axis represents the median STD value. The shaded area around the line represents the variability of the median values estimated by downsampling: at each stage, the metric was recalculated 1000 times using a random subset of 50% of the genes. The narrow shaded area indicates that the median values are stable with respect to gene sampling.

Consistent with pupal samples being closer to embryos than to larvae in the same species, *D. melanogaster* embryos were closer to *D. virilis* pupae than to larvae, and vice versa, but again, with a distinct expression profile prominent at the P4 stage.

The main diagonal of the heatmap in Figure 2a represents the correlation between the same stages during development and is shown separately as Figure 2b, Supplementary Figures 4cd. The strongest correlation was observed at the P3 stage, followed by a decrease at the P4 stage and a subsequent peak at the P5 stage. These observations suggest a convergence in expression profiles during early and late pupal development.

The standard deviation (STD) from the mean expression across replicates was used as an additional metric of gene expression stability. Genes with more conserved expression have lower STD indicating more consistent expression across replicates, while less constrained genes would have higher STD. The distributions of STD values for each developmental stage (Figure 2c,d) exhibited dynamics similar to the Spearman correlation. Variation in gene expression increased at early pupal stages, especially at the P3 and P4 stages, indicative of decreased constraints on expression at these stages.

The P4 (buoyant) stage shows the lowest level of interspecies conservation, with both the correlation coefficient and the standard deviation indicating minimal similarity compared to neighboring pupal stages, which display higher conservation. While mid and late pupal stages suggest a gradual decline in conservation, further sampling around these timepoints is needed to accurately capture the dynamics of this trend.

### Clusters of co-expressed genes

To study gene groups with distinct developmental expression patterns, a hierarchical clustering analysis was performed using the Spearman correlation coefficient as the distance metric (Fig. 3). To focus on the dynamics of pupae, only pupal stages were considered at the clustering step, yielding eight clusters.

**Figure 3.**
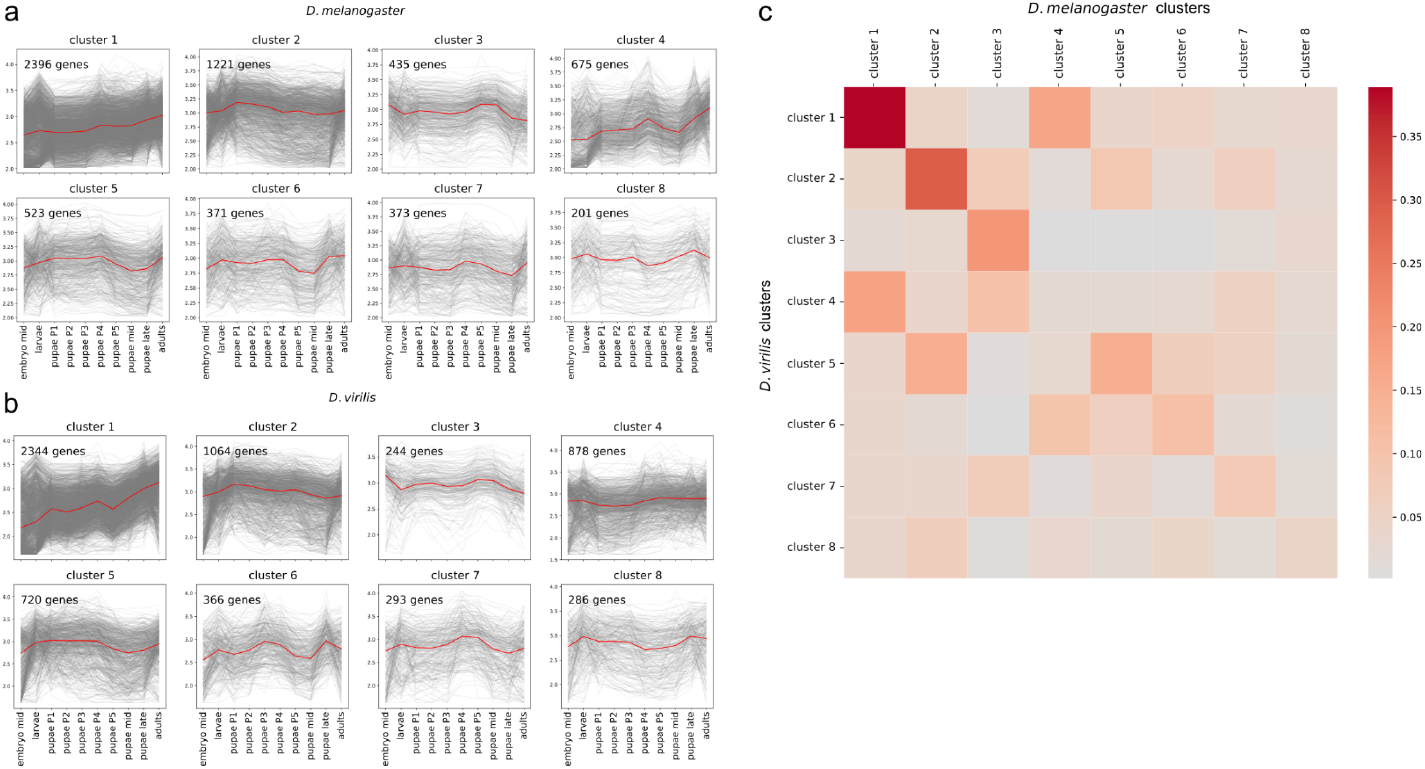
Clustering of expression profiles. Developmental expression profiles for each of the constructed clusters for *D. melanogaster* **(a)** and *D. virilis* **(b)**. Each gene is represented by a gray line, and the median expression profile for each cluster is shown in red. **(c)** Heatmap illustrating the intersection of gene sets in each cluster pair. Each square represents a cluster pair, with the color scheme shown at the right; darker colors indicate higher fractions of common genes.

Clusters were sorted according to the number of common orthologous genes, so that similar pairs of clusters had the same names in both species (Fig. 3b). In most cases the median gene trajectories of paired clusters turned out to be highly similar between the two species, despite independent clustering in the species (Supplementary Fig. 7).

Cluster pair 1 was the largest, had the largest set of common genes and was composed of genes with expression gradually increasing throughout the lifespan in both *D. melanogaster* and *D. virilis*.

The observed similarities demonstrate that orthologous genes tend to have similar dynamics (statistics provided in Supplementary Fig. 7), supporting the biological relevance of the clusters. Therefore, we selected pairs of clusters exhibiting high correlation of expression profiles and containing common orthologous genes, and performed GO enrichment analysis to reveal the functional role of these clusters.

Expression of genes comprising cluster pair 2 increased during early stages of the pupal development followed by a sequential gradual decrease. The enriched GO terms for this cluster were related to the regulation of growth, development and cell migration (Supplementary Fig. 8). Genes in this cluster pair have a pupae-specific expression, since in embryogenesis their expression is relatively low.

Genes in cluster pair 3 had a higher correlation of gene expression between the species for all developmental stages compared to the average correlation (Supplementary Fig. 9). Expression of these genes increased during the P5 and mid-pupae stages and also significantly increased during the embryonic stage. This suggests that these genes are involved in the transcriptome programs of both embryogenesis and metamorphosis. GO enrichment analysis revealed an enrichment of terms related to chromatin organization, cell cycle, and developmental processes.

Cluster 7 in both species has an expression peak at the later pupal stages while also maintaining notable expression in larvae. However, the overlap of genes within these clusters between the two species is relatively small (Figure 3c), indicating that these genes are characterized by species-specific expression patterns rather than a conserved expression pattern across species. Thus, we have not pursued a detailed analysis of these clusters.

GO enrichment results for other cluster pairs with a significant number of intersecting genes and statistically reliable outcomes are given in Supplementary Fig. 10-13.

### Gene age influence on the developmental transcriptome

To gain insights into the evolutionary origin of the observed transcriptome dynamics, the phylostrata analysis was performed. Each gene was assigned to a phylostratum corresponding to its predicted time of emergence (for details, see *Materials and Methods* section).

Genes belonging to different phylostrata exhibit specific temporal dynamics of expression, as shown in Figure 4, while in general having a tendency to increase the expression rate during ontogeny. The most ancient genes (gray) tend to have a higher expression at the larval stage, suggesting their involvement in housekeeping functions associated with larval feeding and growth. Intermediate-aged genes (blue), common to *Arthropoda, Pancrustacea*, and *Insecta*, have an additional prominent expression peak at the P4 stage and in late pupae. Following the trend, the group with the youngest, *Holometabola*-specific genes has only pupa-specific peaks, indicating that these newly acquired genes could have been selected in *Holometabola* to perform metamorphosis-related functions.

**Figure 4.**
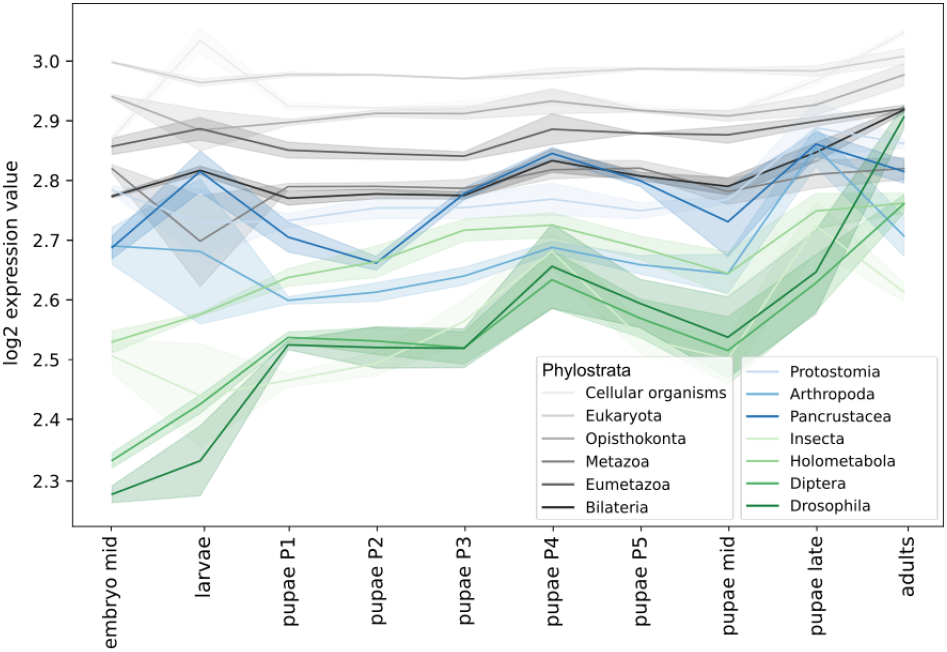
Phylostrata-specific gene expression dynamics. Horizontal axis — the developmental stage, vertical axis — the median log-transformed expression. The shaded area represents the standard deviation (STD) of gene expression. The oldest phylostrata (see the legend) are shown in gray, middle phylostrata, in blue, and the youngest ones, in green.

Many young genes up-regulated during the P4 stage encode proteins with candidate signal peptides that control protein secretion (see *Materials and Methods* for the details). However, signal peptides tend to appear in young genes in general (Supplementary Fig. 14). The distributions of all genes and signal peptide-encoding genes across the phylostrata are statistically different (*p*-value = 8.3×10^−75^, statistic = 384.2, the Chi-squared test), and P4-specific genes follow the same trend with no observed statistically significant difference (the Chi-squared statistic = 0.2).

Genes from different phylostrata exhibit varying impacts on the overall similarity of transcriptomes during metamorphosis, as seen in the plots of correlation between gene expression at each stage, calculated separately for each phylostratum (Fig. 5). For intermediate-aged genes (Fig. 5a, blue), the correlation between expression in two species was larger at the later pupal stages. Thus, the expression of these genes is conserved closer to the end of the pupal development. Genes common to *Pancrustacea* have an additional conservation peak during earlier stages. Conversely, younger genes (Fig. 5a, green) exhibited a higher correlation between species primarily at the onset of the pupal development, when imaginal disks are everted and active differentiation occurs. Notably, however, *Holometabola*-specific genes within this group deviate from this general trend, showing a pattern more similar to that of genes of intermediate ages.

**Figure 5.**
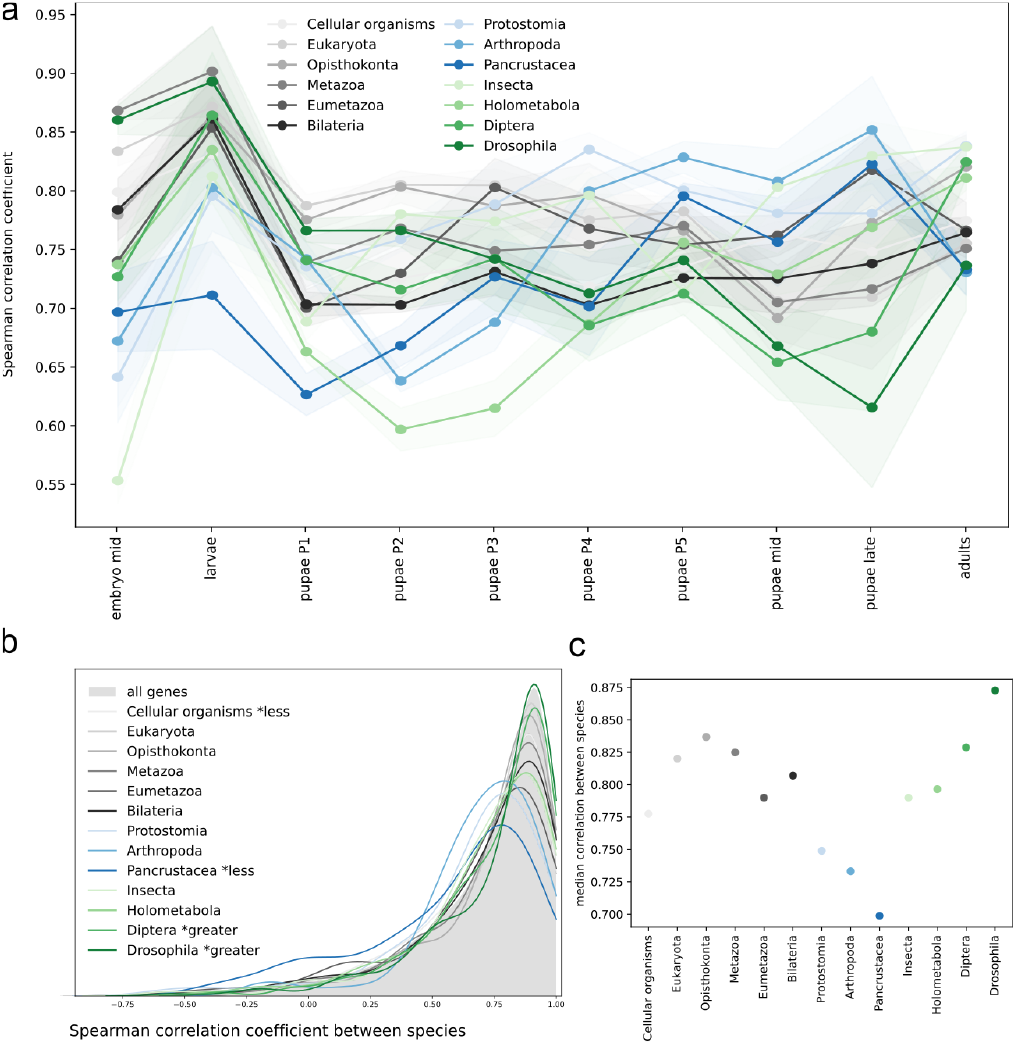
Phylostrata-based analysis of conservation of gene expression. **(a)** Correlation of gene expression profiles in two species, calculated separately for each phylostratum. Each line represents the average correlation coefficient between species with the shaded areas showing the standard deviation across replicates. **(b)** Density plot for the Spearman correlation coefficients for gene expression in two species across development, calculated separately for each phylostratum. The density plot for all genes is shown as the shaded area. **(c)** Median values for the distribution of correlation coefficients from (b). The color code is as in Figure 4.

Looking at another angle, that is, comparing the expression of individual genes throughout development (Fig. 5b, c), we observed that recently emerged genes from the *Diptera* and *Drosophila* phylostrata tended to have a higher correlation between species (*p*-values for the one-sided Mann–Whitney U-test are 9.7×10^−3^ and 1.2×10^−5^, respectively). This suggests that either these genes had less evolutionary time to diverge, or their function is crucial for pupae, leading to negative selection on changes in expression. In contrast, genes belonging to the *Pancrustacea* phylostratum exhibited a lower correlation than the average (*p*-value equal to 4×10^−6^).

The median correlation coefficient for genes comprising different phylostrata tends to decrease with the decrease of the predicted age, see old (grey) and middle (blue) phylostrata in Fig. 5c. However a breakpoint is observed after the transition from *Pancrustacea* to *Insecta* and this tendency is reversed, so that the younger the phylostratum (green in Fig. 5c), the higher is the median correlation between the species. The most similar expression throughout the development was observed for the oldest and the youngest genes.

Clusters 4 and 6 (with increased expression at mid-pupae) were enriched with both young, *Drosophila*-specific genes, genes from the *Insecta* phylostratum, and old genes (Fig. 6). Genes in cluster 4 had an expression burst at the P4 stage, while cluster 6 featured higher expression at the early pupal and adult stages. Cluster 5, consisting of genes expressed at early pupal stages, was dominated by middle phylostrata. Cluster 1, characterized by a constantly increasing expression profile throughout the development, was strongly enriched with the oldest genes and *Bilateria* genes.

**Figure 6.**
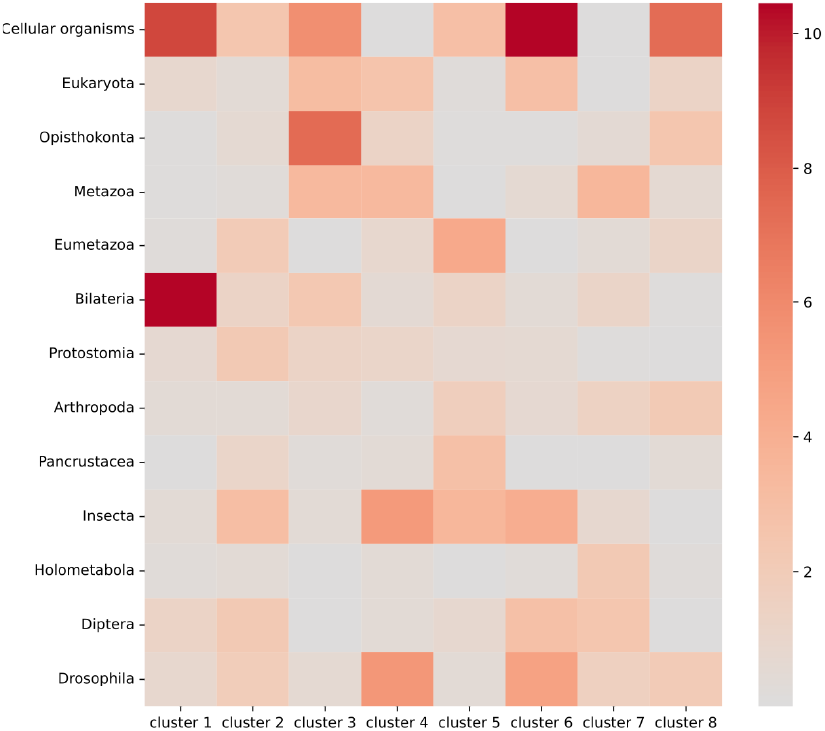
Intersection between phylostrata and clusters. The color of the square corresponds to the significance of the intersection (see *Materials and Methods* for details).

## Discussion

The hourglass model of gene expression, characterized by a phase of higher difference in gene expression at the beginning and end of development with a period of higher conservation in the middle, phylotypic, stage is well-established in the context of embryogenesis (Irie and Kuratani 2014; Drost et al. 2017). This pattern has been documented across a wide range of species ranging from insects and nematodes to mammals (Kalinka et al. 2010; Levin et al. 2012; Hazkani-Covo, Wool, and Graur 2005). Furthermore, it may extend beyond embryogenesis and, in plants, is also observed in processes like the floral transition (Drost et al. 2016), suggesting that this model has broader implications than initially thought. Our initial hypothesis has been that the pupal development of holometabolous insects, when organogenesis re-starts, adheres to a similar hourglass-like pattern. This was based on our observation that pupae tend to recapitulate embryogenesis transcription patterns (Ozerova and Gelfand 2022). The expectation was that a phase of increased similarity in gene expression might occur during the mid-pupal stages.

However, contrary to the classic hourglass model, our findings indicate that while there are stages of conserved gene expression, they predominantly occur during early metamorphosis stages, rather than in the middle (Fig. 2, Supplementary Fig. 4 and 6). Moreover, we observe that the P4 stage is the most divergent stage in terms of gene expression between the species, as measured by correlation coefficients, standard deviation, and the Euclidean distance (Fig. 2a,b, Supplementary Fig. 6). A previous microarray study also identified the P4 stage as being the least similar to the embryonic stages compared to other pupal stages in *D. melanogaster* (Arbeitman et al. 2002). This suggests a spindle-like pattern, where the transcriptome divergence is accentuated during this phase. The main process during this stage is the eversion of the imaginal head sac. Reorganization of the respiratory system could be visually observed since the lateral trunk tracheae becomes obscured. Also, the oral armature of the larva is expelled, signaling the complete removal of larval feeding structures as the organism prepares for its final differentiation into the adult form (Bainbridge and Bownes 1981). Overall, this underscores the dynamic nature of gene expression during pupal transformation, likely driven by the relatively recent emergence of the latter.

An intriguing observation is that genes expressed during the P4 stage, a period characterized by high divergence, tend to be evolutionarily new. For example, clusters 4 and 6, which have an increased expression at this stage, are enriched with young genes (Fig. 6). Moreover, while being relatively highly expressed, these recently emerged genes (specific for *Diptera* and *Drosophila*) demonstrate a rather low correlation in expression at the P4 stage compared to the adjacent stages (Fig. 5a).

To place these findings in the evolutionary context, the *dN/dS* ratio for *D. melanogaster* genes, taken from (Mensch et al. 2013), was used as a measure of natural selection and evolutionary rate. While genes with higher sequence conservation (low *dN/dS* values) have higher expression levels for all stages except the imago (Supplementary Figure 15a), genes from cluster 4 with the expression burst during the P4 stage demonstrate significantly higher substitution rates compared to the complete gene set (*p*-value = 0.006, Supplementary Figure 15c). However, genes upregulated exclusively during P4 do not feature a statistically significant difference in the *dN/dS* ratio, likely due to the small number of genes specifically upregulated during this short stage.

Conversely, genes, upregulated at the earliest prepupal stage (P1) when the expression levels are highly coordinated (Fig. 2, Supplementary Fig. 6) (*p*-value = 0.012, Supplementary Fig. 15b) and genes from cluster 2, which show peak expression at P1 (*p*-value = 0.002, Supplementary Fig. 15c), have significantly lower *dN/dS* ratios and hence are more conserved.

Remarkably, we find that genes with a middle predicted age (emerging in *Arthropoda* and *Pancrustacea*) have conserved expression patterns in the studied *Drosophila* species during later pupal stages (Fig. 5a). The expression range of these genes remains conserved, suggesting that they did not diverge after the emergence. Indeed, genes from these phylostrata are outliers in the *dN/dS* ratio distribution (Supplementary Fig. 15d), whereas genes from younger phylostrata tend to display higher *dN/dS* ratios.

Our study provides insights into the transcriptional dynamics of Drosophila metamorphosis, shedding light on the temporal patterns of gene expression during this critical developmental process. However, delving deeper into the temporal dynamics by analyzing additional stages could offer a more comprehensive understanding of the metamorphosis. By including more initial and intermediate stages, we could capture finer transitions and potentially identify novel regulatory mechanisms underlying metamorphic events. Moreover, analyzing a wider range of *Drosophila* species would enable us to refine the timing of key pupal developmental stages and enhance the statistical power of our comparative analyses.

Additionally, expanding the scope of comparisons beyond *Drosophila* species — studying insects with mobile pupae, such as mosquitoes, presents a unique opportunity to investigate the preservation of larval structures during pupal development. Unlike *Drosophila*, whose immobile pupae undergo extensive tissue remodeling, insects with mobile pupae should retain larval organs due to the need for locomotion and feeding during the pupal stage. Investigating the molecular mechanisms and gene expression dynamics underlying these differences between species with fixed and mobile pupae could uncover fundamental insights into the evolutionary diversity of metamorphic strategies across insects.

## Materials and Methods

### Sample collection

Wild-type *D. melanogaster* (Oregon-R) and wild-type *D. virilis* (Strain 9) were used for RNA sequencing. These two distant species were selected for the analysis because they belong to two different subgenera, *Sophophora* and *Drosophila*, respectively (O’Grady and DeSalle 2018). Therefore, the observed developmental patterns should be common for the whole *Drosophila* genus.

The flies, spanning all developmental stages, were reared on the standard *Drosophila* wheat meal-yeast-sugar-agar medium and maintained at a constant temperature of 25°C. Since the developmental rate of *D. virilis* is approximately 1.5 times slower than that of *D. melanogaster*, to account for the intrinsic differences in developmental timing between the two species, the temporal synchrony of developmental stages was monitored by visual observation.

In order to synchronize the ages of collected embryos, adult flies were allowed to lay eggs in Petri dishes containing a 3% agarose substrate covered with apple juice and yeast for one hour interval three times a day. Subsequently, fresh dishes were used for embryo collection. The collection process took place for a one-hour interval, followed by incubation of the dishes at an appropriate temperature until the embryos reached the phylotypic stage, which required 10 hours in *D. melanogaster* and 15 hours in *D. virilis*. Each biological replicate consisted of approximately 100 embryos.

At the next stage the dishes with synchronized one-hour eggs were incubated at an appropriate temperature to collect certain larval stages. All biological replicates for *D. virilis* larvae were collected approximately midway between molts at the second instar stage (72 hours), with each sample consisting of 10–12 individuals. *D. melanogaster* larvae were also collected approximately midway between molts at all three larval stages, with first-instar samples (24 hours) containing 20–25 individuals and second- and third-instar samples consisting of 15–18 individuals each (48 and 72 hours respectively). All larval samples were treated as replicates, and expression levels were averaged across them to create a single representative larval expression profile for downstream analysis.

Pupae were also collected individually with selective individual monitoring, without distinguishing males and females, including stages P1, P2, P3, P4, and P5, as per the developmental stage descriptions outlined by (Bainbridge and Bownes 1981). Furthermore, intermediate (24 and 36 hours of development for *D. melanogaster* and *D. virilis*, respectively) and late (48 and 72 hours of development for *D. melanogaster* and *D. virilis*, respectively) pupal stages were collected. Each biological replicate consisted of 10–12 individuals for *D. virilis* and 15–18 individuals for *D. melanogaster*. A particular focus was placed on the early pupal stages (P1-P5), as they encompass key developmental transitions, including the reorganization of larval tissues and the differentiation of adult structures. These early stages are particularly dynamic and critical for metamorphosis, marking major shifts in gene expression and tissue remodeling. In contrast, later pupal stages primarily involve growth and refinement processes, with the developing organism already resembling the adult form.

Adult samples consisted of five-day-old males, with each biological replicate containing 10 individuals for *D. virilis* and 15 for *D. melanogaster*. Adult females were excluded from the experiment to avoid the influence of the ovaries with eggs, as in this case the full transcriptome would be skewed towards the egg state. The influence of sex on other stages was considered to be negligible, and each biological replicate was expected to consist of an almost equal proportion of males and females, although the sex ratio in non-adult stages was not directly verified.

All samples were collected in three independent biological replicates. Subsequently, the samples underwent a thorough rinsing process to remove any residual food and yeast, using the PBS buffer. Following this step, total RNA was extracted from the samples using RNAzol RT reagent (MRC, Cincinnati, OH, USA), following the manufacturer’s recommended protocols. RNA quality assessments were performed through gel electrophoresis, and the RNA samples were carefully stored in ethanol at −70°C, in preparation for subsequent experimental procedures.

### Sequencing

Ribosomal RNA depletion was conducted using the NEBNext® Poly(A) mRNA Magnetic Isolation Module. Subsequently, RNA library preparation was carried out with the NEBNext® Ultra™ II Directional RNA Library Prep Kit, and the prepared libraries were sequenced on the Illumina HiSeq4000 platform in the Skoltech Genomics Core Facility. The sequencing yielded paired-end reads with the length of 150 nucleotides. Due to the suboptimal quality observed in the replicates of *D. melanogaster* 2^nd^ instar larvae, we made the decision to use the 1^st^ and 3^rd^ instar samples for the downstream analyses.

### Processing of transcriptome data

Low-quality nucleotides and reads were systematically eliminated using the `fastp` tool with the default parameters. Subsequently, the filtered reads were processed using the `Kallisto` pseudomapping procedure to estimate transcript abundances, with BDGP6.46 and DvirRS2 transcriptomes used as references for *D. melanogaster* and *D. virilis*, respectively. Transcript-level expression estimates were summed to obtain gene-level expression data. Based on the observed high reproducibility, gene expression values were averaged across replicates using the arithmetic mean. For downstream analyses, variance stabilizing transformation (`VST`) from the `DESeq2` package (`Bioconductor`, `R`) was applied to the raw counts.

To mitigate noise, genes expressed at low levels and exhibiting limited variance across the developmental stages were excluded from the subsequent analyses. The threshold for the expression values was set as the inflection point of the expression values density plot, separating the bell-shaped part of the distribution from low values (95% and 93% of genes are preserved for *D. melanogaster* and *D. virilis* respectively). The variance threshold was set on the first inflection point of the cumulative distribution. Overall, 69% of genes passed the specified filters for both species.

### Orthologous groups and functional analysis

Orthologous gene groups were downloaded from the `Ensembl` database using the `Biomart` tool. Subsequently, only genes belonging to orthologous groups shared by both species were considered for the subsequent analyses. Gene ontology (GO) term annotations were obtained from the `AmiGO2` database for *D. melanogaster* and from the `Ensembl` database for *D. virilis*.

GO term enrichment analysis was performed using the `goatools` Python package (Klopfenstein et al. 2018) with the default settings and an adjusted *p*-value threshold of 0.05. The analysis used a reference set of all genes showing positive expression values in the analyzed dataset. The semantic analysis of the enriched GO terms was visualized using the `REVIGO` tool (Supek et al. 2011) and plotted with the R `ggplot` package.

To predict the presence of signal peptides in the protein sequences, the `SignalP-6.0` tool was used with the default eukaryotic settings (Teufel et al. 2022).

### Data analysis

Ad hoc `Python` scripts were used for statistical testing and visualization (Klopfenstein et al. 2018), with figures were prepared using the matplotlib and seaborn packages. Statistical tests were performed using the `Scipy` package.

### Analysis of phylostrata

For the phylostrata annotation of *D. melanogaster* genes, a phylostratigraphy approach was used (Domazet-Loso, Brajković, and Tautz 2007). For example, a gene was assigned to the *Holometabola* phylostratum if its homologs were found exclusively within the *Holometabola* group, indicating an estimated age of approximately 340 million years.

To assess the significance of enrichment for each phylostratum in each cluster, the following ratio was used:

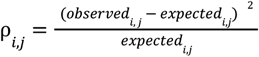

where *observed*_*i,j*_ is the number of common genes in the intersection of cluster *i* and phylostratum *j*, and 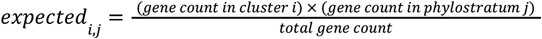. This metric highlights substantial intersections between clusters and phylostrata, while adjusting for the overall gene count within each group.

### Identification of stage-specific genes

Stage-specific genes were identified using differential expression analysis with the `DESeq2` package (`Bioconductor`, `R`), applied separately for each species. A gene was classified as stage-specific if it met two criteria: (1) it was significantly upregulated at a particular stage, with an adjusted *p*-value < 0.05 and a positive log-fold change, and (2) it was not upregulated at any other stage, except for the adjacent ones. Differential expression was assessed by comparing gene expression at the stage of interest against all other developmental stages of the same species, using raw count data.

## Acknowledgements

This study was supported by the Russian Science Foundation via grant 23-14-00136.

We acknowledge the Skoltech Genomics Core Facility for library preparation and sequencing supported by the Skoltech Center of Molecular and Cellular Biology grant for the students’ projects.

Experiments were conducted using the equipment of the Core Centrum of the Institute of Developmental Biology RAS, supported by the IDB RAS research program 0088-2024-0010.

We also acknowledge Ilya Borisovitch Mertsalov for technical assistance in the collection of specimens.

## Data availability

The RNA sequencing data that support the findings of this study are available in the NCBI GEO data repository under the accession GSE269951.

## Author contributions statement

Ozerova AM* - Data curation, Formal analysis, Investigation, Methodology, Software, Validation, Visualization, Writing – original draft

Kulikova DA - Investigation, Methodology, Writing – review & editing

Evgen’ev MB - Resources, Supervision, Writing – review & editing

Gelfand MS - Conceptualization, Funding acquisition, Methodology, Project administration, Resources, Supervision, Validation, Writing – review & editing

## References

Arbeitman, Michelle N., Eileen E. M. Furlong, Farhad Imam, Eric Johnson, Brian H. Null, Bruce S. Baker, Mark A. Krasnow, Matthew P. Scott, Ronald W. Davis, and Kevin P. White. 2002. “Gene Expression during the Life Cycle of Drosophila Melanogaster.” Science 297 (5590): 2270–75.

Bainbridge, S. P., and M. Bownes. 1981. “Staging the Metamorphosis of Drosophila Melanogaster.” Journal of Embryology and Experimental Morphology 66 (December):57–80.

Domazet-Loso, Tomislav, Josip Brajković, and Diethard Tautz. 2007. “A Phylostratigraphy Approach to Uncover the Genomic History of Major Adaptations in Metazoan Lineages.” Trends in Genetics: TIG 23 (11): 533–39.

Domazet-Lošo, Tomislav, and Diethard Tautz. 2010. “A Phylogenetically Based Transcriptome Age Index Mirrors Ontogenetic Divergence Patterns.” Nature 468 (7325): 815–18.

Drost, Hajk-Georg, Julia Bellstädt, Diarmuid S. Ó’Maoiléidigh, Anderson T. Silva, Alexander Gabel, Claus Weinholdt, Patrick T. Ryan, et al. 2016. “Post-Embryonic Hourglass Patterns Mark Ontogenetic Transitions in Plant Development.” Molecular Biology and Evolution 33 (5): 1158–63.

Drost, Hajk-Georg, Philipp Janitza, Ivo Grosse, and Marcel Quint. 2017. “Cross-Kingdom Comparison of the Developmental Hourglass.” Current Opinion in Genetics & Development 45 (August):69–75.

Duboule, D. 1994. “Temporal Colinearity and the Phylotypic Progression: A Basis for the Stability of a Vertebrate Bauplan and the Evolution of Morphologies through Heterochrony.” Development. Supplement, 135–42.

Hazkani-Covo, Einat, David Wool, and Dan Graur. 2005. “In Search of the Vertebrate Phylotypic Stage: A Molecular Examination of the Developmental Hourglass Model and von Baer’s Third Law.” Journal of Experimental Zoology. Part B, Molecular and Developmental Evolution 304 (2): 150–58.

Irie, Naoki. 2017. “Remaining Questions Related to the Hourglass Model in Vertebrate Evolution.” Current Opinion in Genetics & Development 45 (August):103–7.

Irie, Naoki, and Shigeru Kuratani. 2014. “The Developmental Hourglass Model: A Predictor of the Basic Body Plan?” Development 141 (24): 4649–55.

Kalinka, Alex T., Karolina M. Varga, Dave T. Gerrard, Stephan Preibisch, David L. Corcoran, Julia Jarrells, Uwe Ohler, Casey M. Bergman, and Pavel Tomancak. 2010. “Gene Expression Divergence Recapitulates the Developmental Hourglass Model.” Nature 468 (7325): 811–14.

Kaur, Prameet, Helen Jingshu Jin, Jay B. Lusk, and Nicholas S. Tolwinski. 2018. “Modeling the Role of Wnt Signaling in Human and Drosophila Stem Cells.” Genes 9 (2). 10.3390/genes9020101.

Klopfenstein, D. V., Liangsheng Zhang, Brent S. Pedersen, Fidel Ramírez, Alex Warwick Vesztrocy, Aurélien Naldi, Christopher J. Mungall, et al. 2018. “GOATOOLS: A Python Library for Gene Ontology Analyses.” Scientific Reports 8 (1): 10872.

Levin, Michal, Tamar Hashimshony, Florian Wagner, and Itai Yanai. 2012. “Developmental Milestones Punctuate Gene Expression in the Caenorhabditis Embryo.” Developmental Cell 22 (5): 1101–8.

Mayshar, Yoav, Ofir Raz, Saifeng Cheng, Raz Ben-Yair, Ron Hadas, Netta Reines, Markus Mittnenzweig, et al. 2023. “Time-Aligned Hourglass Gastrulation Models in Rabbit and Mouse.” Cell 186 (12): 2610–27.e18.

Mensch, Julián, François Serra, Nicolás José Lavagnino Hernán Dopazo, and Esteban Hasson. 2013. “Positive Selection in Nucleoporins Challenges Constraints on Early Expressed Genes in Drosophila Development.” Genome Biology and Evolution 5 (11): 2231–41.

Micchelli, Craig A., Lisa Sudmeier, Norbert Perrimon, Shan Tang, and Ryan Beehler-Evans. 2011. “Identification of Adult Midgut Precursors in Drosophila.” Gene Expression Patterns: GEP 11 (1-2): 12–21.

Mora, Camilo, Derek P. Tittensor, Sina Adl, Alastair G. B. Simpson, and Boris Worm. 2011. “How Many Species Are There on Earth and in the Ocean?” PLoS Biology 9 (8): e1001127.

O’Grady Patrick M., and Rob DeSalle. 2018. “Phylogeny of the Genus Drosophila.” Genetics 209 (1): 1–25.

Ozerova, Alexandra M., and Mikhail S. Gelfand. 2022. “Recapitulation of the Embryonic Transcriptional Program in Holometabolous Insect Pupae.” Scientific Reports 12 (1): 17570.

Raff, Rudolf A. 2012. The Shape of Life: Genes, Development, and the Evolution of Animal Form. University of Chicago Press.

Richardson, M. K. 1995. “Heterochrony and the Phylotypic Period.” Developmental Biology 172 (2): 412–21.

Schultheis, Dorothea, and Manfred Frasch. 2023. “Longitudinal Visceral Muscles in Drosophila Fully Dedifferentiate and Fragment prior to Their Reestablishment during Metamorphosis.” microPublication Biology 2023 (March). 10.17912/micropub.biology.000756.

Supek, Fran, Matko Bošnjak, Nives Škunca, and Tomislav Šmuc. 2011. “REVIGO Summarizes and Visualizes Long Lists of Gene Ontology Terms.” PloS One 6 (7): e21800.

Tamura, Koichiro, Sankar Subramanian, and Sudhir Kumar. 2004. “Temporal Patterns of Fruit Fly (Drosophila) Evolution Revealed by Mutation Clocks.” Molecular Biology and Evolution 21 (1): 36–44.

Ten Brink, Hanna, André M. de Roos, and Ulf Dieckmann. 2019. “The Evolutionary Ecology of Metamorphosis.” The American Naturalist 193 (5): E116–31.

Teufel, Felix, José Juan Almagro Armenteros, Alexander Rosenberg Johansen, Magnús Halldór Gíslason, Silas Irby Pihl, Konstantinos D. Tsirigos, Ole Winther, Søren Brunak, Gunnar von Heijne, and Henrik Nielsen. 2022. “SignalP 6.0 Predicts All Five Types of Signal Peptides Using Protein Language Models.” Nature Biotechnology 40 (7): 1023–25.

Texada, Michael J., Takashi Koyama, and Kim Rewitz. 2020. “Regulation of Body Size and Growth Control.” Genetics 216 (2): 269–313.

Truman, J. W. 1990. “Metamorphosis of the Central Nervous System of Drosophila.” Journal of Neurobiology 21 (7): 1072–84.

Yi, Peng, Zhe Han, Xiumin Li, and Eric N. Olson. 2006. “The Mevalonate Pathway Controls Heart Formation in Drosophila by Isoprenylation of Ggamma1.” Science 313 (5791): 1301–3.

