## Supplementary Figures 1-15 for "Temporal dynamics of gene expression during metamorphosis in two distant *Drosophila* species"

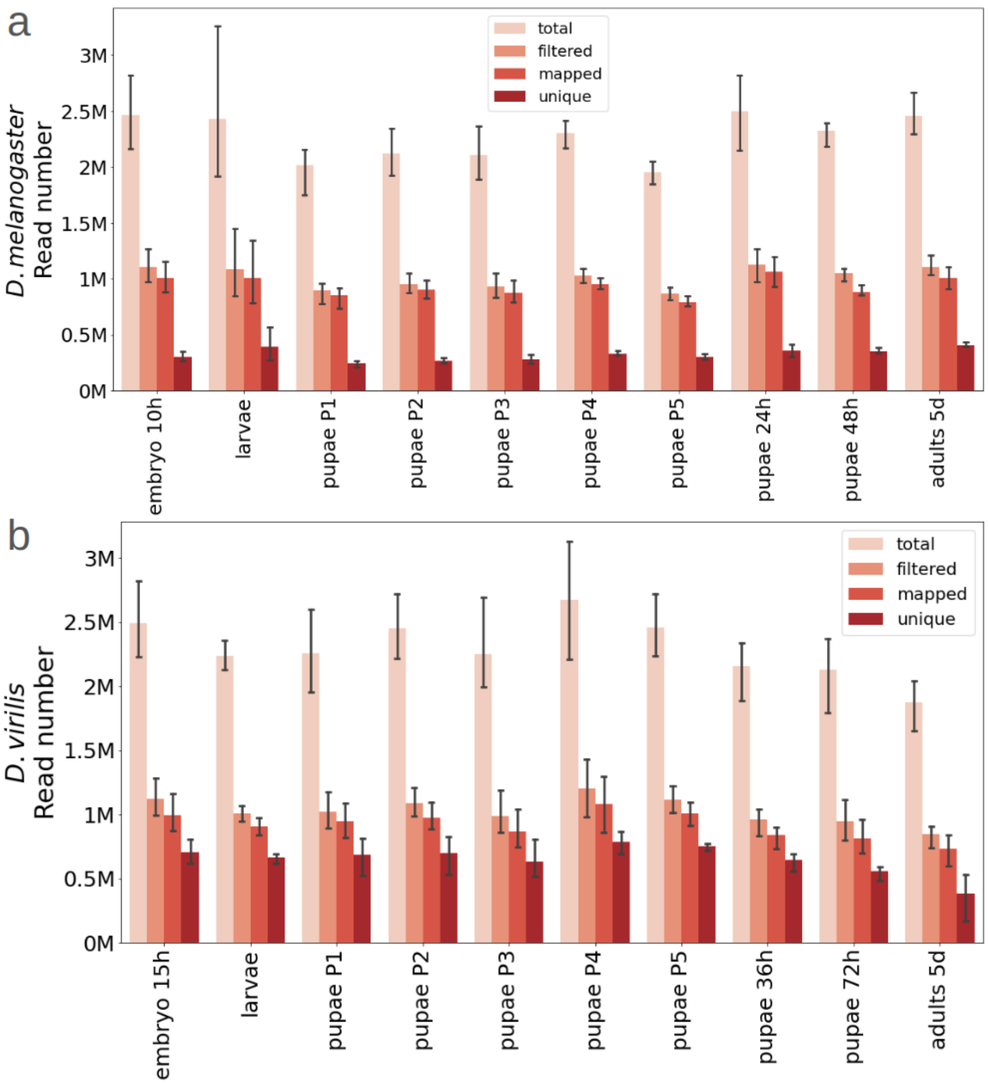


**Supplementary Fig. 1.** Summary statistics of read processing and mapping. The number of reads at different processing stages is shown for each developmental stage: total sequenced, filtered, mapped, and uniquely mapped. **(a)** *D. melanogaster*, **(b)** *D. virilis*.

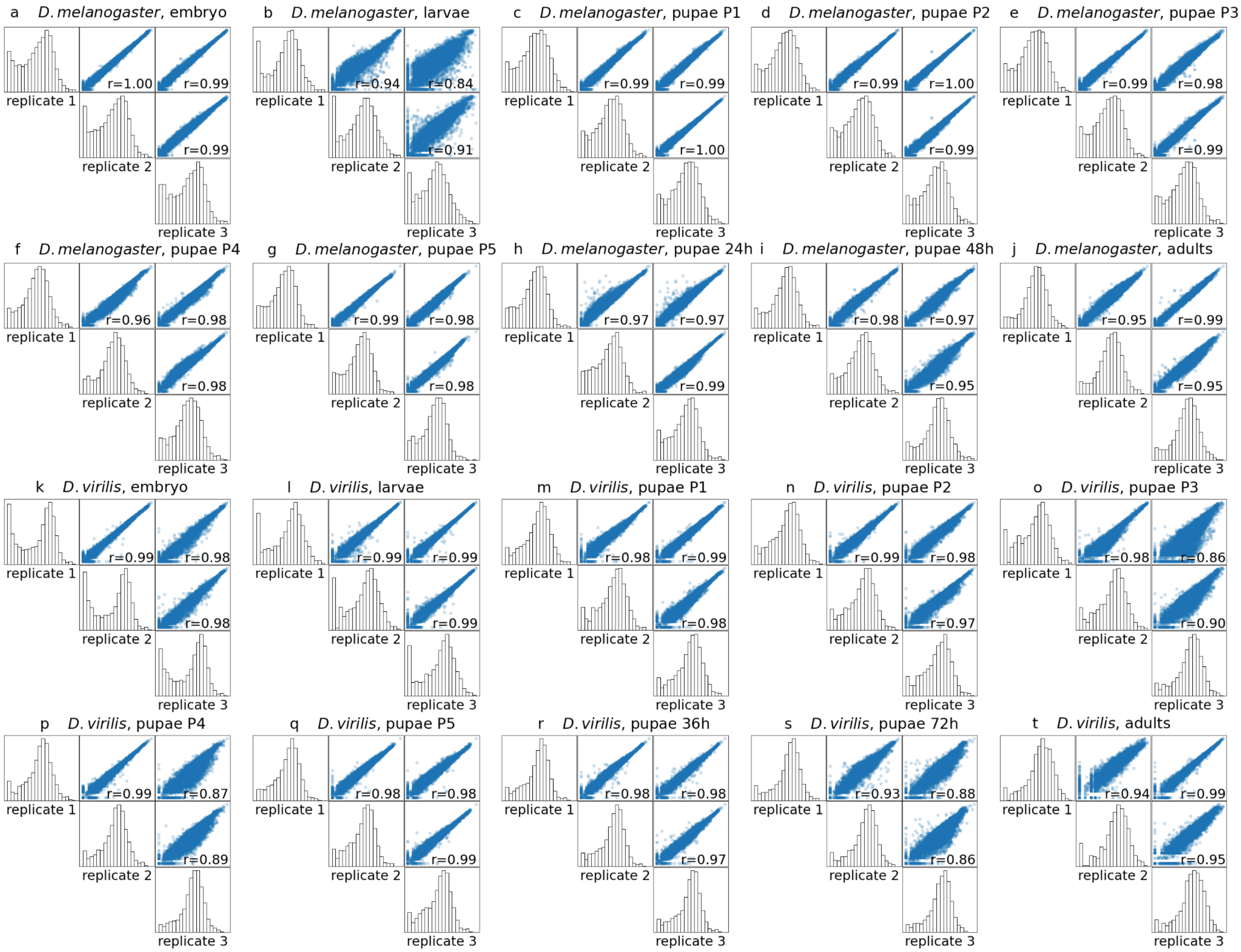


**Supplementary Fig. 2.** Reproducibility of RNA sequencing data. Each half-matrix corresponds to a specific developmental stage of *D. melanogaster* **(a-j)** or *D. virilis* **(k-t)**. Histograms on the diagonal represent the distribution of log-transformed gene expression values for a given developmental stage. Scatter plots above the histograms show pairwise comparisons of gene expression between three biological replicates of the corresponding stage. The Spearman correlation coefficients (*r*) are given.

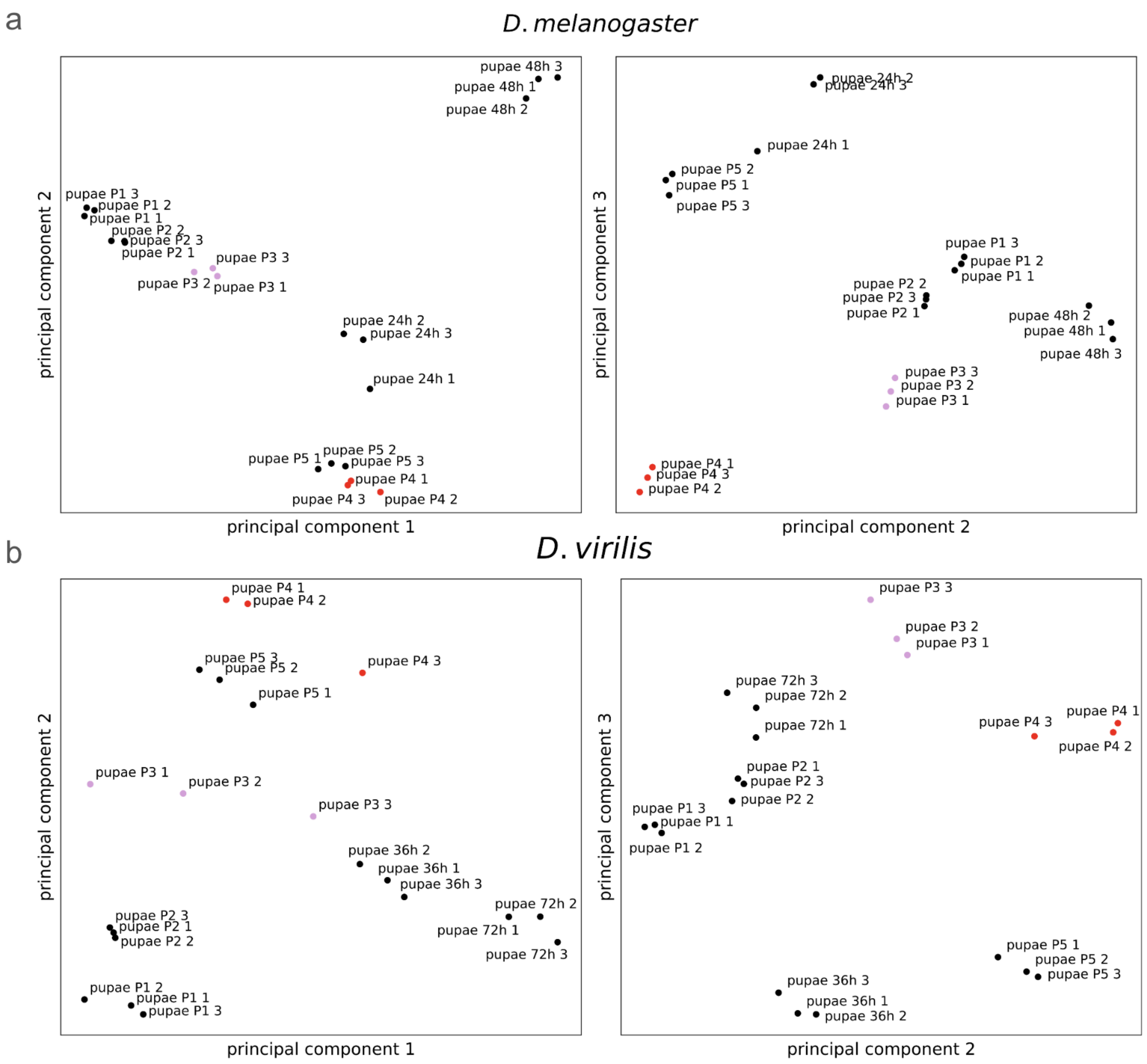


**Supplementary Fig. 3.** Principal component analysis for **(a)** *D.melanogaster* and **(b)** *D. virilis* pupal replicates.

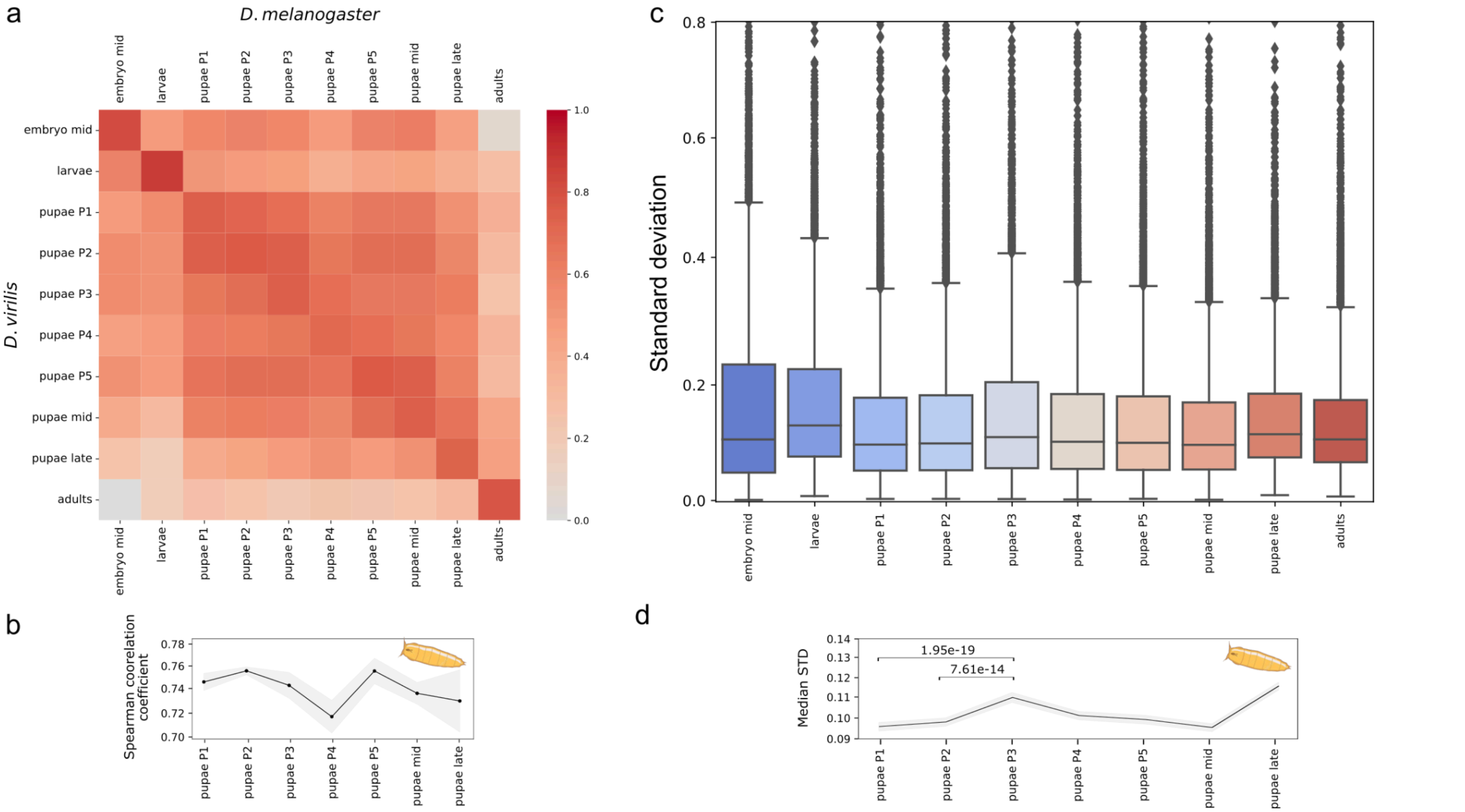


**Supplementary Fig. 4.** Analysis after excluding questionable replicates of pupae P3 and P4 (see Supplementary Fig. 3) from the *D. virilis* dataset. Notation as in Figure 2.

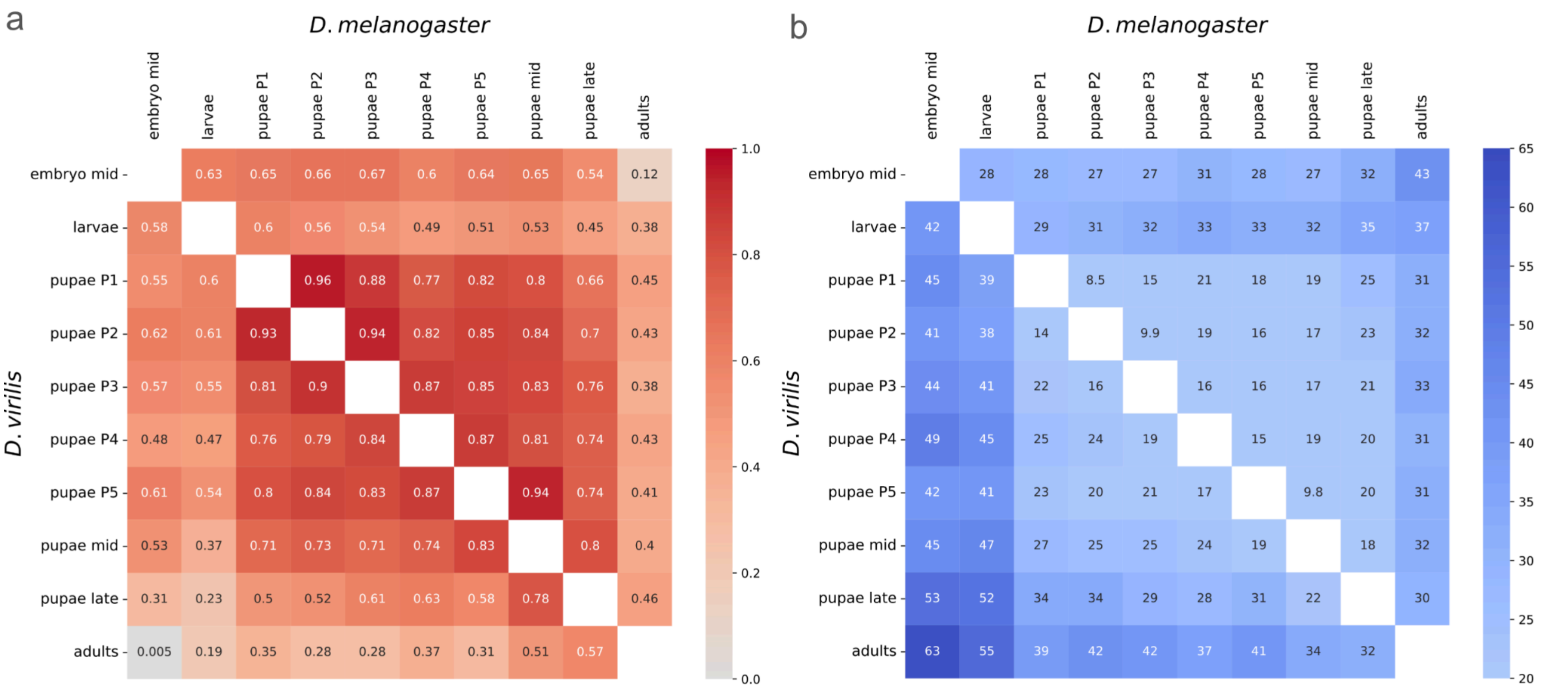


**Supplementary Fig. 5.** Distance heatmaps of the developmental stages for *D. melanogaster* (above the diagonal) and *D. virilis* (below the diagonal) analyzed separately for each species. **(a)** Each square represents the Spearman correlation coefficient between the respective stages of development. The color ranges from grey for low correlation coefficients (close to zero) to dark red for coefficients close to 1, with more intense colors indicating higher correlation coefficients. **(b)** Each square represents the Euclidean distance between the respective stages of development. The color ranges from light blue for small distances (closer to zero) to dark blue for larger ones, with more intense colors indicating larger distance. The Euclidean distance was calculated using log-transformed expression values.

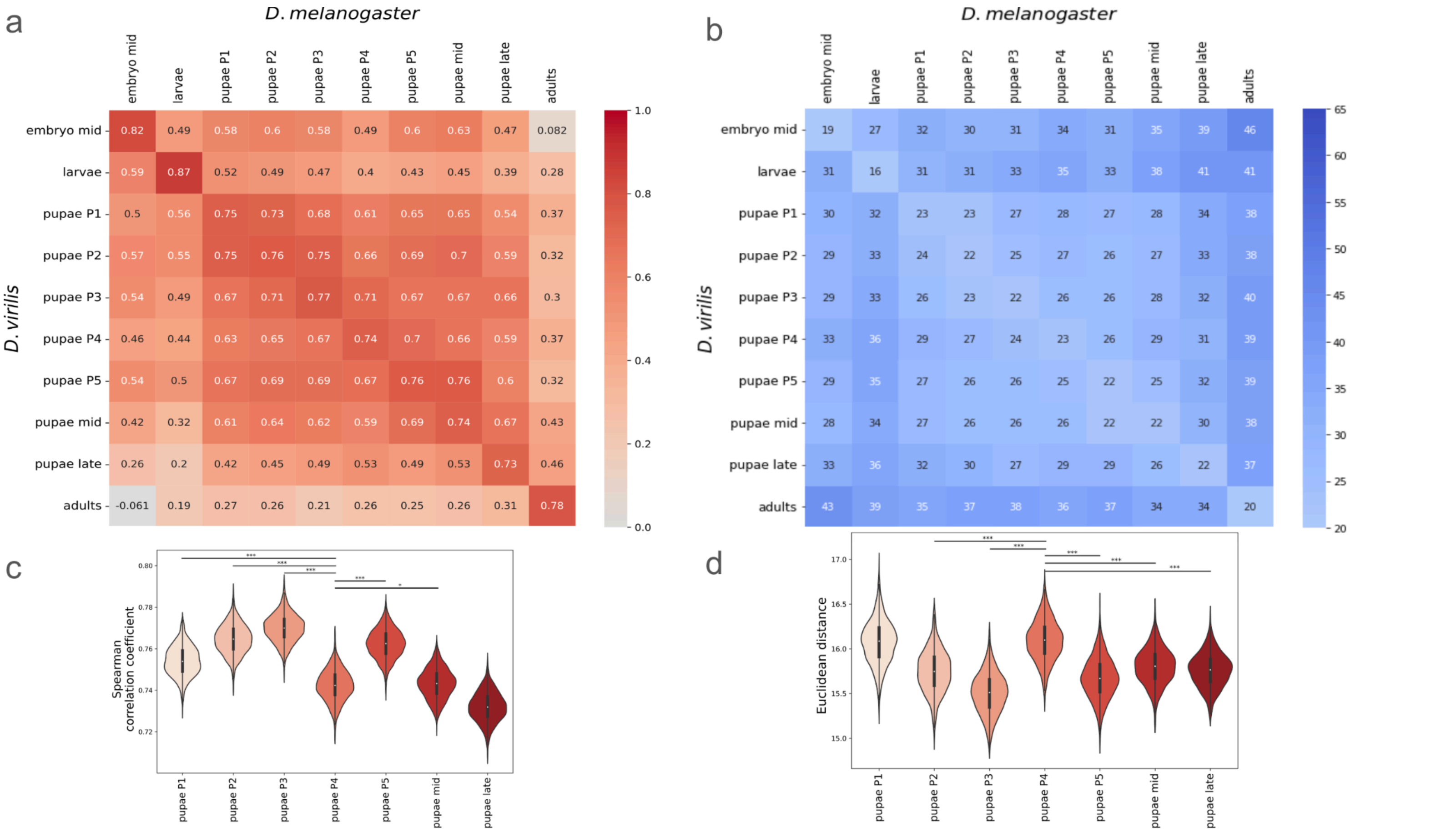


**Supplementary Fig. 6.** Distances between *D. melanogaster* and *D. virilis* developmental stages.

**(a)** Heatmap of the Spearman correlation coefficients. Each square represents the correlation coefficient between the expression profiles of the corresponding stages between the two species. The x-axis lists the stages of *D. melanogaster*, while the y-axis lists the stages of *D. virilis*. The color ranges from grey for low correlation coefficients (close to zero) to dark red for coefficients close to 1, with more intense colors indicating higher correlation coefficients.

**(b)** Heatmap of the Euclidean distances. Each square represents the Euclidean distance between the respective stages of development. The color ranges from light blue for small distances (closer to zero) to dark blue for larger ones, with more intense colors indicating larger distance. The Euclidean distance was calculated using log-transformed expression values after the quantile normalization between species to avoid the batch effect.

**(c and d)** Violin plots showing the distribution of correlation coefficients **(c)** and the Euclidean distances **(d)** between *D. melanogaster* and *D. virilis* across pupal stages. Correlation coefficients and the Euclidean distances were calculated using a random subset of genes (50% of the dataset) across 1000 iterations. Statistical significance of differences between adjacent stages was assessed using the one-sided Wilcoxon-Mann-Whitney test.

Significant differences are indicated with asterisks: * ($p < 0.05$), ** ($p < 0.01$), *** ($p < 0.001$), and **** ($p < 0.0001$).

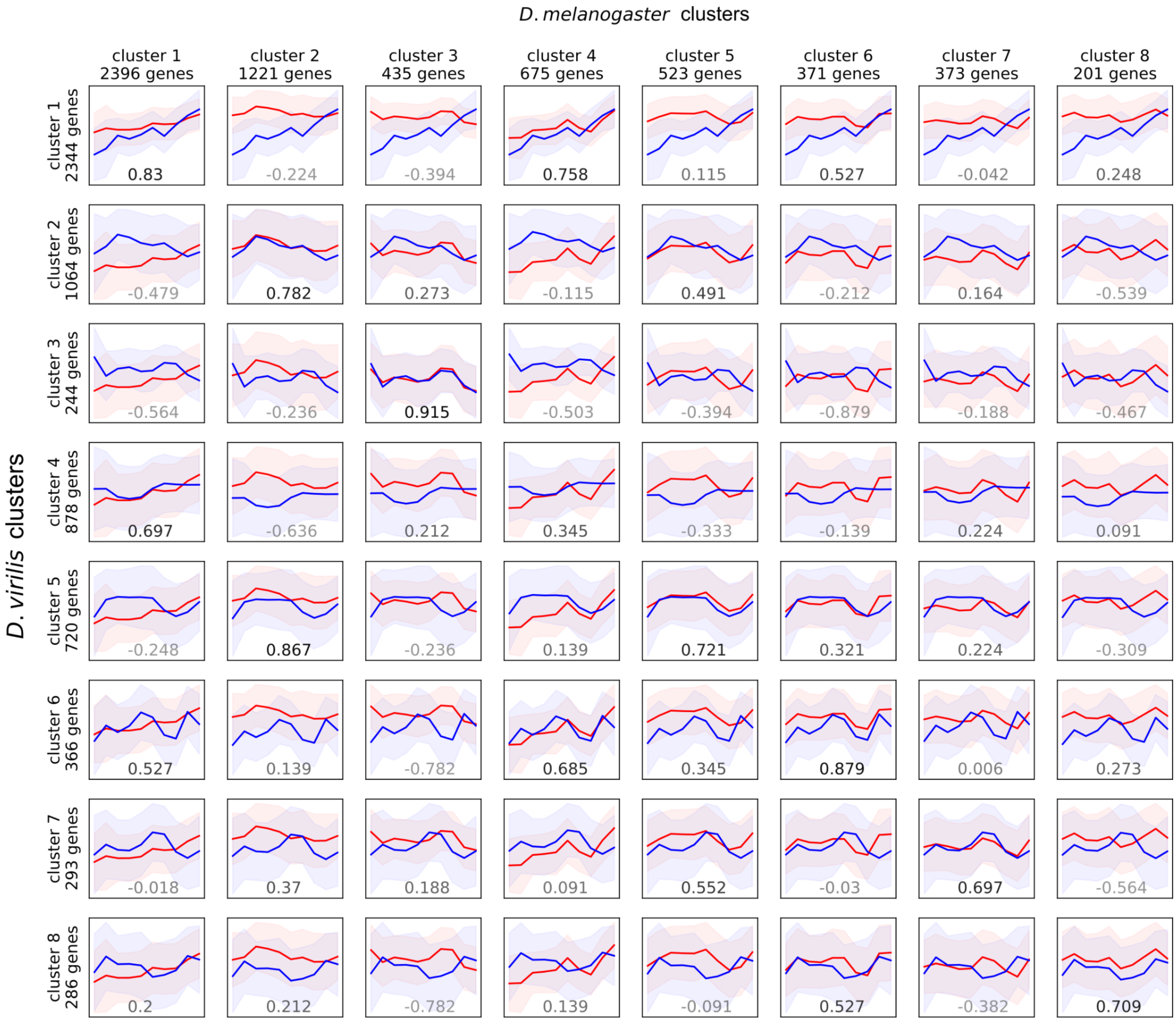


**Supplementary Fig 7.** Similarity of clusters between species. Each plot represents median profiles with the standard deviation displayed as a partially transparent area around the main curves. The median profiles are shown in red for *D. melanogaster* and blue for *D. virilis*. The number displayed on each plot represents the Spearman correlation coefficient between the corresponding median profiles. The x-axis represents *D. melanogaster* clusters, while the y-axis represents *D. virilis* clusters.

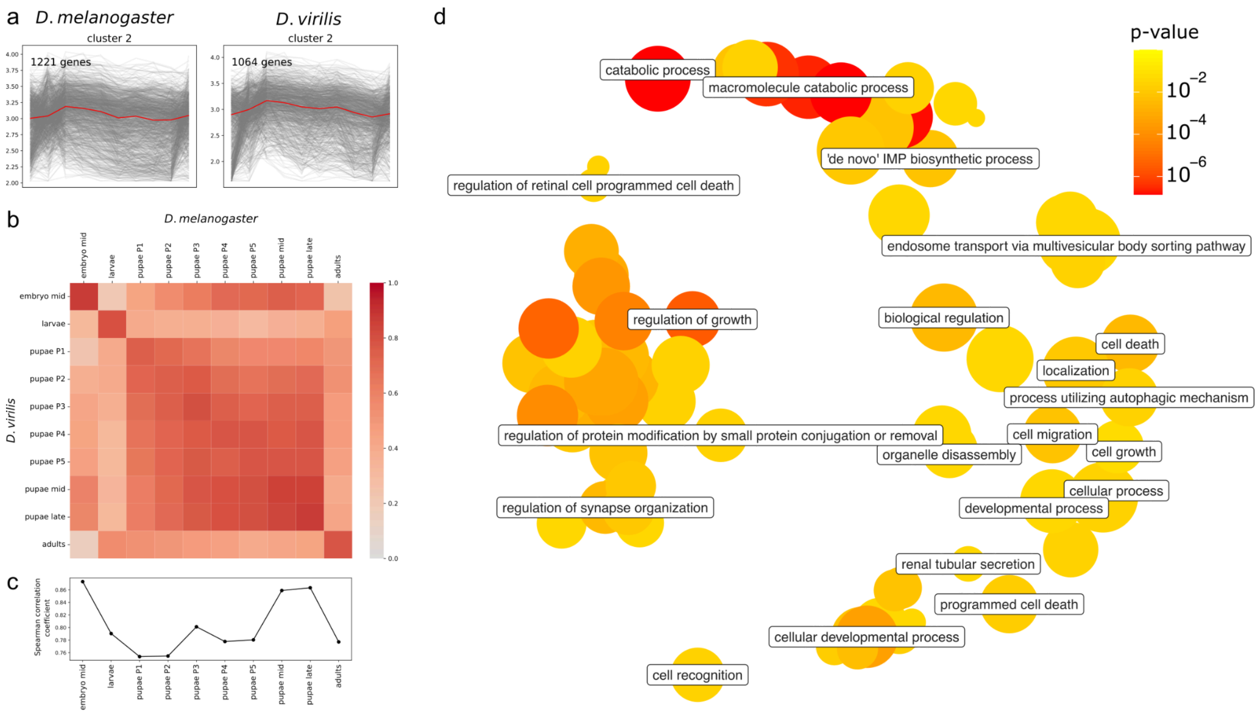


**Supplementary Fig 8**. Analysis of clusters 2 of *D. melanogaster* and *D. virilis*, containing 519 intersecting genes. **(a)** Dynamics of genes comprising the selected clusters for *D. melanogaster* (left) and *D. virilis* (right), the median curve is shown in red. **(b)** A pairwise correlation heatmap comparing the expression profiles of all stages of development. **(c)** Change in the Spearman correlation coefficient for the main diagonal of the heatmap, specifically comparing the corresponding stages in each species. **(d)** Gene ontology (GO) terms enriched in the common gene set, shown in the semantic space, where terms that are close semantically are also close in the plot. The size of each bubble represents the number of annotations for GO Term ID; the color, FDR-corrected enrichment *p*-value. Adapted from the `Revigo` tool, see the *Materials and Methods* section for details.

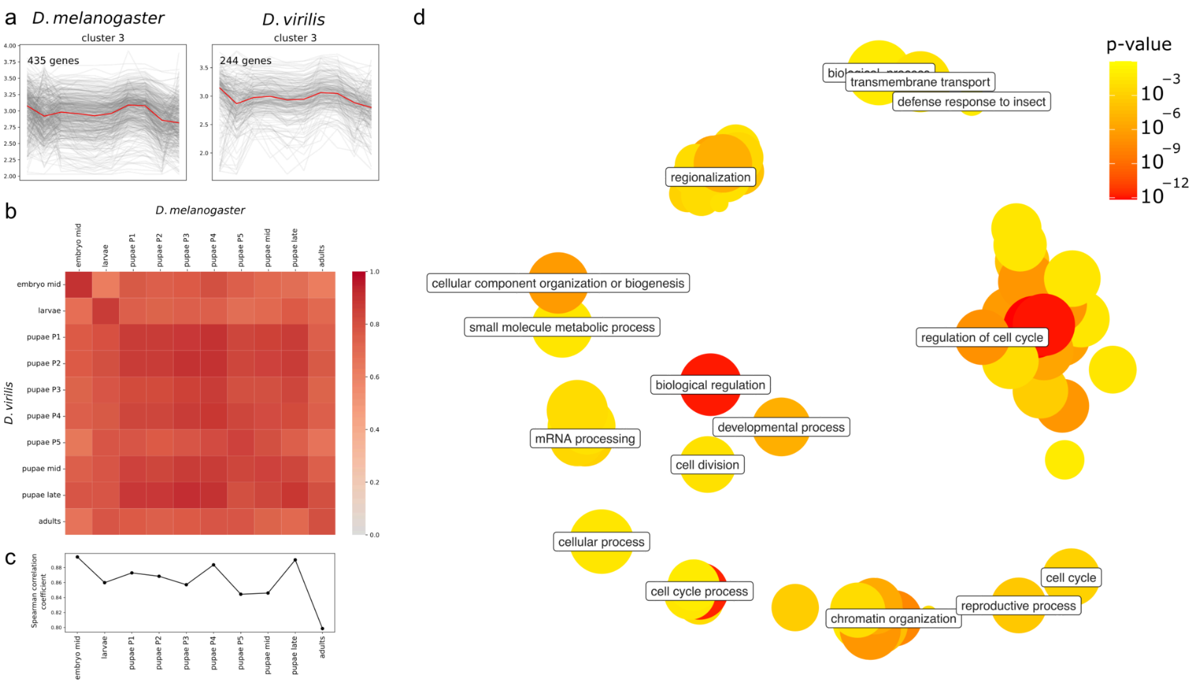


**Supplementary Fig 9**. Analysis of clusters 3 of *D. melanogaster* and *D. virilis*, containing 112 intersecting genes. Notation as in Supplementary Fig. 6.

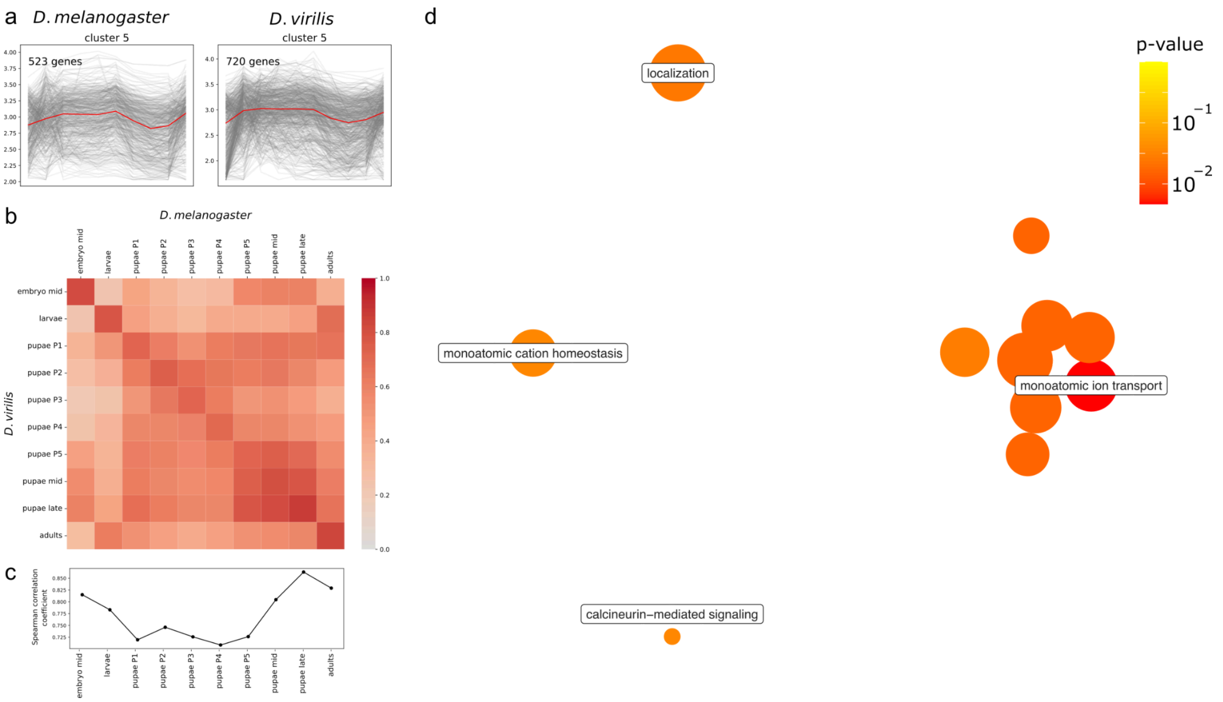


**Supplementary Fig 10**. Analysis of clusters 5 of *D. melanogaster* and *D. virilis*, containing 161 intersecting genes. Notation as in Supplementary Fig. 6.

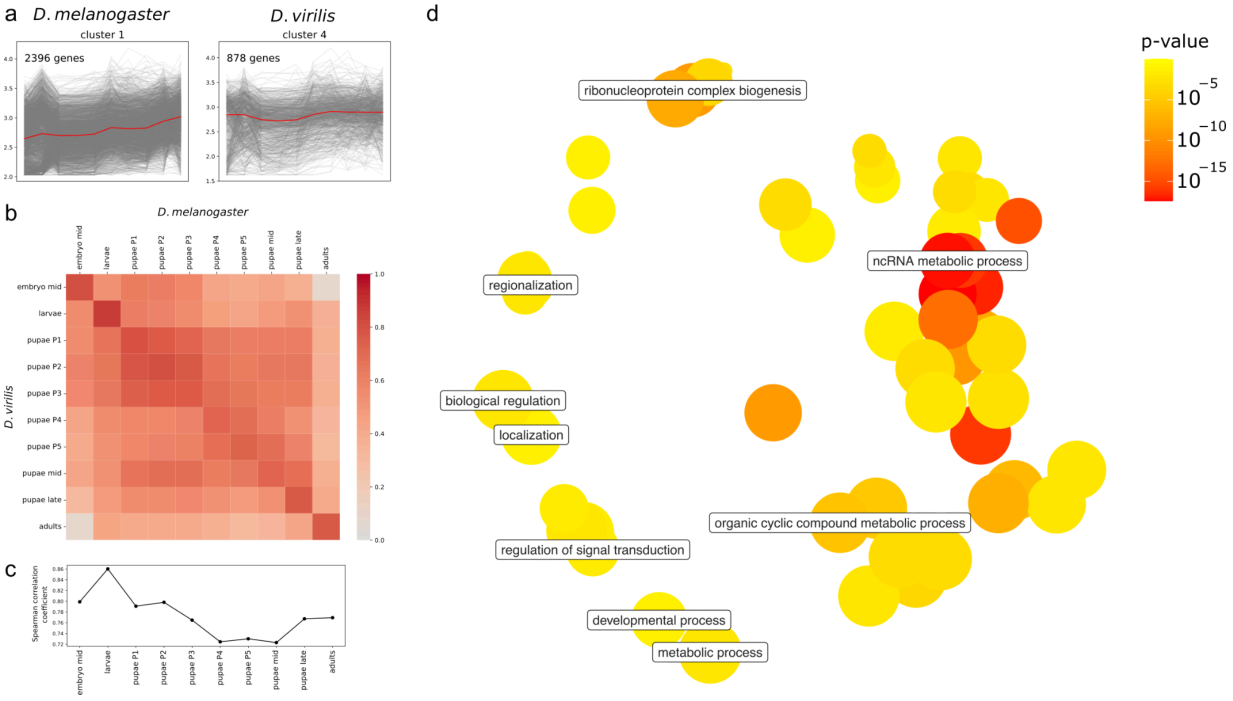


**Supplementary Fig 11**. Analysis of clusters of cluster 1 of *D. melanogaster* and cluster 4 of *D. virilis*, containing 495 intersecting genes. Notation as in Supplementary Fig. 6.

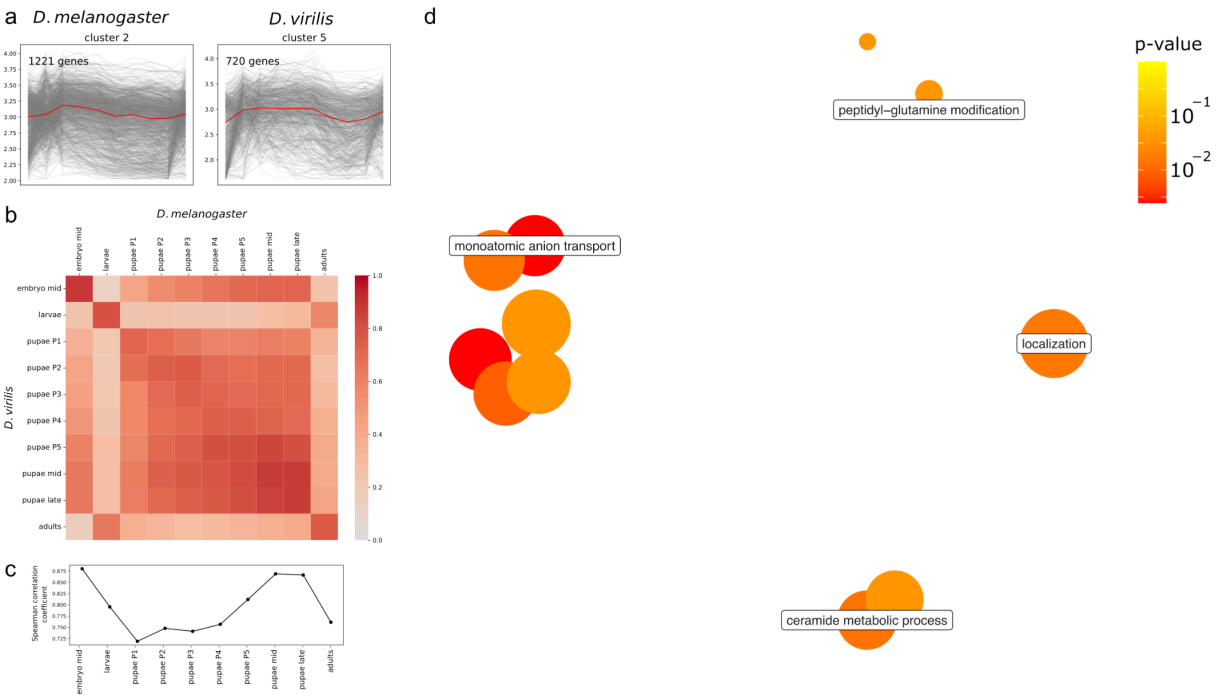


**Supplementary Fig 12**. Analysis of clusters of cluster 2 of *D. melanogaster* and cluster 5 of *D. virilis*, containing 255 intersecting genes. Notation as in Supplementary Fig. 6.

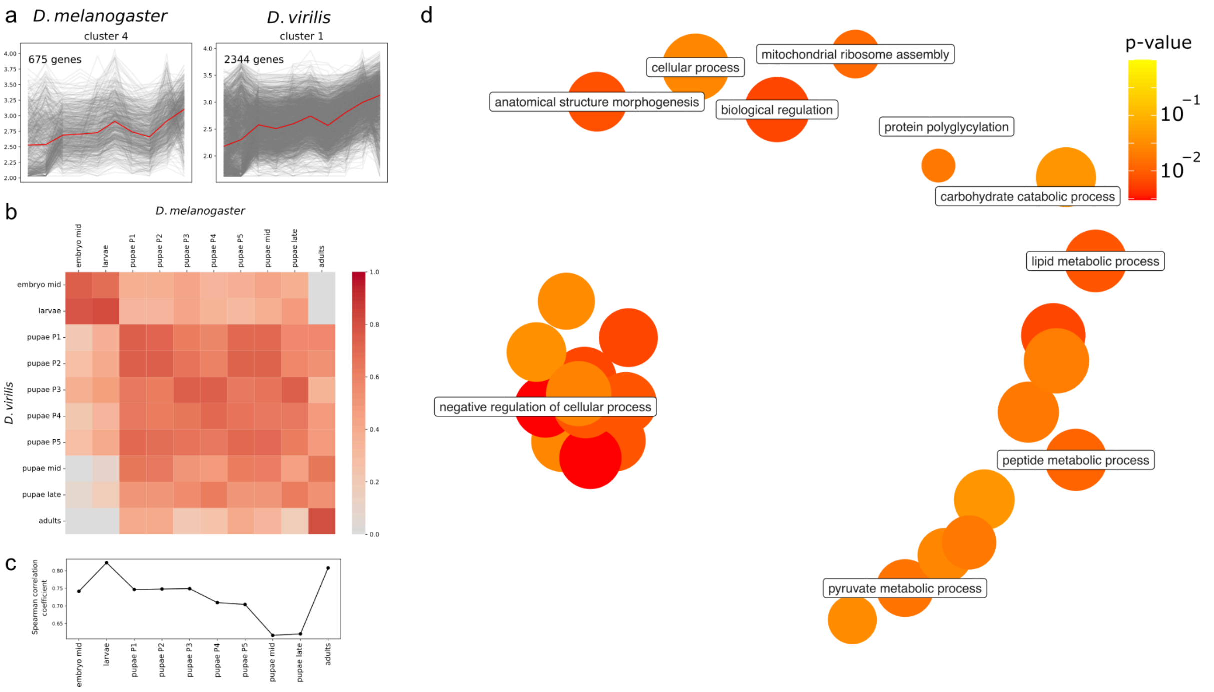


**Supplementary Fig 13**. Analysis of clusters of cluster 4 of *D. melanogaster* and cluster 1 of *D. virilis*, containing 443 intersecting genes. Notation as in Supplementary Fig. 6.

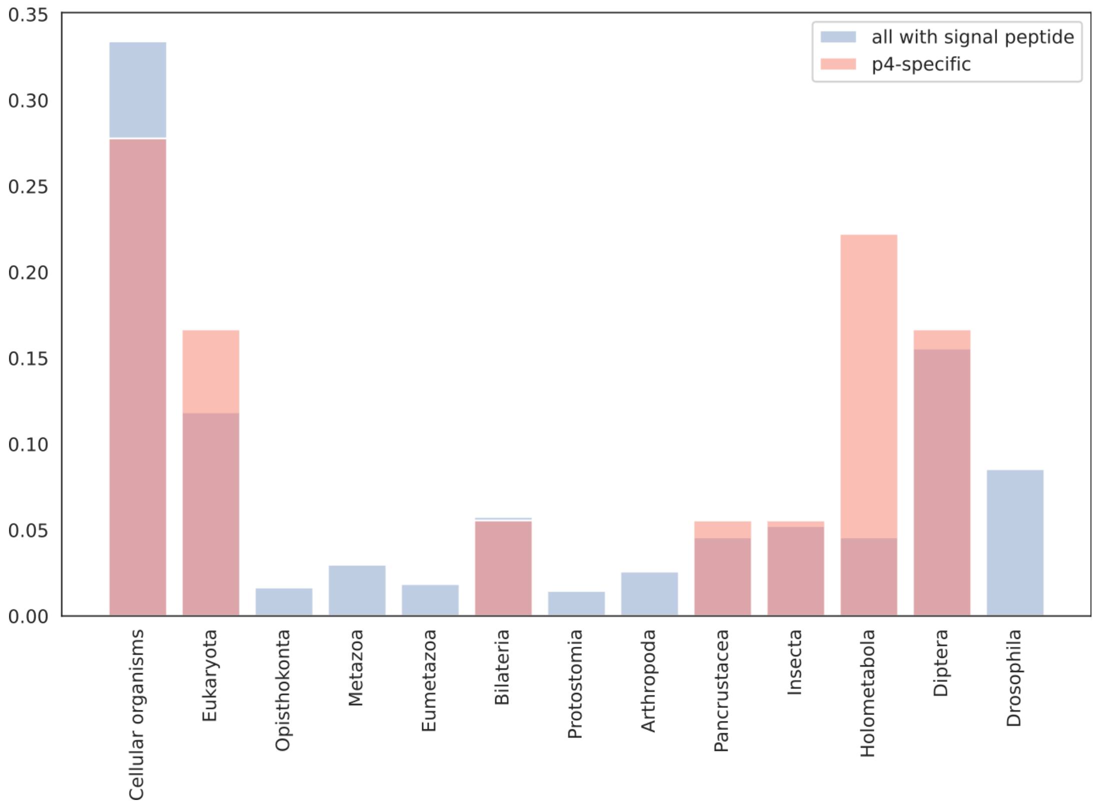


**Supplementary Fig. 14.** The fraction of genes assigned to phylostrata (along the x-axis). The y-axis represents the proportion of genes within each phylostratum. The distribution of all genes is shown in blue, while the distribution of genes encoding proteins with signal peptides is shown in red.

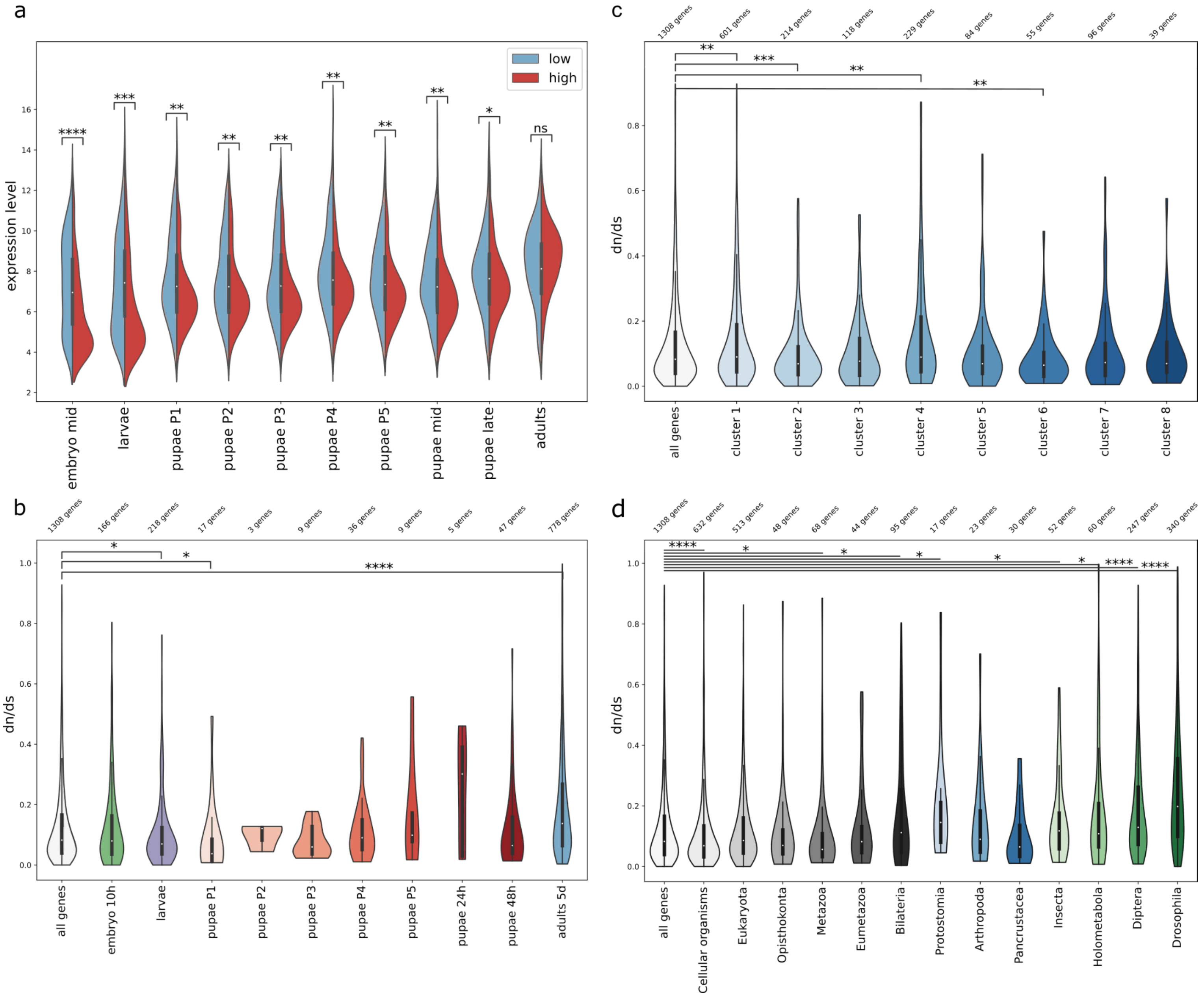


**Supplementary Fig. 15.** Distribution of the *dN/dS* ratio. **(a)** Comparison of the expression level between genes with the lowest and highest 10% *dN/dS* ratio at each stage of the development. **(b)** Distribution of the *dN/dS* ratio for genes up-regulated at the corresponding stage. **(c)** Distribution of the *dN/dS* ratio for genes comprising identified clusters (see Figure 3). **(d)** Distribution of the dN/dS ratio for genes from different phylostrata. Statistical significance was assessed using the one-sided Wilcoxon-Mann-Whitney test, and p-values were corrected for multiple testing using the Benjamini-Hochberg false discovery rate procedure (implemented in the `multipletests` function from the `statsmodels` Python package. Significant differences are indicated with asterisks: * ($p_{adj}$ < 0.05), ** ($p_{adj}$ < 0.01), *** ($p_{adj}$ < 0.001), and **** ($p_{adj}$ < 0.0001).
