## Supplementary Tables 1-6 for "Temporal dynamics of gene expression during metamorphosis in two distant *Drosophila* species"

| GO | name | GO type | p-value FDR *D. melanogaster* | p-value FDR *D. virilis* |
|---|---|---|---|---|
| GO:0003677 | DNA binding | molecular function | 3.09E-101 | 1.22E-56 |
| GO:0003676 | nucleic acid binding | molecular function | 3.75E-81 | 7.39E-69 |
| GO:0051252 | regulation of RNA metabolic process | biological process | 6.14E-90 | 6.56E-37 |
| GO:0019219 | regulation of nucleobase-containing compound metabolic process | biological process | 1.23E-89 | 6.56E-37 |
| GO:2000112 | regulation of cellular macromolecule biosynthetic process | biological process | 1.00E-87 | 6.56E-37 |
| GO:0010556 | regulation of macromolecule biosynthetic process | biological process | 2.00E-87 | 6.56E-37 |
| GO:0006355 | regulation of transcription, DNA-templated | biological process | 2.47E-86 | 6.56E-37 |
| GO:1903506 | regulation of nucleic acid-templated transcription | biological process | 2.47E-86 | 6.56E-37 |
| GO:2001141 | regulation of RNA biosynthetic process | biological process | 2.47E-86 | 6.56E-37 |
| GO:0031326 | regulation of cellular biosynthetic process | biological process | 3.98E-84 | 6.56E-37 |
| GO:0009889 | regulation of biosynthetic process | biological process | 1.59E-83 | 6.56E-37 |
| GO:0010468 | regulation of gene expression | biological process | 8.05E-85 | 2.36E-34 |
| GO:0060255 | regulation of macromolecule metabolic process | biological process | 1.26E-80 | 1.10E-34 |
| GO:0051171 | regulation of nitrogen compound metabolic process | biological process | 4.66E-79 | 2.76E-35 |
| GO:0080090 | regulation of primary metabolic process | biological process | 1.20E-76 | 2.76E-35 |
| GO:0031323 | regulation of cellular metabolic process | biological process | 4.05E-76 | 1.18E-35 |
| GO:0043565 | sequence-specific DNA binding | molecular function | 6.71E-81 | 1.77E-28 |
| GO:1901363 | heterocyclic compound binding | molecular function | 5.25E-55 | 3.66E-53 |
| GO:0019222 | regulation of metabolic process | biological process | 2.03E-74 | 1.06E-33 |
| GO:0097159 | organic cyclic compound binding | molecular function | 3.04E-54 | 3.66E-53 |
| GO:0050794 | regulation of cellular process | biological process | 5.79E-71 | 9.07E-30 |
| GO:0140110 | transcription regulator activity | molecular function | 1.17E-81 | 1.92E-16 |
| GO:0050789 | regulation of biological process | biological process | 2.58E-69 | 1.54E-28 |
| GO:0005488 | binding | molecular function | 1.19E-45 | 6.04E-51 |
| GO:0003700 | DNA-binding transcription factor activity | molecular function | 6.13E-78 | 2.13E-17 |
| GO:0065007 | biological regulation | biological process | 1.19E-61 | 2.02E-27 |
| GO:0005634 | nucleus | cellular component | 3.71E-62 | 9.95E-27 |
| GO:0032502 | developmental process | biological process | 1.03E-81 | 0.01191084257 |
| GO:0048856 | anatomical structure development | biological process | 1.53E-72 | 0.009876830148 |
| GO:0098772 | molecular function regulator | molecular function | 3.69E-59 | 1.15E-09 |
| GO:0005515 | protein binding | molecular function | 7.40E-36 | 5.23E-17 |
| GO:0046983 | protein dimerization activity | molecular function | 6.90E-32 | 9.78E-15 |

| | | | | |
|---|---|---|---|---|
| GO:0043231 | intracellular membrane-bounded organelle | cellular component | 3.39E-26 | 2.96E-20 |
| GO:0043227 | membrane-bounded organelle | cellular component | 8.82E-25 | 3.43E-20 |
| GO:0043229 | intracellular organelle | cellular component | 2.58E-29 | 1.86E-15 |
| GO:0043226 | organelle | cellular component | 3.70E-29 | 1.86E-15 |
| GO:0046982 | protein heterodimerization activity | molecular function | 9.60E-23 | 1.07E-07 |
| GO:0032993 | protein-DNA complex | cellular component | 2.27E-20 | 1.54E-09 |
| GO:0044815 | DNA packaging complex | cellular component | 8.97E-20 | 3.92E-10 |
| GO:0006325 | chromatin organization | biological process | 8.24E-24 | 1.51E-05 |
| GO:0000786 | nucleosome | cellular component | 4.19E-18 | 1.40E-09 |
| GO:0016043 | cellular component organization | biological process | 7.68E-24 | 4.02E-03 |
| GO:0071840 | cellular component organization or biogenesis | biological process | 5.92E-24 | 9.94E-03 |
| GO:0003682 | chromatin binding | molecular function | 2.65E-22 | 2.32E-02 |
| GO:0034728 | nucleosome organization | biological process | 5.12E-17 | 8.97E-06 |
| GO:0006334 | nucleosome assembly | biological process | 3.32E-16 | 8.97E-06 |
| GO:0071824 | protein-DNA complex subunit organization | biological process | 8.96E-15 | 8.97E-06 |
| GO:0065004 | protein-DNA complex assembly | biological process | 3.79E-14 | 8.97E-06 |
| GO:0003674 | molecular_function | molecular function | 1.78E-03 | 4.55E-14 |
| GO:0005694 | chromosome | cellular component | 4.38E-11 | 1.49E-03 |
| GO:0022402 | cell cycle process | biological process | 2.39E-11 | 0.009955903567 |
| GO:0007166 | cell surface receptor signaling pathway | biological process | 3.74E-09 | 7.96E-05 |
| GO:0006270 | DNA replication initiation | biological process | 1.94E-05 | 2.28E-08 |
| GO:0051276 | chromosome organization | biological process | 2.46E-09 | 3.23E-04 |
| GO:0006260 | DNA replication | biological process | 2.14E-05 | 8.16E-08 |
| GO:0042555 | MCM complex | cellular component | 5.46E-05 | 3.01E-06 |
| GO:0006259 | DNA metabolic process | biological process | 9.74E-05 | 8.97E-06 |
| GO:0022607 | cellular component assembly | biological process | 2.81E-07 | 1.42E-02 |
| GO:0007275 | multicellular organism development | biological process | 2.38E-06 | 3.00E-02 |
| GO:0007167 | enzyme linked receptor protein signaling pathway | biological process | 4.75E-06 | 3.93E-02 |
| GO:0034622 | cellular protein-containing complex assembly | biological process | 2.10E-03 | 8.15E-04 |
| GO:0071103 | DNA conformation change | biological process | 1.66E-03 | 2.05E-03 |
| GO:0090304 | nucleic acid metabolic process | biological process | 9.85E-04 | 6.05E-03 |
| GO:0004713 | protein tyrosine kinase activity | molecular function | 1.53E-03 | 5.14E-03 |
| GO:0043933 | protein-containing complex subunit organization | biological process | 2.62E-03 | 6.04E-03 |

| | | | | |
|---|---|---|---|---|
| GO:0065003 | protein-containing complex assembly | biological process | 6.10E-03 | 0.002620579673 |
| GO:0007165 | signal transduction | biological process | 0.04665196958 | 0.006072749481 |
| GO:0006357 | regulation of transcription by RNA polymerase II | biological process | 4.88E-86 | |
| GO:0000981 | DNA-binding transcription factor activity, RNA polymerase II-specific | molecular function | 9.64E-77 | |
| GO:1990837 | sequence-specific double-stranded DNA binding | molecular function | 3.48E-76 | |
| GO:0048869 | cellular developmental process | biological process | 9.76E-76 | |
| GO:0000976 | transcription regulatory region sequence-specific DNA binding | molecular function | 5.36E-73 | |
| GO:0001067 | regulatory region nucleic acid binding | molecular function | 6.52E-73 | |
| GO:0003690 | double-stranded DNA binding | molecular function | 1.55E-72 | |
| GO:0000977 | RNA polymerase II transcription regulatory region sequence-specific DNA | molecular function | 1.56E-71 | |
| GO:0009653 | anatomical structure morphogenesis | biological process | 8.20E-69 | |
| GO:0048513 | animal organ development | biological process | 4.06E-67 | |
| GO:0010558 | negative regulation of macromolecule biosynthetic process | biological process | 2.73E-61 | |
| GO:2000113 | negative regulation of cellular macromolecule biosynthetic process | biological process | 2.73E-61 | |
| GO:0031327 | negative regulation of cellular biosynthetic process | biological process | 7.68E-60 | |
| GO:0009890 | negative regulation of biosynthetic process | biological process | 4.02E-59 | |
| GO:0048523 | negative regulation of cellular process | biological process | 1.55E-58 | |
| GO:0048519 | negative regulation of biological process | biological process | 1.56E-57 | |
| GO:0000978 | RNA polymerase II cis-regulatory region sequence-specific DNA binding | molecular function | 5.66E-56 | |
| GO:0000987 | cis-regulatory region sequence-specific DNA binding | molecular function | 6.52E-55 | |
| GO:0010629 | negative regulation of gene expression | biological process | 7.14E-55 | |
| GO:0051253 | negative regulation of RNA metabolic process | biological process | 4.10E-54 | |
| GO:0045892 | negative regulation of transcription, DNA-templated | biological process | 2.75E-53 | |
| GO:1902679 | negative regulation of RNA biosynthetic process | biological process | 2.75E-53 | |
| GO:1903507 | negative regulation of nucleic acid-templated transcription | biological process | 2.75E-53 | |
| GO:0045934 | negative regulation of nucleobase-containing compound metabolic process | biological process | 8.27E-53 | |
| GO:0010605 | negative regulation of macromolecule metabolic process | biological process | 6.57E-50 | |
| GO:0007389 | pattern specification process | biological process | 2.82E-47 | |
| GO:0009892 | negative regulation of metabolic process | biological process | 1.15E-46 | |
| GO:0051172 | negative regulation of nitrogen compound metabolic process | biological process | 1.43E-46 | |
| GO:0031324 | negative regulation of cellular metabolic process | biological process | 1.85E-45 | |
| GO:0030154 | cell differentiation | biological process | 8.80E-45 | |
| GO:0000122 | negative regulation of transcription by RNA polymerase II | biological process | 8.88E-44 | |

| | | | |
|---|---|---|---|
| GO:0051254 | positive regulation of RNA metabolic process | biological process | 3.96E-38 |
| GO:0003002 | regionalization | biological process | 6.77E-37 |
| GO:0045893 | positive regulation of transcription, DNA-templated | biological process | 6.77E-37 |
| GO:1902680 | positive regulation of RNA biosynthetic process | biological process | 6.77E-37 |
| GO:1903508 | positive regulation of nucleic acid-templated transcription | biological process | 6.77E-37 |
| GO:0050793 | regulation of developmental process | biological process | 8.18E-37 |
| GO:0045935 | positive regulation of nucleobase-containing compound metabolic process | biological process | 1.80E-36 |
| GO:0010628 | positive regulation of gene expression | biological process | 2.90E-36 |
| GO:0048522 | positive regulation of cellular process | biological process | 5.10E-36 |
| GO:0048731 | system development | biological process | 8.94E-35 |
| GO:0010557 | positive regulation of macromolecule biosynthetic process | biological process | 3.63E-34 |
| GO:0048518 | positive regulation of biological process | biological process | 3.81E-34 |
| GO:0045595 | regulation of cell differentiation | biological process | 7.42E-34 |
| GO:0045944 | positive regulation of transcription by RNA polymerase II | biological process | 3.88E-33 |
| GO:0010604 | positive regulation of macromolecule metabolic process | biological process | 6.22E-32 |
| GO:0009887 | animal organ morphogenesis | biological process | 1.11E-31 |
| GO:2000026 | regulation of multicellular organismal development | biological process | 1.20E-31 |
| GO:0051239 | regulation of multicellular organismal process | biological process | 1.40E-31 |
| GO:0006928 | movement of cell or subcellular component | biological process | 1.92E-31 |
| GO:0009891 | positive regulation of biosynthetic process | biological process | 2.51E-31 |
| GO:0031328 | positive regulation of cellular biosynthetic process | biological process | 2.51E-31 |
| GO:0097485 | neuron projection guidance | biological process | 7.22E-30 |
| GO:0009893 | positive regulation of metabolic process | biological process | 1.16E-29 |
| GO:0051173 | positive regulation of nitrogen compound metabolic process | biological process | 6.67E-29 |
| GO:0007411 | axon guidance | biological process | 2.53E-28 |
| GO:0031325 | positive regulation of cellular metabolic process | biological process | 3.61E-28 |
| GO:0050767 | regulation of neurogenesis | biological process | 7.67E-28 |
| GO:0060284 | regulation of cell development | biological process | 4.09E-27 |
| GO:0001709 | cell fate determination | biological process | 4.25E-27 |
| GO:0009888 | tissue development | biological process | 2.06E-26 |
| GO:0051960 | regulation of nervous system development | biological process | 5.76E-26 |
| GO:0032501 | multicellular organismal process | biological process | 1.90E-25 |
| GO:0048468 | cell development | biological process | 4.29E-25 |

| | | | |
|---|---|---|---|
| GO:0009886 | post-embryonic animal morphogenesis | biological process | 2.01E-23 |
| GO:0045165 | cell fate commitment | biological process | 4.40E-23 |
| GO:0001708 | cell fate specification | biological process | 1.70E-21 |
| GO:0002009 | morphogenesis of an epithelium | biological process | 4.77E-21 |
| GO:0030182 | neuron differentiation | biological process | 4.77E-21 |
| GO:0022414 | reproductive process | biological process | 2.19E-20 |
| GO:0048729 | tissue morphogenesis | biological process | 3.50E-20 |
| GO:0035295 | tube development | biological process | 4.97E-20 |
| GO:0007423 | sensory organ development | biological process | 2.05E-19 |
| GO:0061061 | muscle structure development | biological process | 4.15E-17 |
| GO:0035107 | appendage morphogenesis | biological process | 4.56E-17 |
| GO:0035239 | tube morphogenesis | biological process | 4.95E-17 |
| GO:0007417 | central nervous system development | biological process | 6.07E-17 |
| GO:0007419 | ventral cord development | biological process | 1.71E-16 |
| GO:0045664 | regulation of neuron differentiation | biological process | 2.06E-16 |
| GO:0035114 | imaginal disc-derived appendage morphogenesis | biological process | 2.15E-16 |
| GO:0048859 | formation of anatomical boundary | biological process | 4.22E-16 |
| GO:0060429 | epithelium development | biological process | 6.32E-16 |
| GO:0048812 | neuron projection morphogenesis | biological process | 1.30E-15 |
| GO:0048858 | cell projection morphogenesis | biological process | 1.30E-15 |
| GO:0120039 | plasma membrane bounded cell projection morphogenesis | biological process | 1.30E-15 |
| GO:0032989 | cellular component morphogenesis | biological process | 1.49E-15 |
| GO:0032990 | cell part morphogenesis | biological process | 1.49E-15 |
| GO:0048867 | stem cell fate determination | biological process | 1.85E-15 |
| GO:0051093 | negative regulation of developmental process | biological process | 2.07E-15 |
| GO:0048598 | embryonic morphogenesis | biological process | 2.32E-15 |
| GO:0022412 | cellular process involved in reproduction in multicellular organism | biological process | 5.96E-15 |
| GO:0045596 | negative regulation of cell differentiation | biological process | 6.07E-15 |
| GO:0031491 | nucleosome binding | molecular function | 6.21E-15 |
| GO:0051128 | regulation of cellular component organization | biological process | 1.14E-14 |
| GO:0007400 | neuroblast fate determination | biological process | 1.27E-14 |
| GO:0022603 | regulation of anatomical structure morphogenesis | biological process | 1.37E-14 |
| GO:0048608 | reproductive structure development | biological process | 1.52E-14 |

| | | | | |
|---|---|---|---|---|
| GO:0009987 | cellular process | biological process | 1.79E-14 | |
| GO:0003006 | developmental process involved in reproduction | biological process | 2.44E-14 | |
| GO:0008134 | transcription factor binding | molecular function | 2.46E-14 | |
| GO:0007422 | peripheral nervous system development | biological process | 5.35E-14 | |
| GO:0031492 | nucleosomal DNA binding | molecular function | 5.61E-14 | |
| GO:0030030 | cell projection organization | biological process | 1.39E-13 | |
| GO:0060562 | epithelial tube morphogenesis | biological process | 1.60E-13 | |
| GO:0042127 | regulation of cell population proliferation | biological process | 2.64E-13 | |
| GO:0040011 | locomotion | biological process | 3.59E-13 | |
| GO:0008150 | biological_process | biological process | 3.77E-13 | |
| GO:0048592 | eye morphogenesis | biological process | 3.96E-13 | |
| GO:0090596 | sensory organ morphogenesis | biological process | 3.96E-13 | |
| GO:2000027 | regulation of animal organ morphogenesis | biological process | 3.96E-13 | |
| GO:0008406 | gonad development | biological process | 4.21E-13 | |
| GO:0007525 | somatic muscle development | biological process | 4.87E-13 | |
| GO:0035120 | post-embryonic appendage morphogenesis | biological process | 6.59E-13 | |
| GO:0043228 | non-membrane-bounded organelle | cellular component | 8.47E-13 | |
| GO:0043232 | intracellular non-membrane-bounded organelle | cellular component | 8.47E-13 | |
| GO:0016477 | cell migration | biological process | 1.14E-12 | |
| GO:0007447 | imaginal disc pattern formation | biological process | 1.39E-12 | |
| GO:0040012 | regulation of locomotion | biological process | 1.42E-12 | |
| GO:0051241 | negative regulation of multicellular organismal process | biological process | 1.53E-12 | |
| GO:0009952 | anterior/posterior pattern specification | biological process | 1.57E-12 | |
| GO:0051726 | regulation of cell cycle | biological process | 1.97E-12 | |
| GO:0007476 | imaginal disc-derived wing morphogenesis | biological process | 2.31E-12 | |
| GO:0043167 | ion binding | molecular function | | 2.94E-12 |
| GO:0007379 | segment specification | biological process | 3.30E-12 | |
| GO:0042659 | regulation of cell fate specification | biological process | 3.30E-12 | |
| GO:0048646 | anatomical structure formation involved in morphogenesis | biological process | 3.89E-12 | |
| GO:0010453 | regulation of cell fate commitment | biological process | 8.38E-12 | |
| GO:0010160 | formation of animal organ boundary | biological process | 1.03E-11 | |
| GO:0005700 | polytene chromosome | cellular component | 1.17E-11 | |
| GO:0007346 | regulation of mitotic cell cycle | biological process | 1.29E-11 | |

| | | | | |
|---|---|---|---|---|
| GO:0008037 | cell recognition | biological process | 2.42E-11 | |
| GO:0009880 | embryonic pattern specification | biological process | 2.94E-11 | |
| GO:0008038 | neuron recognition | biological process | 5.27E-11 | |
| GO:0048870 | cell motility | biological process | 5.42E-11 | |
| GO:0001216 | DNA-binding transcription activator activity | molecular function | 5.99E-11 | |
| GO:0110165 | cellular anatomical entity | cellular component | 6.85E-11 | |
| GO:0035282 | segmentation | biological process | 8.12E-11 | |
| GO:0048583 | regulation of response to stimulus | biological process | 9.21E-11 | |
| GO:0046872 | metal ion binding | molecular function | | 1.58E-10 |
| GO:0001228 | DNA-binding transcription activator activity, RNA polymerase II-specific | molecular function | 2.24E-10 | |
| GO:0007445 | determination of imaginal disc primordium | biological process | 2.64E-10 | |
| GO:0014019 | neuroblast development | biological process | 2.64E-10 | |
| GO:0043169 | cation binding | molecular function | | 4.29E-10 |
| GO:0051270 | regulation of cellular component movement | biological process | 4.88E-10 | |
| GO:0051094 | positive regulation of developmental process | biological process | 5.65E-10 | |
| GO:0048864 | stem cell development | biological process | 6.42E-10 | |
| GO:0007560 | imaginal disc morphogenesis | biological process | 7.67E-10 | |
| GO:0008045 | motor neuron axon guidance | biological process | 9.49E-10 | |
| GO:0008347 | glial cell migration | biological process | 1.33E-09 | |
| GO:0048813 | dendrite morphogenesis | biological process | 1.68E-09 | |
| GO:0001745 | compound eye morphogenesis | biological process | 1.95E-09 | |
| GO:0007420 | brain development | biological process | 2.13E-09 | |
| GO:0048563 | post-embryonic animal organ morphogenesis | biological process | 2.46E-09 | |
| GO:0005667 | transcription regulator complex | cellular component | 3.18E-09 | |
| GO:0051301 | cell division | biological process | 3.41E-09 | |
| GO:0035225 | determination of genital disc primordium | biological process | 3.74E-09 | |
| GO:0001654 | eye development | biological process | 5.19E-09 | |
| GO:0035287 | head segmentation | biological process | 5.53E-09 | |
| GO:1903047 | mitotic cell cycle process | biological process | 5.53E-09 | |
| GO:0050768 | negative regulation of neurogenesis | biological process | 5.53E-09 | |
| GO:0070983 | dendrite guidance | biological process | 6.05E-09 | |
| GO:0005575 | cellular_component | cellular component | 6.71E-09 | |
| GO:0010564 | regulation of cell cycle process | biological process | 9.39E-09 | |

| | | | |
|---|---|---|---|
| GO:0045597 | positive regulation of cell differentiation | biological process | 9.39E-09 |
| GO:0046532 | regulation of photoreceptor cell differentiation | biological process | 1.15E-08 |
| GO:0007380 | specification of segmental identity, head | biological process | 1.84E-08 |
| GO:0007432 | salivary gland boundary specification | biological process | 1.84E-08 |
| GO:0110116 | regulation of compound eye photoreceptor cell differentiation | biological process | 2.51E-08 |
| GO:0044877 | protein-containing complex binding | molecular function | 2.65E-08 |
| GO:0010720 | positive regulation of cell development | biological process | 3.06E-08 |
| GO:0031490 | chromatin DNA binding | molecular function | 3.21E-08 |
| GO:0051961 | negative regulation of nervous system development | biological process | 3.48E-08 |
| GO:0007610 | behavior | biological process | 3.81E-08 |
| GO:0016198 | axon choice point recognition | biological process | 3.89E-08 |
| GO:0003714 | transcription corepressor activity | molecular function | 3.94E-08 |
| GO:0007399 | nervous system development | biological process | 5.28E-08 |
| GO:0120035 | regulation of plasma membrane bounded cell projection organization | biological process | 5.28E-08 |
| GO:0040008 | regulation of growth | biological process | 5.38E-08 |
| GO:0010454 | negative regulation of cell fate commitment | biological process | 5.90E-08 |
| GO:0031344 | regulation of cell projection organization | biological process | 6.11E-08 |
| GO:0048732 | gland development | biological process | 7.19E-08 |
| GO:0098687 | chromosomal region | cellular component | 7.31E-08 |
| GO:0010975 | regulation of neuron projection development | biological process | 7.52E-08 |
| GO:0048666 | neuron development | biological process | 8.71E-08 |
| GO:0042051 | compound eye photoreceptor development | biological process | 8.94E-08 |
| GO:0007455 | eye-antennal disc morphogenesis | biological process | 9.50E-08 |
| GO:0044786 | cell cycle DNA replication | biological process | 9.50E-08 |
| GO:0001667 | ameboidal-type cell migration | biological process | 1.03E-07 |
| GO:0007526 | larval somatic muscle development | biological process | 1.05E-07 |
| GO:0048749 | compound eye development | biological process | 1.08E-07 |
| GO:0010721 | negative regulation of cell development | biological process | 1.08E-07 |
| GO:0048569 | post-embryonic animal organ development | biological process | 1.08E-07 |
| GO:0048585 | negative regulation of response to stimulus | biological process | 1.41E-07 |
| GO:0003712 | transcription coregulator activity | molecular function | 1.41E-07 |
| GO:0001227 | DNA-binding transcription repressor activity, RNA polymerase II-specific | molecular function | 1.47E-07 |
| GO:0030334 | regulation of cell migration | biological process | 1.63E-07 |

| | | | |
|---|---|---|---|
| GO:0035222 | wing disc pattern formation | biological process | 1.76E-07 |
| GO:0001217 | DNA-binding transcription repressor activity | molecular function | 1.85E-07 |
| GO:0042393 | histone binding | molecular function | 1.85E-07 |
| GO:0042803 | protein homodimerization activity | molecular function | 1.85E-07 |
| GO:0050839 | cell adhesion molecule binding | molecular function | 2.21E-07 |
| GO:0040017 | positive regulation of locomotion | biological process | 2.23E-07 |
| GO:0035277 | spiracle morphogenesis, open tracheal system | biological process | 2.26E-07 |
| GO:0051240 | positive regulation of multicellular organismal process | biological process | 2.60E-07 |
| GO:0007169 | transmembrane receptor protein tyrosine kinase signaling pathway | biological process | 2.68E-07 |
| GO:0042462 | eye photoreceptor cell development | biological process | 2.75E-07 |
| GO:0043549 | regulation of kinase activity | biological process | 3.10E-07 |
| GO:0007444 | imaginal disc development | biological process | 3.16E-07 |
| GO:0045665 | negative regulation of neuron differentiation | biological process | 3.61E-07 |
| GO:2000145 | regulation of cell motility | biological process | 3.80E-07 |
| GO:0008283 | cell population proliferation | biological process | 4.73E-07 |
| GO:0009996 | negative regulation of cell fate specification | biological process | 4.73E-07 |
| GO:0051338 | regulation of transferase activity | biological process | 4.84E-07 |
| GO:0042023 | DNA endoreduplication | biological process | 5.07E-07 |
| GO:0007507 | heart development | biological process | 5.44E-07 |
| GO:0007517 | muscle organ development | biological process | 5.44E-07 |
| GO:0000785 | chromatin | cellular component | 6.28E-07 |
| GO:0008356 | asymmetric cell division | biological process | 6.98E-07 |
| GO:0035289 | posterior head segmentation | biological process | 7.12E-07 |
| GO:0050673 | epithelial cell proliferation | biological process | 7.12E-07 |
| GO:0061331 | epithelial cell proliferation involved in Malpighian tubule morphogenesis | biological process | 7.12E-07 |
| GO:2001013 | epithelial cell proliferation involved in renal tubule morphogenesis | biological process | 7.12E-07 |
| GO:0016318 | ommatidial rotation | biological process | 7.41E-07 |
| GO:2000177 | regulation of neural precursor cell proliferation | biological process | 7.72E-07 |
| GO:0008284 | positive regulation of cell population proliferation | biological process | 7.93E-07 |
| GO:0010631 | epithelial cell migration | biological process | 8.86E-07 |
| GO:0042461 | photoreceptor cell development | biological process | 9.17E-07 |
| GO:0007059 | chromosome segregation | biological process | 1.00E-06 |
| GO:0007409 | axonogenesis | biological process | 1.00E-06 |

| | | | | |
|---|---|---|---|---|
| GO:0009790 | embryo development | biological process | 1.01E-06 | |
| GO:0009792 | embryo development ending in birth or egg hatching | biological process | 1.01E-06 | |
| GO:0098813 | nuclear chromosome segregation | biological process | 1.01E-06 | |
| GO:0046533 | negative regulation of photoreceptor cell differentiation | biological process | 1.02E-06 | |
| GO:0048638 | regulation of developmental growth | biological process | 1.11E-06 | |
| GO:0007424 | open tracheal system development | biological process | 1.16E-06 | |
| GO:0060541 | respiratory system development | biological process | 1.16E-06 | |
| GO:0032991 | protein-containing complex | cellular component | 1.57E-06 | |
| GO:0007435 | salivary gland morphogenesis | biological process | 1.63E-06 | |
| GO:0022612 | gland morphogenesis | biological process | 1.63E-06 | |
| GO:0090175 | regulation of establishment of planar polarity | biological process | 1.73E-06 | |
| GO:0009953 | dorsal/ventral pattern formation | biological process | 1.73E-06 | |
| GO:0035051 | cardiocyte differentiation | biological process | 1.86E-06 | |
| GO:0016199 | axon midline choice point recognition | biological process | 1.92E-06 | |
| GO:0045926 | negative regulation of growth | biological process | 1.98E-06 | |
| GO:0007498 | mesoderm development | biological process | 2.03E-06 | |
| GO:0048477 | oogenesis | biological process | 2.10E-06 | |
| GO:0008270 | zinc ion binding | molecular function | | 2.16E-06 |
| GO:0007292 | female gamete generation | biological process | 2.37E-06 | |
| GO:0007494 | midgut development | biological process | 2.60E-06 | |
| GO:0007450 | dorsal/ventral pattern formation, imaginal disc | biological process | 2.66E-06 | |
| GO:0098609 | cell-cell adhesion | biological process | 2.74E-06 | |
| GO:0017053 | transcription repressor complex | cellular component | 2.76E-06 | |
| GO:0030155 | regulation of cell adhesion | biological process | 3.47E-06 | |
| GO:0009966 | regulation of signal transduction | biological process | 3.58E-06 | |
| GO:0007155 | cell adhesion | biological process | 3.64E-06 | |
| GO:0007156 | homophilic cell adhesion via plasma membrane adhesion molecules | biological process | 3.77E-06 | |
| GO:0050769 | positive regulation of neurogenesis | biological process | 4.07E-06 | |
| GO:0006333 | chromatin assembly or disassembly | biological process | 4.09E-06 | |
| GO:0061318 | renal filtration cell differentiation | biological process | 4.26E-06 | |
| GO:0061319 | nephrocyte differentiation | biological process | 4.26E-06 | |
| GO:0051962 | positive regulation of nervous system development | biological process | 5.22E-06 | |
| GO:0006935 | chemotaxis | biological process | 5.52E-06 | |

| | | | | |
|---|---|---|---|---|
| GO:0022407 | regulation of cell-cell adhesion | biological process | 5.52E-06 | |
| GO:0050920 | regulation of chemotaxis | biological process | 5.52E-06 | |
| GO:0016331 | morphogenesis of embryonic epithelium | biological process | 6.73E-06 | |
| GO:0051272 | positive regulation of cellular component movement | biological process | 7.64E-06 | |
| GO:0042802 | identical protein binding | molecular function | 7.82E-06 | |
| GO:0030335 | positive regulation of cell migration | biological process | 9.19E-06 | |
| GO:0022610 | biological adhesion | biological process | 9.44E-06 | |
| GO:0046530 | photoreceptor cell differentiation | biological process | 9.44E-06 | |
| GO:0009798 | axis specification | biological process | 9.75E-06 | |
| GO:1905330 | regulation of morphogenesis of an epithelium | biological process | 1.07E-05 | |
| GO:0009948 | anterior/posterior axis specification | biological process | 1.21E-05 | |
| GO:0045611 | negative regulation of hemocyte differentiation | biological process | 1.29E-05 | |
| GO:0001558 | regulation of cell growth | biological process | 1.34E-05 | |
| GO:0009968 | negative regulation of signal transduction | biological process | 1.41E-05 | |
| GO:0006338 | chromatin remodeling | biological process | 1.52E-05 | |
| GO:0048565 | digestive tract development | biological process | 1.57E-05 | |
| GO:0005654 | nucleoplasm | cellular component | 1.61E-05 | |
| GO:0034329 | cell junction assembly | biological process | 1.74E-05 | |
| GO:0045467 | R7 cell development | biological process | 1.78E-05 | |
| GO:0007469 | antennal development | biological process | 1.80E-05 | |
| GO:0010001 | glial cell differentiation | biological process | 1.80E-05 | |
| GO:0045466 | R7 cell differentiation | biological process | 1.80E-05 | |
| GO:0110118 | negative regulation of compound eye photoreceptor cell differentiation | biological process | 1.80E-05 | |
| GO:0140097 | catalytic activity, acting on DNA | molecular function | | 1.83E-05 |
| GO:0098742 | cell-cell adhesion via plasma-membrane adhesion molecules | biological process | 1.90E-05 | |
| GO:0045610 | regulation of hemocyte differentiation | biological process | 1.94E-05 | |
| GO:0010648 | negative regulation of cell communication | biological process | 1.99E-05 | |
| GO:0070161 | anchoring junction | cellular component | 2.04E-05 | |
| GO:0023057 | negative regulation of signaling | biological process | 2.07E-05 | |
| GO:0061326 | renal tubule development | biological process | 2.09E-05 | |
| GO:0072002 | Malpighian tubule development | biological process | 2.09E-05 | |
| GO:2000147 | positive regulation of cell motility | biological process | 2.14E-05 | |
| GO:0007391 | dorsal closure | biological process | 2.41E-05 | |

| | | | | |
|---|---|---|---|---|
| GO:0010769 | regulation of cell morphogenesis involved in differentiation | biological process | 2.41E-05 | |
| GO:0010632 | regulation of epithelial cell migration | biological process | 2.48E-05 | |
| GO:0006468 | protein phosphorylation | biological process | | 2.83E-05 |
| GO:0022604 | regulation of cell morphogenesis | biological process | 2.91E-05 | |
| GO:0048190 | wing disc dorsal/ventral pattern formation | biological process | 3.18E-05 | |
| GO:1901987 | regulation of cell cycle phase transition | biological process | 3.32E-05 | |
| GO:0035050 | embryonic heart tube development | biological process | 3.35E-05 | |
| GO:0005524 | ATP binding | molecular function | | 3.35E-05 |
| GO:0051347 | positive regulation of transferase activity | biological process | 3.70E-05 | |
| GO:0050770 | regulation of axonogenesis | biological process | 3.97E-05 | |
| GO:0072499 | photoreceptor cell axon guidance | biological process | 4.21E-05 | |
| GO:0004672 | protein kinase activity | molecular function | | 4.40E-05 |
| GO:0003678 | DNA helicase activity | molecular function | | 4.56E-05 |
| GO:0003730 | mRNA 3'-UTR binding | molecular function | 4.92E-05 | |
| GO:0007474 | imaginal disc-derived wing vein specification | biological process | 5.09E-05 | |
| GO:0008407 | chaeta morphogenesis | biological process | 5.09E-05 | |
| GO:0006277 | DNA amplification | biological process | 5.24E-05 | |
| GO:0032559 | adenyl ribonucleotide binding | molecular function | | 5.75E-05 |
| GO:0017145 | stem cell division | biological process | 5.77E-05 | |
| GO:0001700 | embryonic development via the syncytial blastoderm | biological process | 5.85E-05 | |
| GO:0030554 | adenyl nucleotide binding | molecular function | | 5.86E-05 |
| GO:0034330 | cell junction organization | biological process | 6.03E-05 | |
| GO:0033674 | positive regulation of kinase activity | biological process | 6.75E-05 | |
| GO:0032879 | regulation of localization | biological process | 6.78E-05 | |
| GO:0045859 | regulation of protein kinase activity | biological process | 6.87E-05 | |
| GO:0002682 | regulation of immune system process | biological process | 7.36E-05 | |
| GO:0035153 | epithelial cell type specification, open tracheal system | biological process | 7.36E-05 | |
| GO:0007281 | germ cell development | biological process | 7.83E-05 | |
| GO:0009954 | proximal/distal pattern formation | biological process | 7.87E-05 | |
| GO:1901990 | regulation of mitotic cell cycle phase transition | biological process | 7.89E-05 | |
| GO:0045448 | mitotic cell cycle, embryonic | biological process | 8.49E-05 | |
| GO:0001751 | compound eye photoreceptor cell differentiation | biological process | 8.56E-05 | |
| GO:0045931 | positive regulation of mitotic cell cycle | biological process | 8.56E-05 | |

| | | | | |
|---|---|---|---|---|
| GO:1904949 | ATPase complex | cellular component | 8.96E-05 | |
| GO:0007297 | ovarian follicle cell migration | biological process | 9.23E-05 | |
| GO:0035288 | anterior head segmentation | biological process | 9.60E-05 | |
| GO:0051890 | regulation of cardioblast differentiation | biological process | 9.60E-05 | |
| GO:0060563 | neuroepithelial cell differentiation | biological process | 9.60E-05 | |
| GO:0061101 | neuroendocrine cell differentiation | biological process | 9.60E-05 | |
| GO:0070603 | SWI/SNF superfamily-type complex | cellular component | 9.61E-05 | |
| GO:0048534 | hematopoietic or lymphoid organ development | biological process | 9.97E-05 | |
| GO:0021782 | glial cell development | biological process | 0.0001032104796 | |
| GO:0008094 | DNA-dependent ATPase activity | molecular function | | 0.0001036942719 |
| GO:0032040 | small-subunit processome | cellular component | 0.0001080049527 | |
| GO:0040007 | growth | biological process | 0.0001080510832 | |
| GO:0048589 | developmental growth | biological process | 0.0001080510832 | |
| GO:0019904 | protein domain specific binding | molecular function | 0.000110373254 | |
| GO:0045676 | regulation of R7 cell differentiation | biological process | 0.0001136890076 | |
| GO:0072091 | regulation of stem cell proliferation | biological process | 0.0001160475399 | |
| GO:0045787 | positive regulation of cell cycle | biological process | 0.0001166486259 | |
| GO:0048640 | negative regulation of developmental growth | biological process | 0.0001166486259 | |
| GO:0000278 | mitotic cell cycle | biological process | 0.0001201088695 | |
| GO:0045746 | negative regulation of Notch signaling pathway | biological process | 0.0001350228394 | |
| GO:0017148 | negative regulation of translation | biological process | 0.0001414556701 | |
| GO:0140297 | DNA-binding transcription factor binding | molecular function | 0.0001546851924 | |
| GO:0000070 | mitotic sister chromatid segregation | biological process | 0.0001570517849 | |
| GO:0110111 | negative regulation of animal organ morphogenesis | biological process | 0.0001570517849 | |
| GO:1902667 | regulation of axon guidance | biological process | 0.0001570517849 | |
| GO:0007276 | gamete generation | biological process | 0.0001592014144 | |
| GO:0048865 | stem cell fate commitment | biological process | 0.0001592014144 | |
| GO:0034249 | negative regulation of cellular amide metabolic process | biological process | 0.00017082274 | |
| GO:0001754 | eye photoreceptor cell differentiation | biological process | 0.0001743122525 | |
| GO:1903311 | regulation of mRNA metabolic process | biological process | 0.0001769394859 | |
| GO:0007298 | border follicle cell migration | biological process | 0.0001801376904 | |
| GO:0042330 | taxis | biological process | 0.0001801376904 | |
| GO:0007611 | learning or memory | biological process | 0.0001826960645 | |

| | | | |
|---|---|---|---|
| GO:0010646 | regulation of cell communication | biological process | 0.0001844917594 |
| GO:0023051 | regulation of signaling | biological process | 0.0001844917594 |
| GO:0048663 | neuron fate commitment | biological process | 0.0001956862907 |
| GO:0050890 | cognition | biological process | 0.0002013519762 |
| GO:0007307 | eggshell chorion gene amplification | biological process | 0.0002032941725 |
| GO:0033301 | cell cycle comprising mitosis without cytokinesis | biological process | 0.0002032941725 |
| GO:0035146 | tube fusion | biological process | 0.0002032941725 |
| GO:0035147 | branch fusion, open tracheal system | biological process | 0.0002032941725 |
| GO:0043067 | regulation of programmed cell death | biological process | 0.0002032941725 |
| GO:0050684 | regulation of mRNA processing | biological process | 0.0002032941725 |
| GO:0000819 | sister chromatid segregation | biological process | 0.0002034776264 |
| GO:0055057 | neuroblast division | biological process | 0.0002034776264 |
| GO:0007049 | cell cycle | biological process | 0.0002054443601 |
| GO:0000727 | double-strand break repair via break-induced replication | biological process | 0.000216931291 |
| GO:0007522 | visceral muscle development | biological process | 0.000216931291 |
| GO:0048736 | appendage development | biological process | 0.000216931291 |
| GO:0006275 | regulation of DNA replication | biological process | 0.0002176545378 |
| GO:0048542 | lymph gland development | biological process | 0.0002192528104 |
| GO:1903046 | meiotic cell cycle process | biological process | 0.0002621971606 |
| GO:0043068 | positive regulation of programmed cell death | biological process | 0.0002662476291 |
| GO:2000736 | regulation of stem cell differentiation | biological process | 0.0002699848418 |
| GO:0007398 | ectoderm development | biological process | 0.0002772746886 |
| GO:1905207 | regulation of cardiocyte differentiation | biological process | 0.0002772746886 |
| GO:0007350 | blastoderm segmentation | biological process | 0.0002999612611 |
| GO:0007448 | anterior/posterior pattern specification, imaginal disc | biological process | 0.0003059928332 |
| GO:0046552 | photoreceptor cell fate commitment | biological process | 0.0003284003083 |
| GO:1902692 | regulation of neuroblast proliferation | biological process | 0.0003284003083 |
| GO:0030371 | translation repressor activity | molecular function | 0.0003299854227 |
| GO:0033043 | regulation of organelle organization | biological process | 0.0003471196508 |
| GO:0007416 | synapse assembly | biological process | 0.0003475628038 |
| GO:0090659 | walking behavior | biological process | 0.0003527236486 |
| GO:0008039 | synaptic target recognition | biological process | 0.0003530040092 |
| GO:0048609 | multicellular organismal reproductive process | biological process | 0.0003559047055 |

| | | | | |
|---|---|---|---|---|
| GO:0006261 | DNA-dependent DNA replication | biological process | 0.0004043806489 | |
| GO:0007365 | periodic partitioning | biological process | 0.0004043806489 | |
| GO:0046425 | regulation of receptor signaling pathway via JAK-STAT | biological process | 0.0004154291025 | |
| GO:1904892 | regulation of receptor signaling pathway via STAT | biological process | 0.0004154291025 | |
| GO:0016360 | sensory organ precursor cell fate determination | biological process | 0.0004300607968 | |
| GO:0050919 | negative chemotaxis | biological process | 0.0004300607968 | |
| GO:0060582 | cell fate determination involved in pattern specification | biological process | 0.0004300607968 | |
| GO:0051129 | negative regulation of cellular component organization | biological process | 0.0004498481649 | |
| GO:0046620 | regulation of organ growth | biological process | 0.0004835045618 | |
| GO:0044260 | cellular macromolecule metabolic process | biological process | | 0.0004900966471 |
| GO:0048024 | regulation of mRNA splicing, via spliceosome | biological process | 0.0005357307388 | |
| GO:0002065 | columnar/cuboidal epithelial cell differentiation | biological process | 0.0005464422865 | |
| GO:0007442 | hindgut morphogenesis | biological process | 0.0005464422865 | |
| GO:0035186 | syncytial blastoderm mitotic cell cycle | biological process | 0.0005464422865 | |
| GO:1990709 | presynaptic active zone organization | biological process | 0.0005464422865 | |
| GO:0031290 | retinal ganglion cell axon guidance | biological process | 0.0005465772334 | |
| GO:0048584 | positive regulation of response to stimulus | biological process | 0.0005465772334 | |
| GO:0030516 | regulation of axon extension | biological process | 0.0005494217494 | |
| GO:0055059 | asymmetric neuroblast division | biological process | 0.0005494217494 | |
| GO:0061387 | regulation of extent of cell growth | biological process | 0.0005494217494 | |
| GO:0090090 | negative regulation of canonical Wnt signaling pathway | biological process | 0.0005494217494 | |
| GO:0008595 | anterior/posterior axis specification, embryo | biological process | 0.0006040051636 | |
| GO:0001704 | formation of primary germ layer | biological process | 0.000642279603 | |
| GO:0007438 | oenocyte development | biological process | 0.000642279603 | |
| GO:0007440 | foregut morphogenesis | biological process | 0.000642279603 | |
| GO:0035221 | genital disc pattern formation | biological process | 0.000642279603 | |
| GO:0035224 | genital disc anterior/posterior pattern formation | biological process | 0.000642279603 | |
| GO:0060573 | cell fate specification involved in pattern specification | biological process | 0.000642279603 | |
| GO:0071162 | CMG complex | cellular component | 0.0006654061887 | |
| GO:0035639 | purine ribonucleoside triphosphate binding | molecular function | | 0.0006732344485 |
| GO:0002683 | negative regulation of immune system process | biological process | 0.0006945581059 | |
| GO:0098722 | asymmetric stem cell division | biological process | 0.0007181005894 | |
| GO:0010941 | regulation of cell death | biological process | 0.0007662840611 | |

| | | | | |
|---|---|---|---|---|
| GO:0016310 | phosphorylation | biological process | | 0.0008072839397 |
| GO:0030534 | adult behavior | biological process | 0.0008155995942 | |
| GO:0008354 | germ cell migration | biological process | 0.0008389604499 | |
| GO:0030178 | negative regulation of Wnt signaling pathway | biological process | 0.0008389604499 | |
| GO:0005911 | cell-cell junction | cellular component | 0.0008711933938 | |
| GO:0016348 | imaginal disc-derived leg joint morphogenesis | biological process | 0.0009030097511 | |
| GO:0042387 | plasmatocyte differentiation | biological process | 0.0009030097511 | |
| GO:0042478 | regulation of eye photoreceptor cell development | biological process | 0.0009030097511 | |
| GO:0042689 | regulation of crystal cell differentiation | biological process | 0.0009030097511 | |
| GO:0045613 | regulation of plasmatocyte differentiation | biological process | 0.0009030097511 | |
| GO:0048806 | genitalia development | biological process | 0.0009030097511 | |
| GO:0048665 | neuron fate specification | biological process | 0.0009361093106 | |
| GO:0050808 | synapse organization | biological process | 0.000969624178 | |
| GO:0043902 | positive regulation of multi-organism process | biological process | 0.0009729477613 | |
| GO:0001742 | oenocyte differentiation | biological process | 0.001006730796 | |
| GO:0001752 | compound eye photoreceptor fate commitment | biological process | 0.001006730796 | |
| GO:0007521 | muscle cell fate determination | biological process | 0.001006730796 | |
| GO:0016200 | synaptic target attraction | biological process | 0.001006730796 | |
| GO:0035284 | brain segmentation | biological process | 0.001006730796 | |
| GO:0036058 | filtration diaphragm assembly | biological process | 0.001006730796 | |
| GO:0042686 | regulation of cardioblast cell fate specification | biological process | 0.001006730796 | |
| GO:0042706 | eye photoreceptor cell fate commitment | biological process | 0.001006730796 | |
| GO:2000043 | regulation of cardiac cell fate specification | biological process | 0.001006730796 | |
| GO:0032555 | purine ribonucleotide binding | molecular function | | 0.001022908113 |
| GO:0005730 | nucleolus | cellular component | 0.001090220333 | |
| GO:0032508 | DNA duplex unwinding | biological process | 0.001096707955 | |
| GO:0042386 | hemocyte differentiation | biological process | 0.001096707955 | |
| GO:0042325 | regulation of phosphorylation | biological process | 0.001123437788 | |
| GO:0048863 | stem cell differentiation | biological process | 0.001141923341 | |
| GO:0031261 | DNA replication preinitiation complex | cellular component | 0.001145683225 | |
| GO:0030684 | preribosome | cellular component | 0.001156532413 | |
| GO:0030424 | axon | cellular component | 0.001179953338 | |
| GO:0000904 | cell morphogenesis involved in differentiation | biological process | 0.001214064585 | |

| | | | | |
|---|---|---|---|---|
| GO:0007418 | ventral midline development | biological process | 0.001253804027 | |
| GO:0007523 | larval visceral muscle development | biological process | 0.001253804027 | |
| GO:0046667 | compound eye retinal cell programmed cell death | biological process | 0.001253804027 | |
| GO:0048664 | neuron fate determination | biological process | 0.001253804027 | |
| GO:0061382 | Malpighian tubule tip cell differentiation | biological process | 0.001253804027 | |
| GO:0017076 | purine nucleotide binding | molecular function | | 0.001282386012 |
| GO:0046914 | transition metal ion binding | molecular function | | 0.001282386012 |
| GO:0007613 | memory | biological process | 0.001283427402 | |
| GO:0032268 | regulation of cellular protein metabolic process | biological process | 0.001368185127 | |
| GO:0035206 | regulation of hemocyte proliferation | biological process | 0.001392247863 | |
| GO:0051246 | regulation of protein metabolic process | biological process | 0.001400433858 | |
| GO:0042981 | regulation of apoptotic process | biological process | 0.001400882356 | |
| GO:0070192 | chromosome organization involved in meiotic cell cycle | biological process | 0.001409898101 | |
| GO:0007173 | epidermal growth factor receptor signaling pathway | biological process | 0.001444436731 | |
| GO:0038127 | ERBB signaling pathway | biological process | 0.001444436731 | |
| GO:0035220 | wing disc development | biological process | 0.001523022142 | |
| GO:0007616 | long-term memory | biological process | 0.001555395204 | |
| GO:0035223 | leg disc pattern formation | biological process | 0.001596462511 | |
| GO:0051783 | regulation of nuclear division | biological process | 0.001749784277 | |
| GO:0017116 | single-stranded DNA helicase activity | molecular function | 0.001781340909 | |
| GO:0016319 | mushroom body development | biological process | 0.001794751184 | |
| GO:0060828 | regulation of canonical Wnt signaling pathway | biological process | 0.001869838126 | |
| GO:0032553 | ribonucleotide binding | molecular function | | 0.001892599305 |
| GO:0000578 | embryonic axis specification | biological process | 0.001898073155 | |
| GO:0032392 | DNA geometric change | biological process | 0.001898073155 | |
| GO:0007088 | regulation of mitotic nuclear division | biological process | 0.002051959805 | |
| GO:0030858 | positive regulation of epithelial cell differentiation | biological process | 0.002070990573 | |
| GO:0071897 | DNA biosynthetic process | biological process | 0.002070990573 | |
| GO:0007628 | adult walking behavior | biological process | 0.002117083623 | |
| GO:0022409 | positive regulation of cell-cell adhesion | biological process | 0.002117083623 | |
| GO:0031399 | regulation of protein modification process | biological process | 0.002133649693 | |
| GO:0032101 | regulation of response to external stimulus | biological process | 0.002184243751 | |
| GO:0007431 | salivary gland development | biological process | 0.002189155517 | |

| | | | | |
|---|---|---|---|---|
| GO:0000381 | regulation of alternative mRNA splicing, via spliceosome | biological process | 0.002206515478 | |
| GO:0007096 | regulation of exit from mitosis | biological process | 0.002206515478 | |
| GO:0046847 | filopodium assembly | biological process | 0.002206515478 | |
| GO:0048036 | central complex development | biological process | 0.002206515478 | |
| GO:0061320 | pericardial nephrocyte differentiation | biological process | 0.002206515478 | |
| GO:0000808 | origin recognition complex | cellular component | | 0.002428261182 |
| GO:2000648 | positive regulation of stem cell proliferation | biological process | 0.002438933002 | |
| GO:0000792 | heterochromatin | cellular component | 0.002566351905 | |
| GO:0008052 | sensory organ boundary specification | biological process | 0.002571983621 | |
| GO:0043388 | positive regulation of DNA binding | biological process | 0.002571983621 | |
| GO:0045317 | equator specification | biological process | 0.002571983621 | |
| GO:1900087 | positive regulation of G1/S transition of mitotic cell cycle | biological process | 0.002571983621 | |
| GO:1902808 | positive regulation of cell cycle G1/S phase transition | biological process | 0.002571983621 | |
| GO:0043484 | regulation of RNA splicing | biological process | 0.002632312117 | |
| GO:0045170 | spectrosome | cellular component | 0.002633623405 | |
| GO:0008593 | regulation of Notch signaling pathway | biological process | 0.00267362016 | |
| GO:0003729 | mRNA binding | molecular function | 0.002844781628 | |
| GO:0007480 | imaginal disc-derived leg morphogenesis | biological process | 0.002867649007 | |
| GO:1902806 | regulation of cell cycle G1/S phase transition | biological process | 0.002933530186 | |
| GO:2000045 | regulation of G1/S transition of mitotic cell cycle | biological process | 0.002933530186 | |
| GO:0048754 | branching morphogenesis of an epithelial tube | biological process | 0.002970821394 | |
| GO:1902275 | regulation of chromatin organization | biological process | 0.002970821394 | |
| GO:0007479 | leg disc proximal/distal pattern formation | biological process | 0.003042536642 | |
| GO:0008156 | negative regulation of DNA replication | biological process | 0.003042536642 | |
| GO:0042688 | crystal cell differentiation | biological process | 0.003042536642 | |
| GO:0055001 | muscle cell development | biological process | 0.003042536642 | |
| GO:0007403 | glial cell fate determination | biological process | 0.003087359347 | |
| GO:0007429 | secondary branching, open tracheal system | biological process | 0.003087359347 | |
| GO:0014018 | neuroblast fate specification | biological process | 0.003087359347 | |
| GO:0035385 | Roundabout signaling pathway | biological process | 0.003087359347 | |
| GO:0048866 | stem cell fate specification | biological process | 0.003087359347 | |
| GO:0045786 | negative regulation of cell cycle | biological process | 0.003202532835 | |
| GO:0090068 | positive regulation of cell cycle process | biological process | 0.003329150001 | |

| | | | | |
|---|---|---|---|---|
| GO:0008258 | head involution | biological process | 0.003482044524 | |
| GO:0001763 | morphogenesis of a branching structure | biological process | 0.003508099714 | |
| GO:0030856 | regulation of epithelial cell differentiation | biological process | 0.003508099714 | |
| GO:0061138 | morphogenesis of a branching epithelium | biological process | 0.003508099714 | |
| GO:0006268 | DNA unwinding involved in DNA replication | biological process | 0.003571966652 | |
| GO:0007482 | haltere development | biological process | 0.003571966652 | |
| GO:0045614 | negative regulation of plasmatocyte differentiation | biological process | 0.003571966652 | |
| GO:0046666 | retinal cell programmed cell death | biological process | 0.003571966652 | |
| GO:0048790 | maintenance of presynaptic active zone structure | biological process | 0.003571966652 | |
| GO:0051304 | chromosome separation | biological process | 0.003571966652 | |
| GO:0016773 | phosphotransferase activity, alcohol group as acceptor | molecular function | | 0.003771655533 |
| GO:0010942 | positive regulation of cell death | biological process | 0.00390816122 | |
| GO:0030054 | cell junction | cellular component | 0.004315303232 | |
| GO:0006996 | organelle organization | biological process | 0.004408089672 | |
| GO:0007626 | locomotory behavior | biological process | 0.004408089672 | |
| GO:0019827 | stem cell population maintenance | biological process | 0.004408089672 | |
| GO:0098727 | maintenance of cell number | biological process | 0.004408089672 | |
| GO:0060249 | anatomical structure homeostasis | biological process | 0.004532320564 | |
| GO:0033044 | regulation of chromosome organization | biological process | 0.004798121153 | |
| GO:0007443 | Malpighian tubule morphogenesis | biological process | 0.004816495361 | |
| GO:0035332 | positive regulation of hippo signaling | biological process | 0.004816495361 | |
| GO:0051098 | regulation of binding | biological process | 0.004816495361 | |
| GO:0061333 | renal tubule morphogenesis | biological process | 0.004816495361 | |
| GO:1905881 | positive regulation of oogenesis | biological process | 0.00493582979 | |
| GO:0030308 | negative regulation of cell growth | biological process | 0.004990974738 | |
| GO:1903688 | positive regulation of border follicle cell migration | biological process | 0.005098032446 | |
| GO:0007402 | ganglion mother cell fate determination | biological process | 0.005151046699 | |
| GO:0010032 | meiotic chromosome condensation | biological process | 0.005151046699 | |
| GO:0016204 | determination of muscle attachment site | biological process | 0.005151046699 | |
| GO:0042690 | negative regulation of crystal cell differentiation | biological process | 0.005151046699 | |
| GO:0042694 | muscle cell fate specification | biological process | 0.005151046699 | |
| GO:0045677 | negative regulation of R7 cell differentiation | biological process | 0.005151046699 | |
| GO:0050918 | positive chemotaxis | biological process | 0.005151046699 | |

| | | | | |
|---|---|---|---|---|
| GO:0003697 | single-stranded DNA binding | molecular function | 0.005195392504 | |
| GO:0030111 | regulation of Wnt signaling pathway | biological process | 0.005333783157 | |
| GO:0007464 | R3/R4 cell fate commitment | biological process | 0.005543658215 | |
| GO:0045314 | regulation of compound eye photoreceptor development | biological process | 0.005543658215 | |
| GO:0061327 | anterior Malpighian tubule development | biological process | 0.005543658215 | |
| GO:0030707 | ovarian follicle cell development | biological process | 0.005677474335 | |
| GO:0000079 | regulation of cyclin-dependent protein serine/threonine kinase activity | biological process | 0.00580231283 | |
| GO:1904029 | regulation of cyclin-dependent protein kinase activity | biological process | 0.00580231283 | |
| GO:2000243 | positive regulation of reproductive process | biological process | 0.005921339723 | |
| GO:0007367 | segment polarity determination | biological process | 0.006016884465 | |
| GO:0010634 | positive regulation of epithelial cell migration | biological process | 0.006016884465 | |
| GO:0003727 | single-stranded RNA binding | molecular function | 0.00607142362 | |
| GO:0005886 | plasma membrane | cellular component | 0.006077462692 | |
| GO:0008543 | fibroblast growth factor receptor signaling pathway | biological process | 0.006281253143 | |
| GO:0046621 | negative regulation of organ growth | biological process | 0.006281253143 | |
| GO:0110110 | positive regulation of animal organ morphogenesis | biological process | 0.006281253143 | |
| GO:2000179 | positive regulation of neural precursor cell proliferation | biological process | 0.006281253143 | |
| GO:0002066 | columnar/cuboidal epithelial cell development | biological process | 0.006337886134 | |
| GO:0045666 | positive regulation of neuron differentiation | biological process | 0.006556962541 | |
| GO:0044295 | axonal growth cone | cellular component | 0.006638291024 | |
| GO:0050773 | regulation of dendrite development | biological process | 0.006657289381 | |
| GO:0043065 | positive regulation of apoptotic process | biological process | 0.006689779942 | |
| GO:0090575 | RNA polymerase II transcription regulator complex | cellular component | 0.00689638962 | |
| GO:1903684 | regulation of border follicle cell migration | biological process | 0.007158415353 | |
| GO:0006310 | DNA recombination | biological process | 0.007321435466 | |
| GO:0004386 | helicase activity | molecular function | | 0.007492973556 |
| GO:0043138 | 3'-5' DNA helicase activity | molecular function | 0.007612351397 | |
| GO:0007162 | negative regulation of cell adhesion | biological process | 0.007809118689 | |
| GO:0007449 | proximal/distal pattern formation, imaginal disc | biological process | 0.007809118689 | |
| GO:0000166 | nucleotide binding | molecular function | | 0.007827630665 |
| GO:1901265 | nucleoside phosphate binding | molecular function | | 0.007827630665 |
| GO:0090329 | regulation of DNA-dependent DNA replication | biological process | 0.007992994214 | |
| GO:0006302 | double-strand break repair | biological process | 0.008161462205 | |

| | | | | |
|---|---|---|---|---|
| GO:0048737 | imaginal disc-derived appendage development | biological process | 0.008161462205 | |
| GO:0099558 | maintenance of synapse structure | biological process | 0.008161462205 | |
| GO:0016301 | kinase activity | molecular function | | 0.008225649685 |
| GO:0005856 | cytoskeleton | cellular component | 0.008382300819 | |
| GO:0031056 | regulation of histone modification | biological process | 0.00847823745 | |
| GO:0045570 | regulation of imaginal disc growth | biological process | 0.00847823745 | |
| GO:0060446 | branching involved in open tracheal system development | biological process | 0.00847823745 | |
| GO:0031497 | chromatin assembly | biological process | 0.008613323406 | |
| GO:0016203 | muscle attachment | biological process | 0.008667537074 | |
| GO:0030855 | epithelial cell differentiation | biological process | 0.008667537074 | |
| GO:0000724 | double-strand break repair via homologous recombination | biological process | 0.008979020151 | |
| GO:0000725 | recombinational repair | biological process | 0.008979020151 | |
| GO:0008584 | male gonad development | biological process | 0.009111596668 | |
| GO:0010002 | cardioblast differentiation | biological process | 0.009111596668 | |
| GO:0051383 | kinetochore organization | biological process | 0.009111596668 | |
| GO:0006464 | cellular protein modification process | biological process | | 0.009560459623 |
| GO:0036211 | protein modification process | biological process | | 0.009560459623 |
| GO:0001932 | regulation of protein phosphorylation | biological process | 0.00994803946 | |
| GO:0001706 | endoderm formation | biological process | 0.009971542696 | |
| GO:0002064 | epithelial cell development | biological process | 0.009971542696 | |
| GO:0003151 | outflow tract morphogenesis | biological process | 0.009971542696 | |
| GO:0009997 | negative regulation of cardioblast cell fate specification | biological process | 0.009971542696 | |
| GO:0010092 | specification of animal organ identity | biological process | 0.009971542696 | |
| GO:0035154 | terminal cell fate specification, open tracheal system | biological process | 0.009971542696 | |
| GO:0035561 | regulation of chromatin binding | biological process | 0.009971542696 | |
| GO:0036059 | nephrocyte diaphragm assembly | biological process | 0.009971542696 | |
| GO:0042063 | gliogenesis | biological process | 0.009971542696 | |
| GO:0044728 | DNA methylation or demethylation | biological process | 0.009971542696 | |
| GO:0051382 | kinetochore assembly | biological process | 0.009971542696 | |
| GO:0051892 | negative regulation of cardioblast differentiation | biological process | 0.009971542696 | |
| GO:0061321 | garland nephrocyte differentiation | biological process | 0.009971542696 | |
| GO:1905208 | negative regulation of cardiocyte differentiation | biological process | 0.009971542696 | |
| GO:2000044 | negative regulation of cardiac cell fate specification | biological process | 0.009971542696 | |

| | | | | |
|---|---|---|---|---|
| GO:0051303 | establishment of chromosome localization | biological process | 0.01003184937 | |
| GO:0030261 | chromosome condensation | biological process | 0.01020438457 | |
| GO:0007427 | epithelial cell migration, open tracheal system | biological process | 0.01024540071 | |
| GO:0000775 | chromosome, centromeric region | cellular component | 0.01073855285 | |
| GO:0043412 | macromolecule modification | biological process | | 0.01081352252 |
| GO:0022408 | negative regulation of cell-cell adhesion | biological process | 0.0113442078 | |
| GO:0035065 | regulation of histone acetylation | biological process | 0.0113442078 | |
| GO:0048634 | regulation of muscle organ development | biological process | 0.0113442078 | |
| GO:0048800 | antennal morphogenesis | biological process | 0.0113442078 | |
| GO:1901983 | regulation of protein acetylation | biological process | 0.0113442078 | |
| GO:2000756 | regulation of peptidyl-lysine acetylation | biological process | 0.0113442078 | |
| GO:0001894 | tissue homeostasis | biological process | 0.01136968005 | |
| GO:0040014 | regulation of multicellular organism growth | biological process | 0.01136968005 | |
| GO:0042058 | regulation of epidermal growth factor receptor signaling pathway | biological process | 0.01136968005 | |
| GO:1901184 | regulation of ERBB signaling pathway | biological process | 0.01136968005 | |
| GO:0009967 | positive regulation of signal transduction | biological process | 0.0114537456 | |
| GO:0048495 | Roundabout binding | molecular function | 0.01162553656 | |
| GO:0005701 | polytene chromosome chromocenter | cellular component | 0.01166310223 | |
| GO:0032269 | negative regulation of cellular protein metabolic process | biological process | 0.01174828323 | |
| GO:0032200 | telomere organization | biological process | 0.0121830199 | |
| GO:0051248 | negative regulation of protein metabolic process | biological process | 0.0121830199 | |
| GO:0010647 | positive regulation of cell communication | biological process | 0.0125359341 | |
| GO:0023056 | positive regulation of signaling | biological process | 0.0125359341 | |
| GO:0031507 | heterochromatin assembly | biological process | 0.01280052257 | |
| GO:0034401 | chromatin organization involved in regulation of transcription | biological process | 0.01280052257 | |
| GO:0097549 | chromatin organization involved in negative regulation of transcription | biological process | 0.01280052257 | |
| GO:0007413 | axonal fasciculation | biological process | 0.01284597431 | |
| GO:0010369 | chromocenter | cellular component | 0.01322280993 | |
| GO:0008586 | imaginal disc-derived wing vein morphogenesis | biological process | 0.0132467487 | |
| GO:0031401 | positive regulation of protein modification process | biological process | 0.01329314102 | |
| GO:0031175 | neuron projection development | biological process | 0.01330839446 | |
| GO:0006352 | DNA-templated transcription, initiation | biological process | | 0.01437415859 |
| GO:0001746 | Bolwig's organ morphogenesis | biological process | 0.01444734301 | |

| | | | | |
|---|---|---|---|---|
| GO:0048568 | embryonic organ development | biological process | 0.01444734301 | |
| GO:0051099 | positive regulation of binding | biological process | 0.01444734301 | |
| GO:0060571 | morphogenesis of an epithelial fold | biological process | 0.01444734301 | |
| GO:0034332 | adherens junction organization | biological process | 0.01463046171 | |
| GO:0016278 | lysine N-methyltransferase activity | molecular function | | 0.01465036707 |
| GO:0016279 | protein-lysine N-methyltransferase activity | molecular function | | 0.01465036707 |
| GO:0018024 | histone-lysine N-methyltransferase activity | molecular function | | 0.01465036707 |
| GO:0042054 | histone methyltransferase activity | molecular function | | 0.01465036707 |
| GO:0030307 | positive regulation of cell growth | biological process | 0.01535544212 | |
| GO:0016201 | synaptic target inhibition | biological process | 0.01547366251 | |
| GO:0035168 | larval lymph gland hemocyte differentiation | biological process | 0.01547366251 | |
| GO:0045478 | fusome organization | biological process | 0.01547366251 | |
| GO:0010171 | body morphogenesis | biological process | 0.01632798533 | |
| GO:0034331 | cell junction maintenance | biological process | 0.01632798533 | |
| GO:0106030 | neuron projection fasciculation | biological process | 0.01632798533 | |
| GO:1905879 | regulation of oogenesis | biological process | 0.01656522912 | |
| GO:0005704 | polytene chromosome band | cellular component | 0.01672933636 | |
| GO:0040029 | regulation of gene expression, epigenetic | biological process | 0.01700412717 | |
| GO:0071900 | regulation of protein serine/threonine kinase activity | biological process | 0.01700412717 | |
| GO:0042327 | positive regulation of phosphorylation | biological process | 0.01756706075 | |
| GO:0019220 | regulation of phosphate metabolic process | biological process | 0.01808997297 | |
| GO:0051174 | regulation of phosphorus metabolic process | biological process | 0.01808997297 | |
| GO:0010948 | negative regulation of cell cycle process | biological process | 0.01822950411 | |
| GO:0045930 | negative regulation of mitotic cell cycle | biological process | 0.01822950411 | |
| GO:0045216 | cell-cell junction organization | biological process | 0.0184122945 | |
| GO:0065009 | regulation of molecular function | biological process | 0.01882330958 | |
| GO:0097367 | carbohydrate derivative binding | molecular function | | 0.01901510514 |
| GO:0005604 | basement membrane | cellular component | 0.01965410402 | |
| GO:0005917 | nephrocyte diaphragm | cellular component | 0.01965410402 | |
| GO:0031213 | RSF complex | cellular component | 0.01965410402 | |
| GO:0036056 | filtration diaphragm | cellular component | 0.01965410402 | |
| GO:0051302 | regulation of cell division | biological process | 0.02049351603 | |
| GO:0050000 | chromosome localization | biological process | 0.02062147588 | |

| | | | |
|---|---|---|---|
| GO:0003013 | circulatory system process | biological process | 0.02087679759 |
| GO:0003015 | heart process | biological process | 0.02087679759 |
| GO:0006304 | DNA modification | biological process | 0.02135353956 |
| GO:0007501 | mesodermal cell fate specification | biological process | 0.02135353956 |
| GO:0007502 | digestive tract mesoderm development | biological process | 0.02135353956 |
| GO:0007540 | sex determination, establishment of X:A ratio | biological process | 0.02135353956 |
| GO:0014017 | neuroblast fate commitment | biological process | 0.02135353956 |
| GO:0031509 | subtelomeric heterochromatin assembly | biological process | 0.02135353956 |
| GO:0032202 | telomere assembly | biological process | 0.02135353956 |
| GO:0035171 | lamellocyte differentiation | biological process | 0.02135353956 |
| GO:0035310 | notum cell fate specification | biological process | 0.02135353956 |
| GO:0042682 | regulation of compound eye cone cell fate specification | biological process | 0.02135353956 |
| GO:0060323 | head morphogenesis | biological process | 0.02135353956 |
| GO:0060911 | cardiac cell fate commitment | biological process | 0.02135353956 |
| GO:0071168 | protein localization to chromatin | biological process | 0.02135353956 |
| GO:0140461 | subtelomeric heterochromatin organization | biological process | 0.02135353956 |
| GO:0035202 | tracheal pit formation in open tracheal system | biological process | 0.02158497593 |
| GO:1990511 | piRNA biosynthetic process | biological process | 0.02158497593 |
| GO:2000274 | regulation of epithelial cell migration, open tracheal system | biological process | 0.02158497593 |
| GO:0060560 | developmental growth involved in morphogenesis | biological process | 0.02229988276 |
| GO:0004714 | transmembrane receptor protein tyrosine kinase activity | molecular function | 0.02243102676 |
| GO:0042995 | cell projection | cellular component | 0.02276848392 |
| GO:0050790 | regulation of catalytic activity | biological process | 0.02278169689 |
| GO:0022416 | chaeta development | biological process | 0.02295202045 |
| GO:0030426 | growth cone | cellular component | 0.02323683814 |
| GO:0000075 | cell cycle checkpoint | biological process | 0.02471928595 |
| GO:0009950 | dorsal/ventral axis specification | biological process | 0.02471928595 |
| GO:0030863 | cortical cytoskeleton | cellular component | 0.02472628717 |
| GO:0002052 | positive regulation of neuroblast proliferation | biological process | 0.02499602426 |
| GO:0042059 | negative regulation of epidermal growth factor receptor signaling pathway | biological process | 0.02499602426 |
| GO:0043954 | cellular component maintenance | biological process | 0.02499602426 |
| GO:0045860 | positive regulation of protein kinase activity | biological process | 0.02499602426 |
| GO:0070828 | heterochromatin organization | biological process | 0.02499602426 |

| | | | | |
|---|---|---|---|---|
| GO:1901185 | negative regulation of ERBB signaling pathway | biological process | 0.02499602426 | |
| GO:0019207 | kinase regulator activity | molecular function | 0.02533036803 | |
| GO:0000902 | cell morphogenesis | biological process | 0.02633203508 | |
| GO:0000076 | DNA replication checkpoint | biological process | 0.02656247498 | |
| GO:0007362 | terminal region determination | biological process | 0.02656247498 | |
| GO:0031570 | DNA integrity checkpoint | biological process | 0.02656247498 | |
| GO:0045167 | asymmetric protein localization involved in cell fate determination | biological process | 0.02656247498 | |
| GO:0046426 | negative regulation of receptor signaling pathway via JAK-STAT | biological process | 0.02656247498 | |
| GO:0048639 | positive regulation of developmental growth | biological process | 0.02656247498 | |
| GO:0051310 | metaphase plate congression | biological process | 0.02656247498 | |
| GO:1904893 | negative regulation of receptor signaling pathway via STAT | biological process | 0.02656247498 | |
| GO:0061982 | meiosis I cell cycle process | biological process | 0.02695024767 | |
| GO:0035102 | PRC1 complex | cellular component | 0.02765507168 | |
| GO:0120025 | plasma membrane bounded cell projection | cellular component | 0.02784817406 | |
| GO:0000307 | cyclin-dependent protein kinase holoenzyme complex | cellular component | 0.02890865189 | |
| GO:0016772 | transferase activity, transferring phosphorus-containing groups | molecular function | | 0.02916476497 |
| GO:0019199 | transmembrane receptor protein kinase activity | molecular function | | 0.02916476497 |
| GO:0036094 | small molecule binding | molecular function | | 0.02916476497 |
| GO:0016324 | apical plasma membrane | cellular component | 0.02971493677 | |
| GO:0016327 | apicolateral plasma membrane | cellular component | 0.02971493677 | |
| GO:1901991 | negative regulation of mitotic cell cycle phase transition | biological process | 0.02972577126 | |
| GO:0030718 | germ-line stem cell population maintenance | biological process | 0.02989360888 | |
| GO:0016055 | Wnt signaling pathway | biological process | 0.03050250343 | |
| GO:0045785 | positive regulation of cell adhesion | biological process | 0.03050250343 | |
| GO:1905114 | cell surface receptor signaling pathway involved in cell-cell signaling | biological process | 0.03050250343 | |
| GO:0019887 | protein kinase regulator activity | molecular function | 0.03053294424 | |
| GO:0030723 | ovarian fusome organization | biological process | 0.0305603422 | |
| GO:0035203 | regulation of lamellocyte differentiation | biological process | 0.0305603422 | |
| GO:0035215 | genital disc development | biological process | 0.0305603422 | |
| GO:0035330 | regulation of hippo signaling | biological process | 0.0305603422 | |
| GO:0042766 | nucleosome mobilization | biological process | 0.0305603422 | |
| GO:0044772 | mitotic cell cycle phase transition | biological process | 0.0305603422 | |
| GO:0046427 | positive regulation of receptor signaling pathway via JAK-STAT | biological process | 0.0305603422 | |

| | | | | |
|---|---|---|---|---|
| GO:0048841 | regulation of axon extension involved in axon guidance | biological process | 0.0305603422 | |
| GO:0060232 | delamination | biological process | 0.0305603422 | |
| GO:1904894 | positive regulation of receptor signaling pathway via STAT | biological process | 0.0305603422 | |
| GO:2000737 | negative regulation of stem cell differentiation | biological process | 0.0305603422 | |
| GO:0001221 | transcription cofactor binding | molecular function | 0.03057745278 | |
| GO:0030427 | site of polarized growth | cellular component | 0.03189715307 | |
| GO:0045927 | positive regulation of growth | biological process | 0.0331102548 | |
| GO:1901988 | negative regulation of cell cycle phase transition | biological process | 0.03326929706 | |
| GO:0004674 | protein serine/threonine kinase activity | molecular function | | 0.03341812203 |
| GO:0034333 | adherens junction assembly | biological process | 0.03386355407 | |
| GO:0048100 | wing disc anterior/posterior pattern formation | biological process | 0.03386355407 | |
| GO:0070593 | dendrite self-avoidance | biological process | 0.03386355407 | |
| GO:0007043 | cell-cell junction assembly | biological process | 0.03405205062 | |
| GO:0031208 | POZ domain binding | molecular function | 0.03418961364 | |
| GO:0006305 | DNA alkylation | biological process | 0.03538015606 | |
| GO:0006306 | DNA methylation | biological process | 0.03538015606 | |
| GO:0007157 | heterophilic cell-cell adhesion via plasma membrane cell adhesion molecul | biological process | 0.03538015606 | |
| GO:0007355 | anterior region determination | biological process | 0.03538015606 | |
| GO:0007376 | cephalic furrow formation | biological process | 0.03538015606 | |
| GO:0007381 | specification of segmental identity, labial segment | biological process | 0.03538015606 | |
| GO:0007382 | specification of segmental identity, maxillary segment | biological process | 0.03538015606 | |
| GO:0007383 | specification of segmental identity, antennal segment | biological process | 0.03538015606 | |
| GO:0007421 | stomatogastric nervous system development | biological process | 0.03538015606 | |
| GO:0031099 | regeneration | biological process | 0.03538015606 | |
| GO:0031345 | negative regulation of cell projection organization | biological process | 0.03538015606 | |
| GO:0034114 | regulation of heterotypic cell-cell adhesion | biological process | 0.03538015606 | |
| GO:0034116 | positive regulation of heterotypic cell-cell adhesion | biological process | 0.03538015606 | |
| GO:0035038 | female pronucleus assembly | biological process | 0.03538015606 | |
| GO:0035155 | negative regulation of terminal cell fate specification, open tracheal system | biological process | 0.03538015606 | |
| GO:0035157 | negative regulation of fusion cell fate specification | biological process | 0.03538015606 | |
| GO:0035260 | internal genitalia morphogenesis | biological process | 0.03538015606 | |
| GO:0038007 | netrin-activated signaling pathway | biological process | 0.03538015606 | |
| GO:0042062 | long-term strengthening of neuromuscular junction | biological process | 0.03538015606 | |

| | | | |
|---|---|---|---|
| GO:0042693 | muscle cell fate commitment | biological process | 0.03538015606 |
| GO:0045685 | regulation of glial cell differentiation | biological process | 0.03538015606 |
| GO:0045687 | positive regulation of glial cell differentiation | biological process | 0.03538015606 |
| GO:0048511 | rhythmic process | biological process | 0.03538015606 |
| GO:0048936 | peripheral nervous system neuron axonogenesis | biological process | 0.03538015606 |
| GO:0060233 | oenocyte delamination | biological process | 0.03538015606 |
| GO:0071921 | cohesin loading | biological process | 0.03538015606 |
| GO:1901989 | positive regulation of cell cycle phase transition | biological process | 0.03538015606 |
| GO:1901992 | positive regulation of mitotic cell cycle phase transition | biological process | 0.03538015606 |
| GO:1902337 | regulation of apoptotic process involved in morphogenesis | biological process | 0.03538015606 |
| GO:1902339 | positive regulation of apoptotic process involved in morphogenesis | biological process | 0.03538015606 |
| GO:0007313 | maternal specification of dorsal/ventral axis, oocyte, soma encoded | biological process | 0.03584587043 |
| GO:0016202 | regulation of striated muscle tissue development | biological process | 0.03584587043 |
| GO:0038004 | epidermal growth factor receptor ligand maturation | biological process | 0.03584587043 |
| GO:0043703 | photoreceptor cell fate determination | biological process | 0.03584587043 |
| GO:0046665 | amnioserosa maintenance | biological process | 0.03584587043 |
| GO:0048645 | animal organ formation | biological process | 0.03584587043 |
| GO:0048854 | brain morphogenesis | biological process | 0.03584587043 |
| GO:0060914 | heart formation | biological process | 0.03584587043 |
| GO:1901861 | regulation of muscle tissue development | biological process | 0.03584587043 |
| GO:1902292 | cell cycle DNA replication initiation | biological process | 0.03584587043 |
| GO:1902315 | nuclear cell cycle DNA replication initiation | biological process | 0.03584587043 |
| GO:1902975 | mitotic DNA replication initiation | biological process | 0.03584587043 |
| GO:1904666 | regulation of ubiquitin protein ligase activity | biological process | 0.03584587043 |
| GO:0006323 | DNA packaging | biological process | 0.03612055155 |
| GO:0090287 | regulation of cellular response to growth factor stimulus | biological process | 0.03612055155 |
| GO:0043900 | regulation of multi-organism process | biological process | 0.03612604315 |
| GO:0008344 | adult locomotory behavior | biological process | 0.03695366106 |
| GO:0016569 | covalent chromatin modification | biological process | 0.03722897229 |
| GO:0016570 | histone modification | biological process | 0.03722897229 |
| GO:0031010 | ISWI-type complex | cellular component | 0.03743806775 |
| GO:0006364 | rRNA processing | biological process | 0.03831645941 |
| GO:0044087 | regulation of cellular component biogenesis | biological process | 0.03888470424 |

| | | | |
|---|---|---|---|
| GO:0008285 | negative regulation of cell population proliferation | biological process | 0.03888691072 |
| GO:0030686 | 90S preribosome | cellular component | 0.03906162009 |
| GO:0003007 | heart morphogenesis | biological process | 0.03997015597 |
| GO:0008594 | photoreceptor cell morphogenesis | biological process | 0.03997015597 |
| GO:0016572 | histone phosphorylation | biological process | 0.03997015597 |
| GO:0035285 | appendage segmentation | biological process | 0.03997015597 |
| GO:0036011 | imaginal disc-derived leg segmentation | biological process | 0.03997015597 |
| GO:0060438 | trachea development | biological process | 0.03997015597 |
| GO:0007390 | germ-band shortening | biological process | 0.0401871888 |
| GO:0035212 | cell competition in a multicellular organism | biological process | 0.0401871888 |
| GO:0051101 | regulation of DNA binding | biological process | 0.0401871888 |
| GO:0007472 | wing disc morphogenesis | biological process | 0.04119322802 |
| GO:0003008 | system process | biological process | 0.04150098814 |
| GO:0001736 | establishment of planar polarity | biological process | 0.04247629732 |
| GO:0007164 | establishment of tissue polarity | biological process | 0.04247629732 |
| GO:0030097 | hemopoiesis | biological process | 0.04247629732 |
| GO:0044770 | cell cycle phase transition | biological process | 0.04247629732 |
| GO:0010562 | positive regulation of phosphorus metabolic process | biological process | 0.0433255395 |
| GO:0045937 | positive regulation of phosphate metabolic process | biological process | 0.0433255395 |
| GO:0045132 | meiotic chromosome segregation | biological process | 0.04342460415 |
| GO:0051124 | synaptic growth at neuromuscular junction | biological process | 0.04342460415 |
| GO:0090288 | negative regulation of cellular response to growth factor stimulus | biological process | 0.04342460415 |
| GO:0098631 | cell adhesion mediator activity | molecular function | 0.04375386551 |
| GO:0000118 | histone deacetylase complex | cellular component | 0.0453417026 |
| GO:0045995 | regulation of embryonic development | biological process | 0.0474906856 |
| GO:0035097 | histone methyltransferase complex | cellular component | 0.04778683196 |
| GO:0016538 | cyclin-dependent protein serine/threonine kinase regulator activity | molecular function | 0.04902529885 |
| GO:0009986 | cell surface | cellular component | 0.04908171234 |
| GO:0007369 | gastrulation | biological process | 0.04982781844 |
| GO:0042067 | establishment of ommatidial planar polarity | biological process | 0.04982781844 |

| GO | name | GO type | p-value FDR *D. melanogaster* | p-value FDR *D. virilis* |
|---|---|---|---|---|
| GO:0003735 | structural constituent of ribosome | molecular function | 4.71E-73 | 3.12E-07 |
| GO:0043043 | peptide biosynthetic process | biological process | 6.21E-71 | 1.12E-06 |
| GO:0006518 | peptide metabolic process | biological process | 1.28E-68 | 1.12E-06 |
| GO:0006412 | translation | biological process | 1.28E-68 | 1.96E-06 |
| GO:0043604 | amide biosynthetic process | biological process | 1.62E-67 | 5.00E-06 |
| GO:0043603 | cellular amide metabolic process | biological process | 1.59E-61 | 9.12E-06 |
| GO:0005198 | structural molecule activity | molecular function | 9.43E-53 | 8.84E-09 |
| GO:1901566 | organonitrogen compound biosynthetic process | biological process | 2.70E-53 | 1.05E-06 |
| GO:0009058 | biosynthetic process | biological process | 9.01E-39 | 1.88E-05 |
| GO:1901576 | organic substance biosynthetic process | biological process | 1.98E-36 | 4.92E-05 |
| GO:0044249 | cellular biosynthetic process | biological process | 1.92E-36 | 5.11E-05 |
| GO:0044271 | cellular nitrogen compound biosynthetic process | biological process | 1.94E-37 | 5.62E-04 |
| GO:0005739 | mitochondrion | cellular component | 1.88E-38 | 4.88E-02 |
| GO:0034645 | cellular macromolecule biosynthetic process | biological process | 1.73E-35 | 2.25E-04 |
| GO:0005840 | ribosome | cellular component | 3.15E-32 | 8.09E-06 |
| GO:0008152 | metabolic process | biological process | 1.59E-20 | 1.25E-09 |
| GO:0009059 | macromolecule biosynthetic process | biological process | 8.22E-27 | 2.99E-02 |
| GO:0044281 | small molecule metabolic process | biological process | 6.89E-21 | 1.12E-06 |
| GO:0019752 | carboxylic acid metabolic process | biological process | 4.30E-22 | 5.58E-05 |
| GO:0043436 | oxoacid metabolic process | biological process | 3.46E-21 | 2.51E-05 |
| GO:0006082 | organic acid metabolic process | biological process | 8.19E-21 | 2.51E-05 |
| GO:1901564 | organonitrogen compound metabolic process | biological process | 8.19E-21 | 1.29E-04 |
| GO:0003824 | catalytic activity | molecular function | 5.60E-17 | 3.12E-07 |
| GO:0034641 | cellular nitrogen compound metabolic process | biological process | 1.52E-19 | 3.59E-02 |
| GO:0016491 | oxidoreductase activity | molecular function | 3.46E-14 | 2.81E-07 |
| GO:0071704 | organic substance metabolic process | biological process | 8.47E-17 | 2.52E-04 |
| GO:0006520 | cellular amino acid metabolic process | biological process | 9.84E-13 | 1.61E-03 |
| GO:0015078 | proton transmembrane transporter activity | molecular function | 8.74E-09 | 9.39E-07 |
| GO:0006807 | nitrogen compound metabolic process | biological process | 3.04E-12 | 0.003180212508 |
| GO:1902600 | proton transmembrane transport | biological process | 5.43E-09 | 4.55E-06 |
| GO:0008061 | chitin binding | molecular function | 9.52E-11 | 0.001780169847 |
| GO:0044238 | primary metabolic process | biological process | 2.44E-11 | 4.59E-02 |

| | | | | |
|---|---|---|---|---|
| GO:0044237 | cellular metabolic process | biological process | 7.20E-10 | 0.01518481445 |
| GO:0033178 | proton-transporting two-sector ATPase complex, catalytic domain | cellular component | 6.48E-06 | 8.18E-06 |
| GO:0055114 | oxidation-reduction process | biological process | 2.42E-05 | 1.90E-05 |
| GO:0055085 | transmembrane transport | biological process | 1.24E-04 | 1.90E-05 |
| GO:0015077 | monovalent inorganic cation transmembrane transporter activity | molecular function | 2.64E-04 | 5.94E-05 |
| GO:0098655 | cation transmembrane transport | biological process | 0.003685908629 | 2.22E-05 |
| GO:0098660 | inorganic ion transmembrane transport | biological process | 0.003801638484 | 2.51E-05 |
| GO:0098662 | inorganic cation transmembrane transport | biological process | 0.004768138126 | 2.22E-05 |
| GO:0009678 | pyrophosphate hydrolysis-driven proton transmembrane transporter activity | molecular function | 9.62E-06 | 0.01119516344 |
| GO:0044769 | ATPase activity, coupled to transmembrane movement of ions, rotational m | molecular function | 9.62E-06 | 1.12E-02 |
| GO:0046961 | proton-transporting ATPase activity, rotational mechanism | molecular function | 9.62E-06 | 1.12E-02 |
| GO:0009145 | purine nucleoside triphosphate biosynthetic process | biological process | 1.82E-04 | 7.22E-04 |
| GO:0009205 | purine ribonucleoside triphosphate metabolic process | biological process | 1.82E-04 | 7.22E-04 |
| GO:0009206 | purine ribonucleoside triphosphate biosynthetic process | biological process | 1.82E-04 | 7.22E-04 |
| GO:0015672 | monovalent inorganic cation transport | biological process | 5.54E-05 | 2.64E-03 |
| GO:0009144 | purine nucleoside triphosphate metabolic process | biological process | 2.77E-04 | 7.22E-04 |
| GO:0098796 | membrane protein complex | cellular component | 1.83E-03 | 1.50E-04 |
| GO:0022804 | active transmembrane transporter activity | molecular function | 1.85E-03 | 2.88E-04 |
| GO:0009199 | ribonucleoside triphosphate metabolic process | biological process | 6.06E-04 | 1.06E-03 |
| GO:0009201 | ribonucleoside triphosphate biosynthetic process | biological process | 6.06E-04 | 1.06E-03 |
| GO:0015985 | energy coupled proton transport, down electrochemical gradient | biological process | 1.05E-04 | 8.68E-03 |
| GO:0015986 | ATP synthesis coupled proton transport | biological process | 1.05E-04 | 8.68E-03 |
| GO:0046933 | proton-transporting ATP synthase activity, rotational mechanism | molecular function | 1.23E-04 | 7.40E-03 |
| GO:0006754 | ATP biosynthetic process | biological process | 1.71E-04 | 8.68E-03 |
| GO:1901135 | carbohydrate derivative metabolic process | biological process | 6.97E-04 | 2.41E-03 |
| GO:0009142 | nucleoside triphosphate biosynthetic process | biological process | 1.74E-03 | 1.06E-03 |
| GO:0009141 | nucleoside triphosphate metabolic process | biological process | 1.04E-03 | 2.13E-03 |
| GO:0009260 | ribonucleotide biosynthetic process | biological process | 1.15E-04 | 2.86E-02 |
| GO:0005215 | transporter activity | molecular function | 2.25E-03 | 1.78E-03 |
| GO:0015252 | proton channel activity | molecular function | 5.74E-04 | 7.40E-03 |
| GO:0046390 | ribose phosphate biosynthetic process | biological process | 1.60E-04 | 2.86E-02 |
| GO:0034220 | ion transmembrane transport | biological process | 3.83E-02 | 1.40E-04 |
| GO:0046034 | ATP metabolic process | biological process | 1.57E-04 | 3.75E-02 |

| | | | | |
|---|---|---|---|---|
| GO:0042626 | ATPase-coupled transmembrane transporter activity | molecular function | 8.14E-04 | 0.007403206122 |
| GO:0015399 | primary active transmembrane transporter activity | molecular function | 0.001998922142 | 0.003343515607 |
| GO:0042302 | structural constituent of cuticle | molecular function | 0.0001661145402 | 4.25E-02 |
| GO:0022857 | transmembrane transporter activity | molecular function | 0.01433850954 | 0.001218864239 |
| GO:0033180 | proton-transporting V-type ATPase, V1 domain | cellular component | 0.0019177334 | 0.02227722038 |
| GO:0006811 | ion transport | biological process | 0.005904991124 | 0.01019448124 |
| GO:0022853 | active ion transmembrane transporter activity | molecular function | 0.006306461199 | 0.01464037518 |
| GO:0045261 | proton-transporting ATP synthase complex, catalytic core F(1) | cellular component | 0.01057661459 | 0.01502690943 |
| GO:0005576 | extracellular region | cellular component | 0.005074127193 | 0.04878956165 |
| GO:0090407 | organophosphate biosynthetic process | biological process | 0.02386841205 | 0.01080294792 |
| GO:0006812 | cation transport | biological process | 0.04087218599 | 0.00904012799 |
| GO:0016769 | transferase activity, transferring nitrogenous groups | molecular function | 0.0496509783 | 0.008346280963 |
| GO:0044391 | ribosomal subunit | cellular component | 6.22E-73 | |
| GO:0098798 | mitochondrial protein complex | cellular component | 3.40E-49 | |
| GO:0015934 | large ribosomal subunit | cellular component | 9.22E-49 | |
| GO:0032543 | mitochondrial translation | biological process | 2.10E-42 | |
| GO:0000315 | organellar large ribosomal subunit | cellular component | 2.43E-32 | |
| GO:0005762 | mitochondrial large ribosomal subunit | cellular component | 2.43E-32 | |
| GO:0022626 | cytosolic ribosome | cellular component | 4.57E-32 | |
| GO:0002181 | cytoplasmic translation | biological process | 3.50E-30 | |
| GO:0015935 | small ribosomal subunit | cellular component | 1.01E-23 | |
| GO:1990904 | ribonucleoprotein complex | cellular component | 2.31E-23 | |
| GO:0022625 | cytosolic large ribosomal subunit | cellular component | 2.38E-18 | |
| GO:0000314 | organellar small ribosomal subunit | cellular component | 1.43E-13 | |
| GO:0005763 | mitochondrial small ribosomal subunit | cellular component | 1.43E-13 | |
| GO:0044282 | small molecule catabolic process | biological process | 2.19E-12 | |
| GO:0016054 | organic acid catabolic process | biological process | 3.78E-12 | |
| GO:0046395 | carboxylic acid catabolic process | biological process | 3.78E-12 | |
| GO:0022627 | cytosolic small ribosomal subunit | cellular component | 6.40E-11 | |
| GO:0098800 | inner mitochondrial membrane protein complex | cellular component | 1.52E-10 | |
| GO:0019843 | rRNA binding | molecular function | 4.29E-10 | |
| GO:0009063 | cellular amino acid catabolic process | biological process | 6.17E-10 | |
| GO:0016469 | proton-transporting two-sector ATPase complex | cellular component | 7.70E-10 | |

| | | | |
|---|---|---|---|
| GO:0032787 | monocarboxylic acid metabolic process | biological process | 1.81E-09 |
| GO:1901606 | alpha-amino acid catabolic process | biological process | 1.55E-08 |
| GO:1901605 | alpha-amino acid metabolic process | biological process | 2.13E-08 |
| GO:0034470 | ncRNA processing | biological process | 3.11E-08 |
| GO:0044283 | small molecule biosynthetic process | biological process | 3.54E-08 |
| GO:0034660 | ncRNA metabolic process | biological process | 7.28E-08 |
| GO:0033176 | proton-transporting V-type ATPase complex | cellular component | 8.14E-08 |
| GO:0006749 | glutathione metabolic process | biological process | 9.02E-08 |
| GO:1990204 | oxidoreductase complex | cellular component | 1.13E-07 |
| GO:0098803 | respiratory chain complex | cellular component | 1.23E-07 |
| GO:0009081 | branched-chain amino acid metabolic process | biological process | 1.30E-07 |
| GO:0033181 | plasma membrane proton-transporting V-type ATPase complex | cellular component | 1.90E-07 |
| GO:0006631 | fatty acid metabolic process | biological process | 2.25E-07 |
| GO:0016072 | rRNA metabolic process | biological process | 2.58E-07 |
| GO:0006364 | rRNA processing | biological process | 4.77E-07 |
| GO:0006575 | cellular modified amino acid metabolic process | biological process | 5.30E-07 |
| GO:0005747 | mitochondrial respiratory chain complex I | cellular component | 5.44E-07 |
| GO:0030964 | NADH dehydrogenase complex | cellular component | 5.44E-07 |
| GO:0045271 | respiratory chain complex I | cellular component | 5.44E-07 |
| GO:0006790 | sulfur compound metabolic process | biological process | 8.60E-07 |
| GO:0009083 | branched-chain amino acid catabolic process | biological process | 9.35E-07 |
| GO:0006399 | tRNA metabolic process | biological process | 2.13E-06 |
| GO:0000027 | ribosomal large subunit assembly | biological process | 2.94E-06 |
| GO:0019538 | protein metabolic process | biological process | 8.68E-06 |
| GO:0016616 | oxidoreductase activity, acting on the CH-OH group of donors, NAD or NA | molecular function | 9.62E-06 |
| GO:0022613 | ribonucleoprotein complex biogenesis | biological process | 1.98E-05 |
| GO:0016874 | ligase activity | molecular function | 2.26E-05 |
| GO:0062129 | chitin-based extracellular matrix | cellular component | 2.34E-05 |
| GO:0016765 | transferase activity, transferring alkyl or aryl (other than methyl) groups | molecular function | 2.36E-05 |
| GO:0009150 | purine ribonucleotide metabolic process | biological process | 3.06E-05 |
| GO:0033108 | mitochondrial respiratory chain complex assembly | biological process | 3.48E-05 |
| GO:0004180 | carboxypeptidase activity | molecular function | 3.99E-05 |
| GO:0009259 | ribonucleotide metabolic process | biological process | 5.73E-05 |

| | | | | |
|---|---|---|---|---|
| GO:0031012 | extracellular matrix | cellular component | 6.74E-05 | |
| GO:0072521 | purine-containing compound metabolic process | biological process | 6.77E-05 | |
| GO:0019842 | vitamin binding | molecular function | | 7.03E-05 |
| GO:0009152 | purine ribonucleotide biosynthetic process | biological process | 7.13E-05 | |
| GO:0019693 | ribose phosphate metabolic process | biological process | 8.57E-05 | |
| GO:0006163 | purine nucleotide metabolic process | biological process | 8.58E-05 | |
| GO:0006091 | generation of precursor metabolites and energy | biological process | 8.99E-05 | |
| GO:0016614 | oxidoreductase activity, acting on CH-OH group of donors | molecular function | 9.85E-05 | |
| GO:0019395 | fatty acid oxidation | biological process | 0.0001046221833 | |
| GO:0044272 | sulfur compound biosynthetic process | biological process | 0.0001171261428 | |
| GO:0042273 | ribosomal large subunit biogenesis | biological process | 0.0001386986191 | |
| GO:0034440 | lipid oxidation | biological process | 0.0001452860779 | |
| GO:0006164 | purine nucleotide biosynthetic process | biological process | 0.0001494080021 | |
| GO:0044085 | cellular component biogenesis | biological process | 0.0001856725122 | |
| GO:0055086 | nucleobase-containing small molecule metabolic process | biological process | 0.000199043776 | |
| GO:0019829 | ATPase-coupled cation transmembrane transporter activity | molecular function | 0.0002180134635 | |
| GO:0005214 | structural constituent of chitin-based cuticle | molecular function | 0.0002253155765 | |
| GO:0017171 | serine hydrolase activity | molecular function | 0.0002556510454 | |
| GO:0072329 | monocarboxylic acid catabolic process | biological process | 0.0002564189099 | |
| GO:0043228 | non-membrane-bounded organelle | cellular component | | 0.000261781106 |
| GO:0043232 | intracellular non-membrane-bounded organelle | cellular component | | 0.000261781106 |
| GO:0006753 | nucleoside phosphate metabolic process | biological process | 0.0002619868391 | |
| GO:0000221 | vacuolar proton-transporting V-type ATPase, V1 domain | cellular component | 0.0002785858967 | |
| GO:0010257 | NADH dehydrogenase complex assembly | biological process | 0.0003298477803 | |
| GO:0032981 | mitochondrial respiratory chain complex I assembly | biological process | 0.0003298477803 | |
| GO:0008033 | tRNA processing | biological process | 0.0003825697999 | |
| GO:0072522 | purine-containing compound biosynthetic process | biological process | 0.000387143632 | |
| GO:0006551 | leucine metabolic process | biological process | 0.0004007967166 | |
| GO:0006573 | valine metabolic process | biological process | 0.0004007967166 | |
| GO:0070071 | proton-transporting two-sector ATPase complex assembly | biological process | 0.0004013200695 | |
| GO:0016471 | vacuolar proton-transporting V-type ATPase complex | cellular component | 0.0004091865977 | |
| GO:0008150 | biological_process | biological process | | 0.0005059512276 |
| GO:0008233 | peptidase activity | molecular function | 0.0005431395459 | |

| | | | | |
|---|---|---|---|---|
| GO:0004364 | glutathione transferase activity | molecular function | 0.0005743363221 | |
| GO:0042625 | ATPase-coupled ion transmembrane transporter activity | molecular function | 0.0005743363221 | |
| GO:0009117 | nucleotide metabolic process | biological process | 0.0005815834619 | |
| GO:0033177 | proton-transporting two-sector ATPase complex, proton-transporting domai | cellular component | 0.0005890047867 | |
| GO:0016861 | intramolecular oxidoreductase activity, interconverting aldoses and ketoses | molecular function | 0.000598709471 | |
| GO:0009062 | fatty acid catabolic process | biological process | 0.0006971776712 | |
| GO:0006635 | fatty acid beta-oxidation | biological process | 0.0007406368421 | |
| GO:0008236 | serine-type peptidase activity | molecular function | 0.0009053898173 | |
| GO:0022890 | inorganic cation transmembrane transporter activity | molecular function | | 0.001111657327 |
| GO:0006030 | chitin metabolic process | biological process | | 0.001141836046 |
| GO:0006022 | aminoglycan metabolic process | biological process | | 0.001247902434 |
| GO:0016810 | hydrolase activity, acting on carbon-nitrogen (but not peptide) bonds | molecular function | 0.001338585225 | |
| GO:0006040 | amino sugar metabolic process | biological process | | 0.001428513299 |
| GO:1901071 | glucosamine-containing compound metabolic process | biological process | | 0.001428513299 |
| GO:0016053 | organic acid biosynthetic process | biological process | 0.001432578695 | |
| GO:0046394 | carboxylic acid biosynthetic process | biological process | 0.001432578695 | |
| GO:0006839 | mitochondrial transport | biological process | 0.001649529561 | |
| GO:0008970 | phospholipase A1 activity | molecular function | 0.001913812403 | |
| GO:0016811 | hydrolase activity, acting on carbon-nitrogen (but not peptide) bonds, in lin | molecular function | 0.002019041973 | |
| GO:0005777 | peroxisome | cellular component | 0.002154455465 | |
| GO:0008010 | structural constituent of chitin-based larval cuticle | molecular function | 0.002252399852 | |
| GO:0008324 | cation transmembrane transporter activity | molecular function | | 0.002476469495 |
| GO:0040003 | chitin-based cuticle development | biological process | 0.002541176052 | |
| GO:0042579 | microbody | cellular component | 0.00281720241 | |
| GO:0006552 | leucine catabolic process | biological process | 0.00295625856 | |
| GO:0006574 | valine catabolic process | biological process | 0.00295625856 | |
| GO:0009987 | cellular process | biological process | | 0.002957370882 |
| GO:0004252 | serine-type endopeptidase activity | molecular function | 0.003346737478 | |
| GO:0042335 | cuticle development | biological process | 0.003467218044 | |
| GO:0000062 | fatty-acyl-CoA binding | molecular function | 0.003781994656 | |
| GO:1901567 | fatty acid derivative binding | molecular function | 0.003781994656 | |
| GO:0045851 | pH reduction | biological process | 0.003864591794 | |
| GO:0051452 | intracellular pH reduction | biological process | 0.003864591794 | |

| | | | | |
|---|---|---|---|---|
| GO:0009165 | nucleotide biosynthetic process | biological process | 0.003912290758 | |
| GO:1901293 | nucleoside phosphate biosynthetic process | biological process | 0.004128769852 | |
| GO:0030170 | pyridoxal phosphate binding | molecular function | | 0.004232591179 |
| GO:0070279 | vitamin B6 binding | molecular function | | 0.004232591179 |
| GO:0004175 | endopeptidase activity | molecular function | 0.004432912034 | |
| GO:0004181 | metallocarboxypeptidase activity | molecular function | 0.004485445737 | |
| GO:0006400 | tRNA modification | biological process | 0.004909433931 | |
| GO:0009068 | aspartate family amino acid catabolic process | biological process | 0.005162601373 | |
| GO:0140101 | catalytic activity, acting on a tRNA | molecular function | 0.005452433944 | |
| GO:0000462 | maturation of SSU-rRNA from tricistronic rRNA transcript (SSU-rRNA, 5. | biological process | 0.005454184733 | |
| GO:0008097 | 5S rRNA binding | molecular function | 0.005463126611 | |
| GO:0016787 | hydrolase activity | molecular function | 0.00570035844 | |
| GO:0006743 | ubiquinone metabolic process | biological process | 0.005871119209 | |
| GO:0006744 | ubiquinone biosynthetic process | biological process | 0.005871119209 | |
| GO:1901661 | quinone metabolic process | biological process | 0.005871119209 | |
| GO:1901663 | quinone biosynthetic process | biological process | 0.005871119209 | |
| GO:0044267 | cellular protein metabolic process | biological process | 0.006036567387 | |
| GO:0008238 | exopeptidase activity | molecular function | 0.006306461199 | |
| GO:0120227 | acyl-CoA binding | molecular function | 0.006306461199 | |
| GO:1901137 | carbohydrate derivative biosynthetic process | biological process | 0.006477464704 | |
| GO:0070013 | intracellular organelle lumen | cellular component | 0.007006197721 | |
| GO:0008237 | metallopeptidase activity | molecular function | 0.007040024529 | |
| GO:0051453 | regulation of intracellular pH | biological process | 0.007597684389 | |
| GO:0070585 | protein localization to mitochondrion | biological process | 0.007755423853 | |
| GO:0072655 | establishment of protein localization to mitochondrion | biological process | 0.007755423853 | |
| GO:0031974 | membrane-enclosed lumen | cellular component | 0.007847114258 | |
| GO:0043233 | organelle lumen | cellular component | 0.007847114258 | |
| GO:0043170 | macromolecule metabolic process | biological process | 0.008065305058 | |
| GO:0033293 | monocarboxylic acid binding | molecular function | 0.008294015515 | |
| GO:0015318 | inorganic molecular entity transmembrane transporter activity | molecular function | | 0.008307471494 |
| GO:0005783 | endoplasmic reticulum | cellular component | | 0.008339105447 |
| GO:0008483 | transaminase activity | molecular function | | 0.008346280963 |
| GO:0045039 | protein insertion into mitochondrial inner membrane | biological process | 0.008413266061 | |

| | | | | |
|---|---|---|---|---|
| GO:0006885 | regulation of pH | biological process | 0.008507783905 | |
| GO:0030641 | regulation of cellular pH | biological process | 0.008507783905 | |
| GO:0044255 | cellular lipid metabolic process | biological process | 0.008507783905 | |
| GO:0008235 | metalloexopeptidase activity | molecular function | 0.009118843356 | |
| GO:0051204 | protein insertion into mitochondrial membrane | biological process | 0.009284119557 | |
| GO:0006414 | translational elongation | biological process | 0.01048456506 | |
| GO:0000313 | organellar ribosome | cellular component | 0.01057661459 | |
| GO:0005761 | mitochondrial ribosome | cellular component | 0.01057661459 | |
| GO:0006591 | ornithine metabolic process | biological process | 0.01087918789 | |
| GO:0006750 | glutathione biosynthetic process | biological process | 0.01087918789 | |
| GO:0019184 | nonribosomal peptide biosynthetic process | biological process | 0.01087918789 | |
| GO:0070070 | proton-transporting V-type ATPase complex assembly | biological process | 0.01091329543 | |
| GO:0070072 | vacuolar proton-transporting V-type ATPase complex assembly | biological process | 0.01091329543 | |
| GO:0008535 | respiratory chain complex IV assembly | biological process | 0.01238468104 | |
| GO:0042398 | cellular modified amino acid biosynthetic process | biological process | 0.01238468104 | |
| GO:0005736 | RNA polymerase I complex | cellular component | 0.0137727307 | |
| GO:0035384 | thioester biosynthetic process | biological process | 0.01403994531 | |
| GO:0071616 | acyl-CoA biosynthetic process | biological process | 0.01403994531 | |
| GO:0004656 | procollagen-proline 4-dioxygenase activity | molecular function | | 0.01464037518 |
| GO:0009055 | electron transfer activity | molecular function | | 0.01464037518 |
| GO:0015075 | ion transmembrane transporter activity | molecular function | | 0.01464037518 |
| GO:0019798 | procollagen-proline dioxygenase activity | molecular function | | 0.01464037518 |
| GO:0031406 | carboxylic acid binding | molecular function | | 0.01464037518 |
| GO:0031543 | peptidyl-proline dioxygenase activity | molecular function | | 0.01464037518 |
| GO:0031545 | peptidyl-proline 4-dioxygenase activity | molecular function | | 0.01464037518 |
| GO:0043177 | organic acid binding | molecular function | | 0.01464037518 |
| GO:0005759 | mitochondrial matrix | cellular component | 0.01618402047 | |
| GO:0030684 | preribosome | cellular component | 0.01633881831 | |
| GO:0044242 | cellular lipid catabolic process | biological process | 0.01642262549 | |
| GO:0030490 | maturation of SSU-rRNA | biological process | 0.01810870988 | |
| GO:0005587 | collagen type IV trimer | cellular component | 0.01853272113 | |
| GO:0098642 | network-forming collagen trimer | cellular component | 0.01853272113 | |
| GO:0098651 | basement membrane collagen trimer | cellular component | 0.01853272113 | |

| | | | | |
|---|---|---|---|---|
| GO:0000177 | cytoplasmic exosome (RNase complex) | cellular component | 0.01864468139 | |
| GO:0000220 | vacuolar proton-transporting V-type ATPase, V0 domain | cellular component | 0.01864468139 | |
| GO:0003012 | muscle system process | biological process | 0.01875134577 | |
| GO:0004185 | serine-type carboxypeptidase activity | molecular function | 0.01912242894 | |
| GO:0016624 | oxidoreductase activity, acting on the aldehyde or oxo group of donors, disu | molecular function | 0.01912242894 | |
| GO:0036042 | long-chain fatty acyl-CoA binding | molecular function | 0.01912242894 | |
| GO:0070181 | small ribosomal subunit rRNA binding | molecular function | 0.01912242894 | |
| GO:0016042 | lipid catabolic process | biological process | 0.01930424834 | |
| GO:0016407 | acetyltransferase activity | molecular function | | 0.0193687732 |
| GO:0000178 | exosome (RNase complex) | cellular component | 0.01983238558 | |
| GO:0071722 | detoxification of arsenic-containing substance | biological process | 0.01989735232 | |
| GO:0006094 | gluconeogenesis | biological process | 0.0203378684 | |
| GO:0019319 | hexose biosynthetic process | biological process | 0.0203378684 | |
| GO:0008652 | cellular amino acid biosynthetic process | biological process | 0.0205590804 | |
| GO:0006629 | lipid metabolic process | biological process | 0.02171804837 | |
| GO:0007035 | vacuolar acidification | biological process | 0.02217951741 | |
| GO:0045240 | dihydrolipoyl dehydrogenase complex | cellular component | 0.02279992911 | |
| GO:0003723 | RNA binding | molecular function | 0.02296568616 | |
| GO:0009066 | aspartate family amino acid metabolic process | biological process | 0.02324629639 | |
| GO:0005506 | iron ion binding | molecular function | | 0.02343903549 |
| GO:0033866 | nucleoside bisphosphate biosynthetic process | biological process | 0.02352238892 | |
| GO:0034030 | ribonucleoside bisphosphate biosynthetic process | biological process | 0.02352238892 | |
| GO:0034033 | purine nucleoside bisphosphate biosynthetic process | biological process | 0.02352238892 | |
| GO:0009451 | RNA modification | biological process | 0.02365637099 | |
| GO:0000470 | maturation of LSU-rRNA | biological process | 0.02385845099 | |
| GO:0017004 | cytochrome complex assembly | biological process | 0.02385845099 | |
| GO:0005730 | nucleolus | cellular component | 0.02467463525 | |
| GO:0033179 | proton-transporting V-type ATPase, V0 domain | cellular component | 0.02480265 | |
| GO:0002814 | negative regulation of biosynthetic process of antibacterial peptides active a | biological process | 0.0255810848 | |
| GO:0022900 | electron transport chain | biological process | 0.0262425427 | |
| GO:0019205 | nucleobase-containing compound kinase activity | molecular function | | 0.02648357792 |
| GO:1901607 | alpha-amino acid biosynthetic process | biological process | 0.02666350585 | |
| GO:0016705 | oxidoreductase activity, acting on paired donors, with incorporation or redu | molecular function | | 0.0280205798 |

| | | | | |
|---|---|---|---|---|
| GO:0016747 | transferase activity, transferring acyl groups other than amino-acyl groups | molecular function | | 0.0280205798 |
| GO:0045239 | tricarboxylic acid cycle enzyme complex | cellular component | 0.02827301997 | |
| GO:1905354 | exoribonuclease complex | cellular component | 0.02827301997 | |
| GO:0000176 | nuclear exosome (RNase complex) | cellular component | 0.02843422115 | |
| GO:0005753 | mitochondrial proton-transporting ATP synthase complex | cellular component | 0.02843422115 | |
| GO:0045259 | proton-transporting ATP synthase complex | cellular component | 0.02843422115 | |
| GO:0008080 | N-acetyltransferase activity | molecular function | | 0.02852329209 |
| GO:0007006 | mitochondrial membrane organization | biological process | 0.02938717754 | |
| GO:0007007 | inner mitochondrial membrane organization | biological process | 0.03037009605 | |
| GO:0004576 | oligosaccharyl transferase activity | molecular function | | 0.03105609791 |
| GO:0016776 | phosphotransferase activity, phosphate group as acceptor | molecular function | | 0.03105609791 |
| GO:0016627 | oxidoreductase activity, acting on the CH-CH group of donors | molecular function | 0.03203123429 | |
| GO:0016706 | 2-oxoglutarate-dependent dioxygenase activity | molecular function | | 0.03225799337 |
| GO:1990542 | mitochondrial transmembrane transport | biological process | 0.03256368185 | |
| GO:0004090 | carbonyl reductase (NADPH) activity | molecular function | 0.03522793176 | |
| GO:0005504 | fatty acid binding | molecular function | 0.035555088 | |
| GO:0006936 | muscle contraction | biological process | 0.03726772367 | |
| GO:0006081 | cellular aldehyde metabolic process | biological process | 0.03760516377 | |
| GO:0016410 | N-acyltransferase activity | molecular function | | 0.03778679892 |
| GO:0072593 | reactive oxygen species metabolic process | biological process | 0.03866436421 | |
| GO:0004035 | alkaline phosphatase activity | molecular function | 0.0386851109 | |
| GO:0006633 | fatty acid biosynthetic process | biological process | 0.03919230145 | |
| GO:0016746 | transferase activity, transferring acyl groups | molecular function | | 0.04248438207 |
| GO:0070069 | cytochrome complex | cellular component | 0.04316291569 | |
| GO:0000276 | mitochondrial proton-transporting ATP synthase complex, coupling factor F | cellular component | 0.04353596999 | |
| GO:0045263 | proton-transporting ATP synthase complex, coupling factor F(o) | cellular component | 0.04353596999 | |
| GO:0009069 | serine family amino acid metabolic process | biological process | 0.04531735226 | |
| GO:0004046 | aminoacylase activity | molecular function | 0.04595440937 | |
| GO:0070008 | serine-type exopeptidase activity | molecular function | 0.04595440937 | |
| GO:0055067 | monovalent inorganic cation homeostasis | biological process | 0.04632853321 | |
| GO:0016903 | oxidoreductase activity, acting on the aldehyde or oxo group of donors | molecular function | 0.04696247282 | |
| GO:0002787 | negative regulation of antibacterial peptide production | biological process | 0.04762779815 | |
| GO:0002806 | negative regulation of antimicrobial peptide biosynthetic process | biological process | 0.04762779815 | |

| | | | | |
|---|---|---|---|---|
| GO:0002809 | negative regulation of antibacterial peptide biosynthetic process | biological process | 0.04762779815 | |
| GO:0034475 | U4 snRNA 3'-end processing | biological process | 0.04762779815 | |
| GO:0000275 | mitochondrial proton-transporting ATP synthase complex, catalytic sector F | cellular component | 0.04800918052 | |
| GO:0005581 | collagen trimer | cellular component | 0.04800918052 | |
| GO:0005756 | mitochondrial proton-transporting ATP synthase, central stalk | cellular component | 0.04800918052 | |
| GO:0005903 | brush border | cellular component | 0.04800918052 | |
| GO:0031314 | extrinsic component of mitochondrial inner membrane | cellular component | 0.04800918052 | |
| GO:0045269 | proton-transporting ATP synthase, central stalk | cellular component | 0.04800918052 | |
| GO:0098862 | cluster of actin-based cell projections | cellular component | 0.04800918052 | |
| GO:0005622 | intracellular | cellular component | | 0.04878956165 |
| GO:1901292 | nucleoside phosphate catabolic process | biological process | 0.04944957929 | |

| GO | name | GO type | p-value FDR *D. melanogaster* | p-value FDR *D. virilis* |
|---|---|---|---|---|
| GO:0042302 | structural constituent of cuticle | molecular function | 1.05E-06 | 1.05E-04 |
| GO:0042335 | cuticle development | biological process | 3.10E-09 | |
| GO:0031012 | extracellular matrix | cellular component | 2.15E-08 | |
| GO:0040003 | chitin-based cuticle development | biological process | 6.58E-08 | |
| GO:0062129 | chitin-based extracellular matrix | cellular component | 6.84E-07 | |
| GO:0005214 | structural constituent of chitin-based cuticle | molecular function | 1.05E-06 | |
| GO:1901071 | glucosamine-containing compound metabolic process | biological process | 3.61E-04 | |
| GO:0006040 | amino sugar metabolic process | biological process | 5.01E-04 | |
| GO:0008010 | structural constituent of chitin-based larval cuticle | molecular function | 1.23E-03 | |
| GO:0004099 | chitin deacetylase activity | molecular function | 1.39E-03 | |
| GO:0048856 | anatomical structure development | biological process | 7.39E-03 | |
| GO:0005198 | structural molecule activity | molecular function | 1.10E-02 | |
| GO:0019213 | deacetylase activity | molecular function | 1.85E-02 | |
| GO:0004252 | serine-type endopeptidase activity | molecular function | | 2.53E-02 |
| GO:0008236 | serine-type peptidase activity | molecular function | | 3.58E-02 |
| GO:0017171 | serine hydrolase activity | molecular function | | 3.58E-02 |
| GO:0005887 | integral component of plasma membrane | cellular component | 4.69E-02 | |

| GO | name | GO type | p-value FDR *D. melanogaster* | p-value FDR *D. virilis* |
|---|---|---|---|---|
| GO:0005506 | iron ion binding | molecular function | | 6.57E-06 |
| GO:0016705 | oxidoreductase activity, acting on paired donors, with incorporation or redu | molecular function | | 6.57E-06 |
| GO:0020037 | heme binding | molecular function | | 6.57E-06 |
| GO:0046906 | tetrapyrrole binding | molecular function | | 6.57E-06 |
| GO:0016491 | oxidoreductase activity | molecular function | | 6.73E-04 |
| GO:0055114 | oxidation-reduction process | biological process | | 1.04E-03 |
| GO:0035080 | heat shock-mediated polytene chromosome puffing | biological process | 2.68E-02 | |
| GO:0035079 | polytene chromosome puffing | biological process | 3.48E-02 | |

ST5. Late pupae-specific enriched GO terms

| GO | name | GO type | p-value FDR *D. melanogaster* | p-value FDR *D. virilis* |
|---|---|---|---|---|
| GO:0042302 | structural constituent of cuticle | molecular function | | 1.16E-09 |
| GO:0031012 | extracellular matrix | cellular component | 4.36E-04 | |
| GO:0005198 | structural molecule activity | molecular function | | 3.07E-03 |
| GO:0005887 | integral component of plasma membrane | cellular component | 1.19E-02 | |
| GO:0008061 | chitin binding | molecular function | | 1.48E-02 |
| GO:0062129 | chitin-based extracellular matrix | cellular component | 2.11E-02 | |
| GO:0031226 | intrinsic component of plasma membrane | cellular component | 2.55E-02 | |

| GO | name | GO type | p-value FDR *D. melanogaster* | p-value FDR *D. virilis* |
|---|---|---|---|---|
| GO:0031966 | mitochondrial membrane | cellular component | 5.49E-03 | 6.19E-04 |
| GO:0006090 | pyruvate metabolic process | biological process | 2.13E-02 | 7.94E-04 |
| GO:0006165 | nucleoside diphosphate phosphorylation | biological process | 9.44E-03 | 2.28E-03 |
| GO:0006096 | glycolytic process | biological process | 4.28E-03 | 6.58E-03 |
| GO:0006757 | ATP generation from ADP | biological process | 4.28E-03 | 6.58E-03 |
| GO:0046939 | nucleotide phosphorylation | biological process | 1.32E-02 | 2.28E-03 |
| GO:0007601 | visual perception | biological process | 2.20E-02 | 2.28E-03 |
| GO:0009135 | purine nucleoside diphosphate metabolic process | biological process | 9.79E-03 | 1.01E-02 |
| GO:0009179 | purine ribonucleoside diphosphate metabolic process | biological process | 9.79E-03 | 1.01E-02 |
| GO:0046031 | ADP metabolic process | biological process | 9.79E-03 | 1.01E-02 |
| GO:0016052 | carbohydrate catabolic process | biological process | 1.28E-02 | 0.01013766833 |
| GO:0009185 | ribonucleoside diphosphate metabolic process | biological process | 2.01E-02 | 1.01E-02 |
| GO:0004743 | pyruvate kinase activity | molecular function | 3.00E-02 | 1.62E-02 |
| GO:0030955 | potassium ion binding | molecular function | 3.00E-02 | 1.62E-02 |
| GO:0031420 | alkali metal ion binding | molecular function | 3.00E-02 | 1.62E-02 |
| GO:0032504 | multicellular organism reproduction | biological process | 2.95E-52 | |
| GO:0000003 | reproduction | biological process | 3.23E-51 | |
| GO:0005615 | extracellular space | cellular component | 1.09E-15 | |
| GO:0003341 | cilium movement | biological process | 1.11E-09 | |
| GO:0031514 | motile cilium | cellular component | 2.47E-09 | |
| GO:0036126 | sperm flagellum | cellular component | 8.61E-08 | |
| GO:0097729 | 9+2 motile cilium | cellular component | 8.61E-08 | |
| GO:0030286 | dynein complex | cellular component | 2.85E-07 | |
| GO:0060294 | cilium movement involved in cell motility | biological process | 9.51E-07 | |
| GO:0048029 | monosaccharide binding | molecular function | 5.32E-06 | |
| GO:0019236 | response to pheromone | biological process | 6.05E-06 | |
| GO:0005929 | cilium | cellular component | 6.22E-06 | |
| GO:0005858 | axonemal dynein complex | cellular component | 6.72E-06 | |
| GO:0001539 | cilium or flagellum-dependent cell motility | biological process | 7.93E-06 | |
| GO:0060285 | cilium-dependent cell motility | biological process | 7.93E-06 | |
| GO:0046008 | regulation of female receptivity, post-mating | biological process | 2.60E-05 | |
| GO:0007602 | phototransduction | biological process | | 3.15E-05 |

| | | | | |
|---|---|---|---|---|
| GO:0009581 | detection of external stimulus | biological process | | 3.15E-05 |
| GO:0009582 | detection of abiotic stimulus | biological process | | 3.15E-05 |
| GO:0009583 | detection of light stimulus | biological process | | 3.15E-05 |
| GO:0009605 | response to external stimulus | biological process | | 3.15E-05 |
| GO:0005975 | carbohydrate metabolic process | biological process | | 3.33E-05 |
| GO:0044703 | multi-organism reproductive process | biological process | 4.24E-05 | |
| GO:0051704 | multi-organism process | biological process | 4.24E-05 | |
| GO:0045434 | negative regulation of female receptivity, post-mating | biological process | 6.28E-05 | |
| GO:0043695 | detection of pheromone | biological process | 7.61E-05 | |
| GO:0044706 | multi-multicellular organism process | biological process | 7.86E-05 | |
| GO:0046692 | sperm competition | biological process | 7.86E-05 | |
| GO:0006099 | tricarboxylic acid cycle | biological process | 8.16E-05 | |
| GO:0045924 | regulation of female receptivity | biological process | 8.16E-05 | |
| GO:0030317 | flagellated sperm motility | biological process | 1.16E-04 | |
| GO:0097722 | sperm motility | biological process | 1.16E-04 | |
| GO:0007606 | sensory perception of chemical stimulus | biological process | 1.28E-04 | |
| GO:0009628 | response to abiotic stimulus | biological process | | 1.97E-04 |
| GO:0009314 | response to radiation | biological process | | 2.13E-04 |
| GO:0009416 | response to light stimulus | biological process | | 2.13E-04 |
| GO:0016616 | oxidoreductase activity, acting on the CH-OH group of donors, NAD or NA | molecular function | | 2.63E-04 |
| GO:0015556 | C4-dicarboxylate transmembrane transporter activity | molecular function | 2.67E-04 | |
| GO:0018095 | protein polyglutamylation | biological process | 3.11E-04 | |
| GO:0016614 | oxidoreductase activity, acting on CH-OH group of donors | molecular function | | 3.56E-04 |
| GO:0007621 | negative regulation of female receptivity | biological process | 3.56E-04 | |
| GO:0070286 | axonemal dynein complex assembly | biological process | 4.00E-04 | |
| GO:0007600 | sensory perception | biological process | 4.99E-04 | |
| GO:0070739 | protein-glutamic acid ligase activity | molecular function | 5.23E-04 | |
| GO:0070740 | tubulin-glutamic acid ligase activity | molecular function | 5.23E-04 | |
| GO:0055085 | transmembrane transport | biological process | 5.47E-04 | |
| GO:0098656 | anion transmembrane transport | biological process | 5.72E-04 | |
| GO:0018401 | peptidyl-proline hydroxylation to 4-hydroxy-L-proline | biological process | 6.93E-04 | |
| GO:0004656 | procollagen-proline 4-dioxygenase activity | molecular function | 9.18E-04 | |
| GO:0019798 | procollagen-proline dioxygenase activity | molecular function | 9.18E-04 | |

| | | | | |
|---|---|---|---|---|
| GO:0004867 | serine-type endopeptidase inhibitor activity | molecular function | 1.05E-03 | |
| GO:0003998 | acylphosphatase activity | molecular function | | 0.001074445715 |
| GO:0016701 | oxidoreductase activity, acting on single donors with incorporation of mole | molecular function | 0.001138151711 | |
| GO:0019511 | peptidyl-proline hydroxylation | biological process | 0.001174931459 | |
| GO:0015141 | succinate transmembrane transporter activity | molecular function | 0.001540820312 | |
| GO:0031545 | peptidyl-proline 4-dioxygenase activity | molecular function | 0.001540820312 | |
| GO:0030246 | carbohydrate binding | molecular function | 0.001756309379 | |
| GO:0050953 | sensory perception of light stimulus | biological process | | 0.002277478376 |
| GO:0060180 | female mating behavior | biological process | 0.002279700881 | |
| GO:0005310 | dicarboxylic acid transmembrane transporter activity | molecular function | 0.002358207656 | |
| GO:0031543 | peptidyl-proline dioxygenase activity | molecular function | 0.002552627972 | |
| GO:0016702 | oxidoreductase activity, acting on single donors with incorporation of mole | molecular function | 0.002595857901 | |
| GO:0045505 | dynein intermediate chain binding | molecular function | 0.002595857901 | |
| GO:0006839 | mitochondrial transport | biological process | 0.002992801262 | |
| GO:0015740 | C4-dicarboxylate transport | biological process | 0.003067219939 | |
| GO:0051606 | detection of stimulus | biological process | | 0.003197185802 |
| GO:0015291 | secondary active transmembrane transporter activity | molecular function | 0.003376042937 | |
| GO:0031418 | L-ascorbic acid binding | molecular function | 0.003519252682 | |
| GO:0045504 | dynein heavy chain binding | molecular function | 0.004059195433 | |
| GO:1990542 | mitochondrial transmembrane transport | biological process | 0.004315409788 | |
| GO:0009132 | nucleoside diphosphate metabolic process | biological process | | 0.005609414293 |
| GO:0070735 | protein-glycine ligase activity | molecular function | 0.005832763966 | |
| GO:0004457 | lactate dehydrogenase activity | molecular function | | 0.005855681456 |
| GO:0004459 | L-lactate dehydrogenase activity | molecular function | | 0.005855681456 |
| GO:0018126 | protein hydroxylation | biological process | 0.006173644484 | |
| GO:1990204 | oxidoreductase complex | cellular component | | 0.006260984624 |
| GO:0015744 | succinate transport | biological process | 0.006548399453 | |
| GO:0018094 | protein polyglycylation | biological process | 0.006548399453 | |
| GO:0005342 | organic acid transmembrane transporter activity | molecular function | 0.007534349177 | |
| GO:0046943 | carboxylic acid transmembrane transporter activity | molecular function | 0.007534349177 | |
| GO:0015370 | solute:sodium symporter activity | molecular function | 0.008286116424 | |
| GO:0005534 | galactose binding | molecular function | 0.008862267696 | |
| GO:0070585 | protein localization to mitochondrion | biological process | 0.009127739366 | |

| | | | | |
|---|---|---|---|---|
| GO:0072655 | establishment of protein localization to mitochondrion | biological process | 0.009127739366 | |
| GO:0015709 | thiosulfate transport | biological process | 0.009411630239 | |
| GO:0015729 | oxaloacetate transport | biological process | 0.009411630239 | |
| GO:0015743 | malate transport | biological process | 0.009411630239 | |
| GO:0071422 | succinate transmembrane transport | biological process | 0.009411630239 | |
| GO:0071423 | malate transmembrane transport | biological process | 0.009411630239 | |
| GO:0030150 | protein import into mitochondrial matrix | biological process | 0.009438879178 | |
| GO:0006835 | dicarboxylic acid transport | biological process | 0.01004178763 | |
| GO:0005875 | microtubule associated complex | cellular component | 0.01081981585 | |
| GO:0019200 | carbohydrate kinase activity | molecular function | 0.01092403046 | |
| GO:0018208 | peptidyl-proline modification | biological process | 0.01137570845 | |
| GO:1903825 | organic acid transmembrane transport | biological process | 0.01141927478 | |
| GO:1905039 | carboxylic acid transmembrane transport | biological process | 0.01141927478 | |
| GO:0005930 | axoneme | cellular component | 0.01159448831 | |
| GO:0036157 | outer dynein arm | cellular component | 0.01163098088 | |
| GO:0036158 | outer dynein arm assembly | biological process | 0.01262759041 | |
| GO:0071482 | cellular response to light stimulus | biological process | 0.01359238018 | |
| GO:0098800 | inner mitochondrial membrane protein complex | cellular component | 0.01367588853 | |
| GO:0016491 | oxidoreductase activity | molecular function | | 0.0162450171 |
| GO:0098573 | intrinsic component of mitochondrial membrane | cellular component | 0.0168996024 | |
| GO:0005343 | organic acid:sodium symporter activity | molecular function | 0.01708080832 | |
| GO:0005506 | iron ion binding | molecular function | 0.01727608323 | |
| GO:0015117 | thiosulfate transmembrane transporter activity | molecular function | 0.01842848004 | |
| GO:0015131 | oxaloacetate transmembrane transporter activity | molecular function | 0.01842848004 | |
| GO:0015140 | malate transmembrane transporter activity | molecular function | 0.01842848004 | |
| GO:0036156 | inner dynein arm | cellular component | 0.01874540299 | |
| GO:0015293 | symporter activity | molecular function | 0.01970413871 | |
| GO:0005326 | neurotransmitter transmembrane transporter activity | molecular function | 0.01971606196 | |
| GO:0008643 | carbohydrate transport | biological process | 0.02014684702 | |
| GO:0045333 | cellular respiration | biological process | 0.02014684702 | |
| GO:0005576 | extracellular region | cellular component | 0.02028527686 | |
| GO:0007617 | mating behavior | biological process | 0.02109159141 | |
| GO:0015849 | organic acid transport | biological process | 0.02175553457 | |

| | | | | |
|---|---|---|---|---|
| GO:0046942 | carboxylic acid transport | biological process | 0.02175553457 | |
| GO:0009060 | aerobic respiration | biological process | 0.02261434586 | |
| GO:0061135 | endopeptidase regulator activity | molecular function | 0.02311732195 | |
| GO:0006817 | phosphate ion transport | biological process | 0.02349200894 | |
| GO:0004396 | hexokinase activity | molecular function | 0.02418472997 | |
| GO:0008865 | fructokinase activity | molecular function | 0.02418472997 | |
| GO:0009055 | electron transfer activity | molecular function | 0.02418472997 | |
| GO:0015175 | neutral amino acid transmembrane transporter activity | molecular function | 0.02418472997 | |
| GO:0045503 | dynein light chain binding | molecular function | 0.02418472997 | |
| GO:0007608 | sensory perception of smell | biological process | 0.02549526356 | |
| GO:0015294 | solute:cation symporter activity | molecular function | 0.02616130439 | |
| GO:0015144 | carbohydrate transmembrane transporter activity | molecular function | 0.02715304443 | |
| GO:0022804 | active transmembrane transporter activity | molecular function | 0.02783061999 | |
| GO:0004866 | endopeptidase inhibitor activity | molecular function | 0.02998259105 | |
| GO:0019825 | oxygen binding | molecular function | 0.02998259105 | |
| GO:0035082 | axoneme assembly | biological process | 0.03032641158 | |
| GO:0061134 | peptidase regulator activity | molecular function | 0.03083962483 | |
| GO:0035303 | regulation of dephosphorylation | biological process | | 0.03088727878 |
| GO:0030414 | peptidase inhibitor activity | molecular function | 0.03224527458 | |
| GO:0005839 | proteasome core complex | cellular component | | 0.03443734758 |
| GO:0098803 | respiratory chain complex | cellular component | | 0.03443734758 |
| GO:0044743 | protein transmembrane import into intracellular organelle | biological process | 0.03478333172 | |
| GO:0032592 | integral component of mitochondrial membrane | cellular component | 0.03622541158 | |
| GO:0002027 | regulation of heart rate | biological process | 0.03654629174 | |
| GO:0006123 | mitochondrial electron transport, cytochrome c to oxygen | biological process | 0.03654629174 | |
| GO:0016059 | deactivation of rhodopsin mediated signaling | biological process | 0.03654629174 | |
| GO:0019646 | aerobic electron transport chain | biological process | 0.03654629174 | |
| GO:0035435 | phosphate ion transmembrane transport | biological process | 0.03654629174 | |
| GO:0006102 | isocitrate metabolic process | biological process | 0.0406032645 | |
| GO:0022857 | transmembrane transporter activity | molecular function | 0.04074344244 | |
| GO:0005283 | amino acid:sodium symporter activity | molecular function | 0.04241459612 | |
| GO:0008569 | ATP-dependent microtubule motor activity, minus-end-directed | molecular function | 0.04241459612 | |
| GO:0004298 | threonine-type endopeptidase activity | molecular function | | 0.04308418597 |

| | | | | |
|---|---|---|---|---|
| GO:0070003 | threonine-type peptidase activity | molecular function | | 0.04308418597 |
| GO:0030145 | manganese ion binding | molecular function | 0.04334717208 | |
| GO:0044782 | cilium organization | biological process | 0.04391098746 | |
| GO:0005744 | TIM23 mitochondrial import inner membrane translocase complex | cellular component | 0.04454098939 | |
| GO:0051959 | dynein light intermediate chain binding | molecular function | 0.04488570713 | |
| GO:0043648 | dicarboxylic acid metabolic process | biological process | 0.04613827247 | |
| GO:0019098 | reproductive behavior | biological process | 0.04902730586 | |
